# Cerebellar stimulations prevent Levodopa-induced dyskinesia in mice and normalize brain activity

**DOI:** 10.1101/2021.09.17.460625

**Authors:** Bérénice Coutant, Jimena Laura Frontera, Elodie Perrin, Adèle Combes, Thibault Tarpin, Fabien Menardy, Caroline Mailhes-Hamon, Sylvie Perez, Bertrand Degos, Laurent Venance, Clément Léna, Daniela Popa

## Abstract

Chronic Levodopa therapy, the gold-standard treatment of Parkinson’s Disease (PD), leads to the emergence of involuntary movements, called levodopa-induced dyskinesia (LID). Cerebellar stimulations have been shown to decrease LID severity in PD patients. Here, in order to determine how cerebellar stimulations induce LID alleviation, we performed daily short trains of optogenetic stimulations of Purkinje cells (PC) in freely moving mice. We demonstrated that these stimulations are sufficient to suppress LID or even prevent their development. This symptomatic relief is accompanied by the normalization of aberrant neuronal discharge in the cerebellar nuclei, the motor cortex and the parafascicular thalamus. Inhibition of the cerebello-parafascicular pathway counteracted the beneficial effect of cerebellar stimulations. Moreover, cerebellar stimulations reversed plasticity in D1 striatal neurons and normalized the overexpression of FosB, a transcription factor causally linked to LID. These findings demonstrate LID alleviation and prevention by daily PC stimulations, which restore the function of a wide brain motor network, and may be valuable for LID treatment.

## INTRODUCTION

Motor symptoms of Parkinson’s disease (PD) are caused by a progressive loss of dopaminergic neurons in the substantia nigra *pars compacta*, and of their dense projections to the striatum. The gold-standard symptomatic therapy for PD patients is Levodopa (L-DOPA). However, with disease progression and chronic exposure to L-DOPA, 50-80% of patients experience a range of motor levodopa-induced complications within 5 years of treatment ^1^ including debilitating abnormal involuntary movements, called levodopa-induced dyskinesia (LID) ^2^. So far, very few therapeutic options are available to circumvent the advent of LID in the course of L-DOPA treatment. A better understanding of the brain networks controlling LID generation and expression is critical to the development of appropriate treatments.

LID-associated abnormalities have been consistently observed in the basal ganglia, the thalamus and the motor cortex in humans ^3,4,5,6^, primates ^7,8,9,10^ and rodents ^11,12,13,14,15^. In line with these observations, interactions in this inter-connected motor network contribute to LID pathopysiology ^16^. More recently, alleviation of LID in humans have been observed following stimulations of the cerebellum ^5, 17,18,19,20^. While a single short (1-2 minutes) session of repetitive transcranial magnetic (rTMS) continuous theta burst (cTBS) stimulation over the cerebellum only transiently reduced LID, the repetition of stimulation sessions over 2 weeks yielded a reduction of peak-dose LID over weeks after the sessions ^18, 19^. This showed that cerebellar stimulations could reduce the expression of LID. The impact of these stimulations was observed in the cerebellar nuclei ^17^, suggesting that their effect is mediated by the output cells of the cerebellar cortex, the Purkinje cells (PC) and propagated to downstream structures.

A first possibility is that cerebellar stimulations correct motor cortex dysfunction observed in dyskinesia. Indeed, dyskinetic patients present an increase in cerebral blood flow in the primary motor cortex ^21^, as well as abnormal synaptic plasticity ^22^. Similarly, dyskinetic rats exhibit changes in gene expression ^23^ and an increased activity in about half of the neurons of the motor cortex ^24^. In addition, subthalamic deep brain stimulation, which reduce PD symptoms and thus prevent the need of high L-DOPA dosage producing LID, have been proposed to act via an effect on the motor cortex ^25, 26^. Likewise, cerebellar cTBS has been shown to exert a control on motor cortex plasticity ^27^. Moreover, anodal direct current stimulation over the cerebellum, which is thought to increase the cerebello-cortical coupling ^28^, also led to a decrease in LID ^29^. Therefore, the motor cortex could be the relay of cerebellar stimulations in the treatment of LID.

LID is also directly linked to abnormal molecular events taking place in striatal neurons ^30, 31^. Most notably, LID has been causally linked to changes in the expression of FosB, a transcription factor, and its truncated splice variant ΔFosB. Dyskinetic patients ^32^, primates ^33, 34^ and rodents ^35,36,37,38,39^ show an overexpression of FosB/ΔFosB that strongly correlates with the severity of dyskinesia^38^. The upregulation of FosB/ΔFosB in striatal neurons of experimental animals is sufficient to trigger LID in response to acute administration of levodopa ^7, 40^, and reciprocally the inactivation of striatal FosB/ΔFosB reduces LID ^34, 41^ establishing the causal contribution of this transcription factor to LID. LID is associated with strong changes in striatal synaptic plasticity ^14, 42^. These aberrant corticostriatal plasticity’s are indeed a feature shared with a number of other hyperkinetic movement disorders, suggesting that they participate to the pathological state ^43, 44^. Besides its cortical inputs, the striatum receives massive inputs from the thalamus ^45^. The thalamo-striatal pathway could also relay therapeutic activities as demonstrated by the reduction of LID following deep brain stimulation of the intralaminar thalamo-striatal CM-PF complex in PD patients ^46^ and dyskinetic rats ^47^. The cerebellum indeed projects to the basal ganglia by way of the intralaminar thalamus ^48,49,50,51^ and may control the cortico-striatal plasticity ^49^. Cerebellar stimulations could therefore directly restore striatal function in LID.

To investigate the mechanisms underlying the alleviation of LID by cerebellar stimulations, we studied the effect of optogenetic PC stimulations on these abnormal involuntary movements using L7-ChR2-YFP mice ^52^ in combination with a well-known mouse model of LID ^53^. We performed daily brief sessions of theta-rhythm optogenetic stimulations of PC in Crus II, the region associated with orolingual sensorimotor function of the cerebellum ^54, 55^. These stimulations did specifically suppress, or even prevent, if administered early enough, severe orolingual LID. These behavioral findings were paralleled with a normalization of the aberrant neuronal activity in the deep cerebellar nuclei, especially the interposed nucleus, in the oral primary motor cortex and in the parafascicular thalamus, indicating a wide-scale action of cerebellar stimulations on the motor system. The chemogenetic inactivation of the cerebello-parafascicular pathway counteracted the beneficial effects of cerebellar stimulations, suggesting that they are mediated via the cerebello-thalamo-striatal pathway. Indeed, cerebellar stimulations reversed the sign of corticostriatal plasticity by promoting long-term depression in D1-expressing neurons and normalized the striatal expression of FosB/ΔFosB indicating that cerebellar stimulations act on the core of LID genesis.

## RESULTS

### Optogenetic Purkinje cell stimulations in the orolingual region of the cerebellar hemisphere specifically suppress or prevent orolingual dyskinesia

To study the effect of repeated sessions of optogenetic stimulations of PC on dyskinesia, we used a classical mouse model of LID. LID were produced by repeated systemic injections of levodopa in mice that underwent dopaminergic depletion following 6-OHDA injection in the median forebrain bundle, which project mainly to the dorsal striatum (Figure 1a-c). 6-OHDA-lesioned animals chronically treated with levodopa alone (condition “LID”, N=19) indeed exhibited severe oral, axial and limb dyskinesia, compared to non-lesioned levodopa-treated sham mice (condition “SHAM”, N=17) (Figure 1d-g). The dyskinesia score peaked around 30-40 minutes after levodopa injection (Figures S1b, S2b, S3b) as described in previous studies ^56, 57^, consistent with LID severity following plasmatic levels of levodopa ^53, 58^, hence referred to as peak-dose dyskinesia. These effects were observed during the 6 weeks of daily levodopa administration. Sham-lesioned mice exposed to chronic levodopa treatment received either corrective or preventive optogenetic cerebellar stimulations, or none. They did not exhibit any kind of severe dyskinesia, neither before nor during cerebellar stimulations (Figure S4), and were therefore pooled together for behavioral analysis. To examine whether PC stimulations efficiently reversed or prevented LID, brief trains of optogenetic stimulations at theta frequency ^19, 59^ were delivered daily in mice expressing ChR2 specifically in PC ^52, 60^ (Figure 2b,c). Stimulations either started 2 weeks after (“corrective stimulations”), or preceded (“preventive stimulations”), LID onset (Figure 1a).

**Fig. 1.**
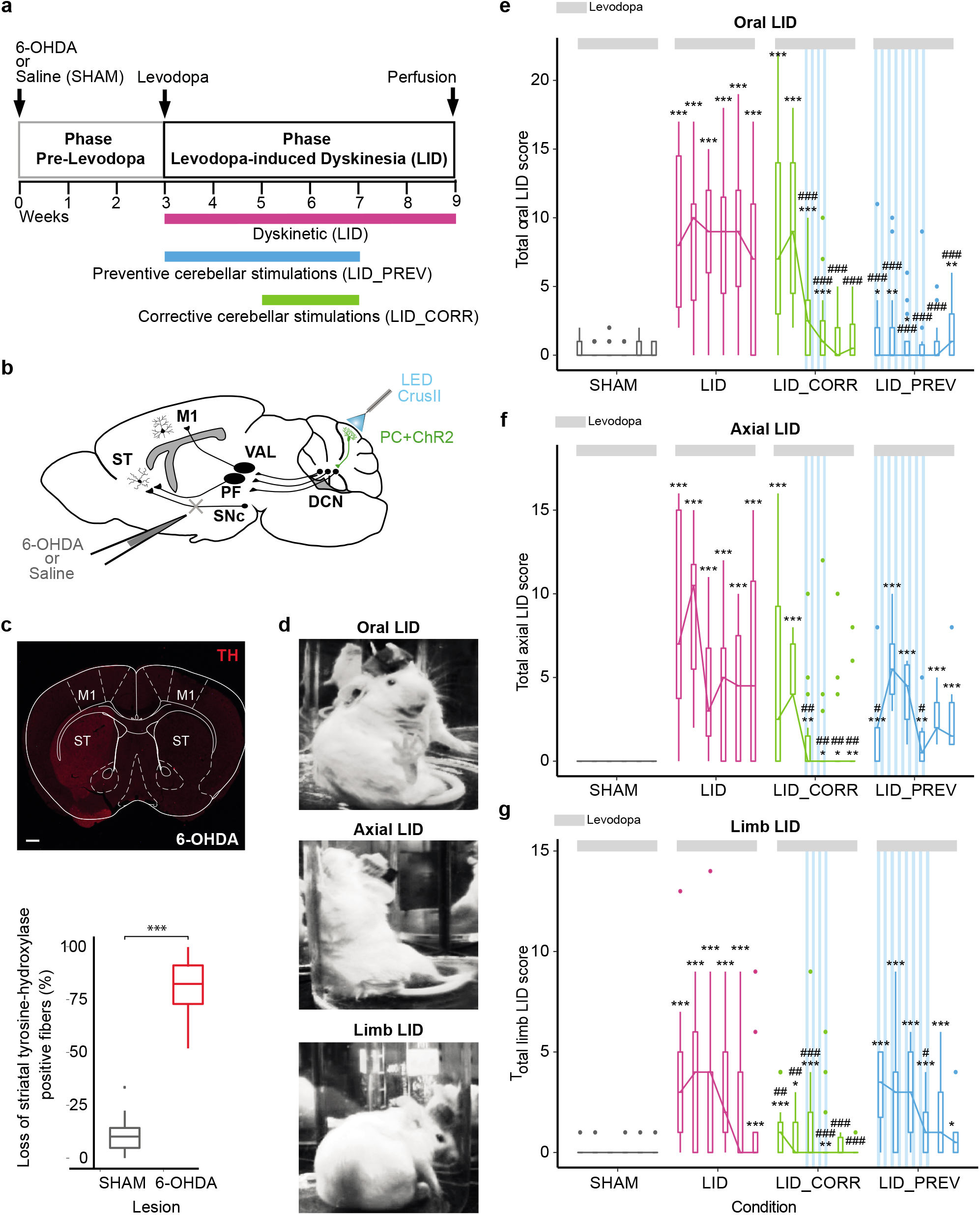
Optogenetic stimulations of Crus II Purkinje cells both reduce and prevent severe oral peak-dose dyskinesia. **a** Experimental timeline. Dyskinetic mice (LID, magenta): 6 weeks of levodopa treatment. Preventive mice (LID_PREV, blue): 6 weeks of levodopa treatment + 4 weeks of cerebellar stimulations. Corrective mice (LID_CORR, green): 6 weeks of levodopa treatment + 2 weeks of cerebellar stimulations. **b** Sagittal schematic of a mouse brain showing cerebello-thalamo-cortical and -striatal pathways, ChR2-YFP in Purkinje cells (PC+ChR2, green), and injection site of 6-OHDA or saline. M1: Primary motor cortex, ST: striatum, VAL: Ventroanterior-ventrolateral complex of the thalamus, PF: Parafascicular nucleus of the thalamus, SNc: Substantia nigra *pars compacta*, DCN: deep cerebellar nuclei, CrusII: Crus2 of the ansiform lobule. **c** *Upper panel*: Coronal section from a mouse unilaterally-lesioned with 6-OHDA stained with anti-tyrosine hydroxylase (TH). Scale bar: 0.5 mm. M1: Primary motor cortex, ST: Striatum. *Bottom panel*: Loss of striatal TH-positive fibers (%) between the lesioned and the intact striatum in control mice (grey, N=17) and parkinsonian animals (red, N=40). **d** Examples of orolingual (top), axial (middle), and limb (bottom) levodopa-induced dyskinesia in dyskinetic mice. **e** Boxplot showing the sum of oral LID scores across the 6 weeks of levodopa treatment (light grey bar) for SHAM (grey, N=18), LID (magenta, N=19); LID_CORR (green, N=17); LID_PREV (bleu, N=24). Stripped blue lines: weeks of theta-burst PC stimulations. **f** Boxplot showing the sum of axial LID across the 6 weeks of levodopa treatment (light grey bar) for SHAM (grey, N=12), LID (magenta, N=8), LID_CORR (green, N=14), LID_PREV (bleu, N=6). Stripped blue lines: weeks of theta-burst PC stimulations. **g** Boxplot showing the sum of limb LID scores across the 6 weeks of levodopa treatment (light grey bar) for SHAM (grey, N=14), LID (magenta, N=9), LID_CORR (green, N=15), LID_PREV (bleu, N=9). Stripped blue lines: weeks of theta-burst PC stimulations. Boxplots represents the lower and the upper quartiles as well as the median of LID score. Kruskal-Wallis test with pairwise Wilcoxon test and Benjamini & Hochberg correction. ***p < 0.001; **p < 0.01; *p < 0.05; * compared to SHAM; # compared to LID. See also Table S1.

**Fig. 2.**
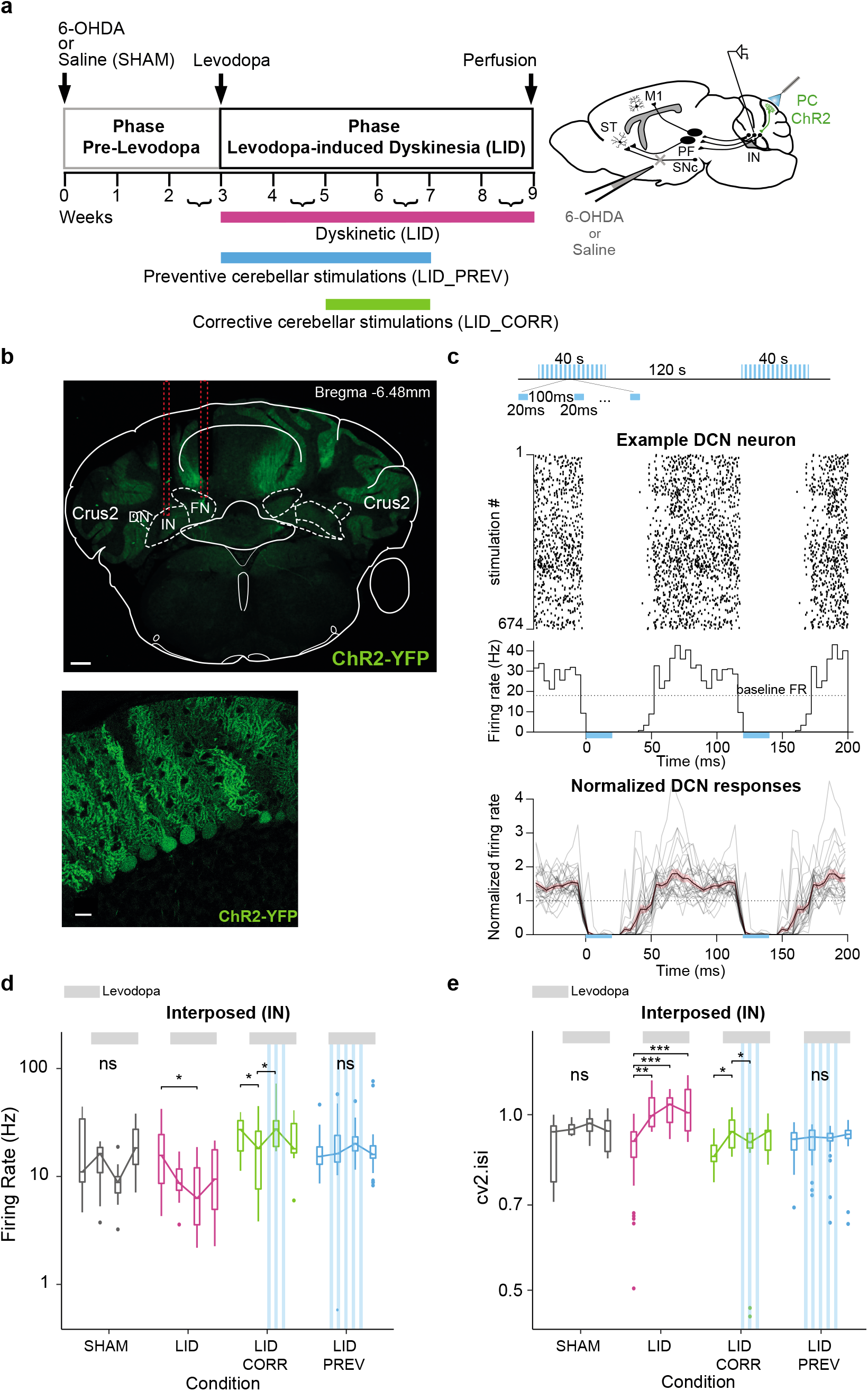
Purkinje cell stimulations normalize firing rate and regularize pattern of activity in the interposed nucleus. **a** *Left*: Experimental timeline. *Right*: Schematic of electrode implantation in the interposed nucleus (IN), ChR2-YFP expression in Purkinje cells (PC+ChR2, green) and injection site of 6-OHDA or saline. ST: Striatum; SNc: substantia nigra *pars compacta*; M1: Primary motor cortex; PF: Parafascicular nucleus of the thalamus. **b** *Top*: Coronal section from L7-ChR2-YFP mouse. Red lines: electrode’s trajectory. Dotted white lines: IN and fastigial (FN) nuclei. Scale bar: 0.5 mm. Crus2: Crus2 of the ansiform lobule, DN: Dentate cerebellar nucleus. *Bottom*: PC expressing YFP. Scale bar: 20µm. **c** *Top*: Theta-burst protocol. *Middle*: Raster plot of a deep cerebellar nuclei (DCN) neuron for each stimulation. Dotted line: Basal firing rate (FR) before the onset of stimulations. Blue box: Time of optogenetic stimulation. *Bottom*: Summary of DCN firing profiles (n=27; N=3) exhibiting a strong inhibition (>90%) during PC stimulations. The firing rate of each unit was normalized to its baseline. Shaded lines: mean +/- std. **d** Firing rate (Hz) across 9 weeks in the IN. Boxplots show the median rate (horizontal bars) over 4 categories of weeks. First boxplot: 2^nd^ and 3^rd^ weeks; second boxplot: 4^th^ and 5^th^ weeks when levodopa begins; third boxplot: 6^th^ and 7^th^ weeks; last boxplot: 8^th^ and 9^th^ weeks when stimulations stopped. Grey = SHAM (N=5); Magenta = LID (N=3); Green = LID_CORR (N=4); Blue = LID_PREV (N=3). Light grey lines: 6 weeks of levodopa treatment (3 boxplots). Stripped blue lines: weeks of theta-burst stimulations. **e** Coefficient of variation 2 (cv2.isi) across 9 weeks in the IN. Same order of boxplot as panel **d**. Grey = SHAM (N=5); Magenta = LID (N=3); Green = LID_CORR (N=4); Blue = LID_PREV (N=3). Light grey lines: 6 weeks of levodopa treatment (3 boxplots). Stripped blue lines: weeks of theta-burst stimulations. Boxplots represents the lower and the upper quartiles. Welch Anova with Games Howell post-hoc test and one-way Anova’s with Tukey post-hoc test based on Levene test. ***p < 0.001; **p < 0.01; *p < 0.05; ns: p > 0.5. See also Tables S2, S3, and S4.

In 6-OHDA-lesioned mice exhibiting severe dyskinesia upon repeated levodopa injections, 2 weeks of daily corrective PC stimulations on the cerebellar orolingual region Crus II significantly reduced oral dyskinesia (condition “LID_CORR”, N=24, Figures 1e, S1b, Table S1). This effect persisted at least for 2 weeks after the end of cerebellar stimulations (weeks 8 and 9, Figures 1e, S1b, Table S1) and the dyskinesia scores were then similar to those of the control group (Table S1). Furthermore, after corrective PC stimulations over orolingual CrusII, the reduction in oral LID was more pronounced than in axial and limb dyskinesia (Figures 1f-g, S2b, S3b).

Another group of 6-OHDA-lesioned mice received daily cerebellar stimulations starting from the first day of levodopa administration (3mg/kg) i.e before the development of dyskinesia (condition “LID_PREV”, N=18). Remarkably, this group exhibited only few to none orolingual dyskinesia, contrarily to LID animals (Figures 1e, S1b, Table S1). In conclusion, 2 weeks of daily cerebellar stimulations led to a dramatic decrease of LID expression that outlasted the stimulations for at least 2 weeks, while stimulations starting concomitantly with levodopa administration prevented LID development. Therefore, these results indicate a strong suppressive effect of peak-dose dyskinesia by cerebellar PC stimulations.

Previous studies in animals models of LID addressed exclusively “peak-dose” dyskinesia ^61^. Yet, mild dyskinesia also occurred outside the 2 hours following the injection time as in PD patients at the trough of blood levodopa concentration (“off-period” dyskinesia) or during the rising and falling phase of blood levodopa concentrations (diphasic dyskinesia) review in ^62^. Therefore, analysis of dyskinesia observed 20 minutes before levodopa injection revealed that chronic PC stimulations on CrusII also suppressed or prevented oral “off-period” dyskinesia depending on the protocol used (Figure S1c, Table S1).

In conclusion, daily sessions of opto-stimulations of PC in CrusII, which corresponds to the orolingual region of the cerebellar cortex, is sufficient to obtain a significant decrease of oral LID. These results bear resemblance with those obtained in PD patients in whom rTMS targeting posterior cerebellum improved LID scores ^18, 19^ but show a stronger effect than in humans were the severity of dyskinesia was only reduced at the peak effect of levodopa.

### Purkinje cell stimulations over CrusII modulate aberrant activity of the cerebellar nuclei

To test whether and how systemic levodopa treatment results in changes of activity in the cerebellum, we chronically recorded neurons in the three deep cerebellar nuclei (DCN): the interposed nucleus (IN), the dentate nucleus (DN), and the fastigial nucleus (FN) (Figures 2b, S5b). Neuronal activity was recorded both before and after levodopa administration in freely moving 6-OHDA-lesioned and control mice, for a total of 9 weeks (Figures 2a,b, S5a-c). Recordings in three mice from DCN neurons during the stimulation protocol revealed that cells in the three DCN, strongly inhibited by the stimulation (hence likely receiving inputs from the stimulated area), exhibited an alternation of cessation of firing and increased firing relative to the baseline activity along the protocol (Figure 2c). We observed that levodopa decreased the global activity of IN and DN, but not FN, in LID animals (Figures 2d, S5d-f, Tables S2, S3). We verified that this did not reflect changes in motor activity (**Supp. Text,** Figure S6). Altogether, these results constitute the first evidence of a dysregulated activity of the output nuclei of the cerebellum by levodopa in LID.

#### Purkinje cell stimulations over CrusII prevent the decrease of activity in the interposed nucleus

For animals receiving 2 weeks of PC stimulations (LID_CORR), we found that the depressed activity in IN induced by levodopa treatment was restored during the period of cerebellar stimulations, but this effect did not last after the end of the stimulations (Figures 2d, Tables S2, S3). The effect was stronger in animals receiving 4 weeks of preventive PC stimulations (LID_PREV). The most striking effect was observed in IN where PC stimulations prevented the global decrease of firing rate (Figure 2d, Tables S2, S3). The effects of PC stimulations were less clear in DN and FN (Figure S5d-f, Tables S2, S3). Taken together, these data show that repeated sessions of cerebellar stimulations affected the aberrant activities observed under levodopa treatment in the three DCN. However, only IN exhibited a consistent normalization, which suggests a prominent role of this structure in the normalization of the dyskinesia.

#### Levodopa treatment causes DCN neurons to develop a more erratic activity in dyskinetic mice

The irregularity of neural discharge in the cerebellum is detrimental to motor control^63^ and has been observed in rapid-onset dystonia-Parkinsonism ^64, 65^ and tremor ^66^. The average normalized difference of successive interspike intervals (cv2.isi) is a measure of irregularity of directly adjacent interspike intervals, and therefore higher cv2.isi value indicates a more irregular cell activity ^67^. Interestingly, LID mice exhibited a higher cv2.isi in IN, DN and FN during the entire period of levodopa treatment (Figures 2e, S5e-g, Tables S2, S4). The higher values of cv2.isi did not simply reflect increased bursting (**Supp. Text,** Figure S8a). Therefore, these results showed a more erratic and irregular pattern in DCN neurons in LID mice, mostly during periods of activity, during levodopa treatment.

#### Purkinje cells stimulations in CrusII prevent changes in pattern of activity in the interposed nucleus

In mice receiving 2 weeks of PC stimulations (LID_CORR), the cv2.isi significantly increased the first 2 weeks of levodopa treatment in IN and was normalized during PC stimulations (Figure 2e, Tables S2, S4). In contrast, cv2.isi values remained significantly elevated in DN and FN (Figure S5e-g, Tables S2, S4).

In mice receiving 4 weeks of PC stimulations (LID_PREV), we found that the increased cv2.isi in IN was prevented by PC stimulations (Figure 2e, Tables S2, S4). However, increased cv2.isi was still present in DN and FN (Figure S5e-g, Tables S2, S4). Moreover, as in LID animals, locomotor activity did not change the cv2.isi values neither in LID_CORR nor in LID_PREV mice.

Overall, these electrophysiological data suggest that dyskinesia-related abnormal activity is conveyed to the DCN, especially to IN, leading to aberrant firing rate and firing patterns, which are reversed by chronic PC stimulations. Finally, these experiments suggest a tighter association between the changes in IN activity and the alleviation of the pathological phenotype.

### Chronic levodopa treatment increases the activity of the oral motor cortex and decreases the firing rate in the parafascicular thalamic nucleus of dyskinetic mice

Since LID strongly involve the forebrain motor circuits ^16^, and cerebellar nuclei have multiple ascending projections toward these circuits ^68, 69^, we next investigated the impact of cerebellar stimulations in the thalamus and motor cortex. Changes in motor cortex activity, intralaminar nuclei of the thalamus, including the parafascicular nucleus (PF), and the ventroanterior-ventrolateral (VAL) complex of the thalamus have been observed in dyskinetic patients ^21, 22, 46^ and animals ^23, 24, 47^. Moreover, cerebello-cortical loops ^70, 71^ and parafascicular projections to the striatum and to the cerebral cortex ^72^ are topographically organized. Therefore, to examine the impact of cerebellar stimulations on the thalamus and motor cortex, we chronically recorded neurons in the oral region of M1, and in the thalamic PF and VAL, during 5 weeks of chronic levodopa treatment (Figures 3a,b, S9a,b).

**Fig. 3.**
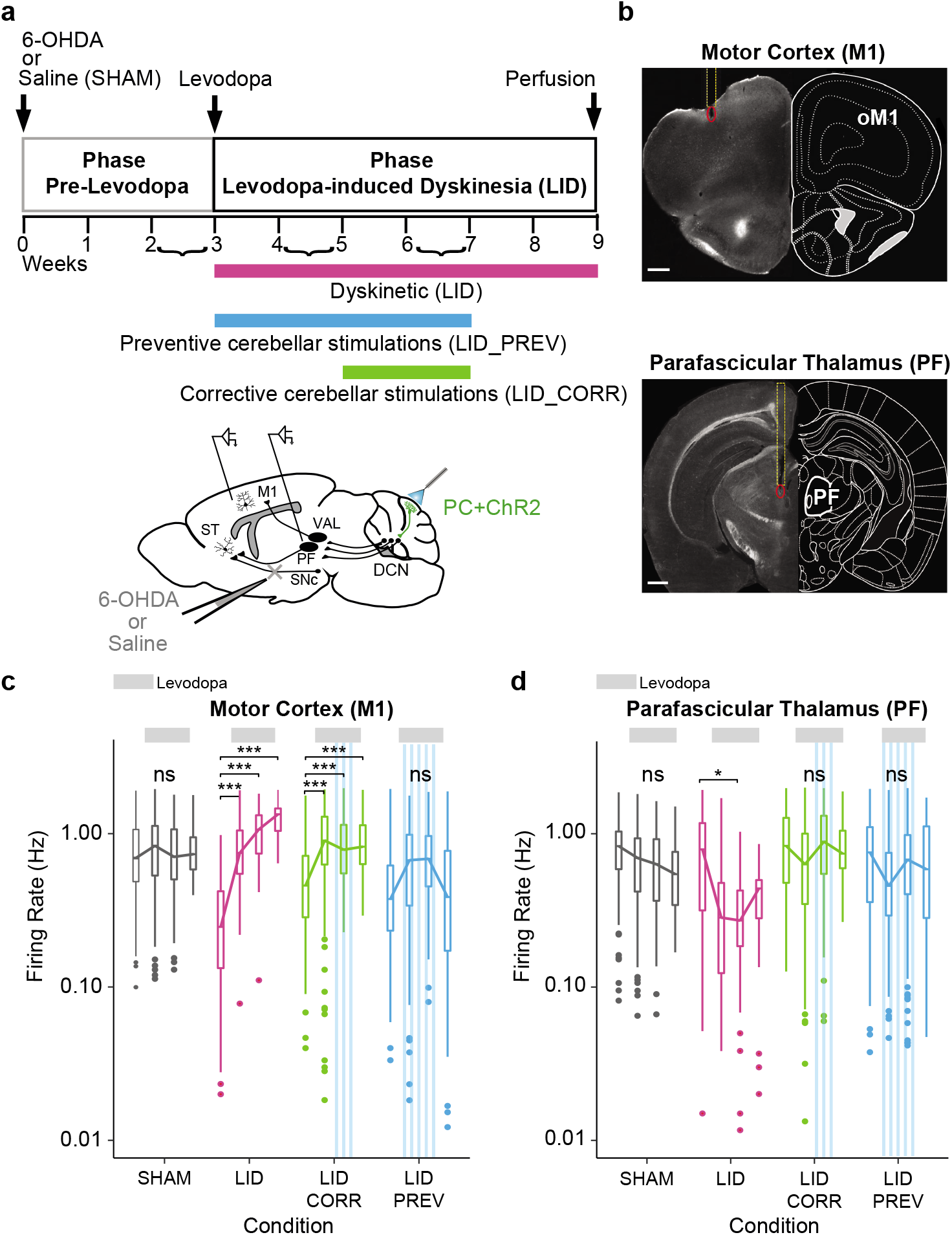
Aberrant activity in the motor cortex and the parafascicular nucleus of the thalamus in dyskinesia is restored by Purkinje cell stimulations. **a** *Top*: Experimental timeline. *Bottom*: Schematic of electrode implantation in the primary motor cortex (M1) and the parafascicular nucleus of the thalamus (PF), ChR2-YFP expression in the Purkinje cells (PC+ChR2, green), and injection site of 6-OHDA or saline. ST: Striatum; SNc: substantia nigra *pars compacta*; DCN: deep cerebellar nuclei: VAL: Ventroanterior-ventrolateral complex of the thalamus. **b** *Top*: Coronal section from L7-ChR2-YFP mouse showing the electrode’s trajectory (dotted yellow line) and the lesion site (red circle) in layer 5 of the oral M1 (oM1). Scale bar: 0.5 mm. *Bottom*: Coronal section from L7-ChR2-YFP mouse showing the electrode’s trajectory (dotted yellow line) and the lesion site (red circle) in the parafascicular nucleus of the thalamus (PF). Scale bar: 0.5 mm. **c** Firing rate (Hz) across 9 weeks in M1. Boxplots show the median rate (horizontal bars), over 4 categories of weeks. First boxplot: 2^nd^ and 3^rd^ week of the protocol, second boxplot: 4^th^ and 5^th^ weeks when levodopa begins, third boxplot: 6^th^ and 7^th^ weeks, last boxplot: 8^th^ of the protocol when stimulations stopped. Grey = SHAM (N=5); Magenta = LID (N=4); Green = LID_CORR (N=6); Blue = LID_PREV (N=8). Light grey lines: 6 weeks of levodopa treatment (3 boxplots). Stripped blue lines: weeks of theta-burst PC stimulations. **d** Firing rate (Hz) across 9 weeks in the parafascicular nucleus of the thalamus (PF). Same order of boxplot as panel **c**. Grey = SHAM (N=5); Magenta = LID (N=4); Green = LID_CORR (N=6); Blue = LID_PREV (N=8). Light grey lines: 6 weeks of levodopa treatment (3 boxplots). Stripped blue lines: weeks of theta-burst PC stimulations. Boxplots represents the lower and the upper quartiles. One-way Anova with Tukey HSD post-hoc test. ***p < 0.001; **p < 0.01; *p < 0.05; ns: p > 0.5. See also Tables S5 and S6.

The activity in M1 and PF varied slightly over the course of levodopa treatment in SHAM animals, whereas LID mice exhibited a significant increase of the firing rate in M1 (Figure 3c, Tables S5, S6) and a significant decrease in the firing rate in PF after levodopa administration (Figure 3d, Tables S5, S6). The effects were more inconsistent in VAL (Figure S9d, Tables S5, S6). Altogether, these results confirm the presence of functional alterations in PF and oral M1 in dyskinesia.

### Purkinje cell stimulations over CrusII prevent both the abnormal increase of activity in M1 and the decrease in PF in dyskinetic mice

We then investigated whether the abnormal activities observed in M1 and PF could also be normalized by chronic PC stimulations. Chronic extracellular recordings in dyskinetic mice receiving 2 weeks of PC stimulations (LID_CORR) showed that the global firing rate of oral M1 significantly increased the first 2 weeks of levodopa treatment (Figure 3c, 2^nd^ boxplot), as observed in LID animals (Figure 3c, Tables S5, S6). However, contrarily to these animals, no further increase was found after the stimulations started (Figure 3c, 3^rd^ boxplot). In PF, the decrease observed in dyskinetic mice was not observed during cerebellar stimulations (Figure 3d, Tables S5, S6).

In animals receiving 4 weeks of preventive cerebellar stimulations (LID_PREV), both the increased activity in oral M1 observed in LID (Figure 3c, Tables S5, S6) and the decreased activity in PF in LID were prevented (Figure 3d, Tables S5, S6), and remained normal even after the end of the stimulations. As for LID mice, the effects observed in the motor thalamus VAL were more variable in the corrective and the preventive conditions, suggesting that both levodopa and repeated sessions of cerebellar stimulations have less impact on this structure (Figure S9d, Tables S5, S6).

Taken together, these results suggest that the effects induced by cerebellar stimulations restore activities of both oral M1 and PF, by being able to reverse the changes in firing rate associated with dyskinesia.

### The cerebellum is connected to the parafascicular nucleus of the thalamus

Because both DCN and PF showed a similar modulation of their firing rate, we examined how IN, DN, and FN project to PF. For this purpose, we used retrograde viral tracing that allowed us to determine the distribution of DCN neurons projecting to PF (Figure 4a). Quantification of retrograde labeled neurons from PF showed 42.1 ± 9.0 % of projecting neurons in IN, 41.4 ± 2.6 % of neurons in DN, and 16.5 ± 1.7 % of neurons in FN (N=6, Figure 4b-d), suggesting that IN and DN might have an important contribution in the cerebellar control of PF activity. Moreover, PF-projecting DCN neurons were localized along the entire antero-posterior axis in the three nuclei (Figure 4b,c,e).

**Fig. 4.**
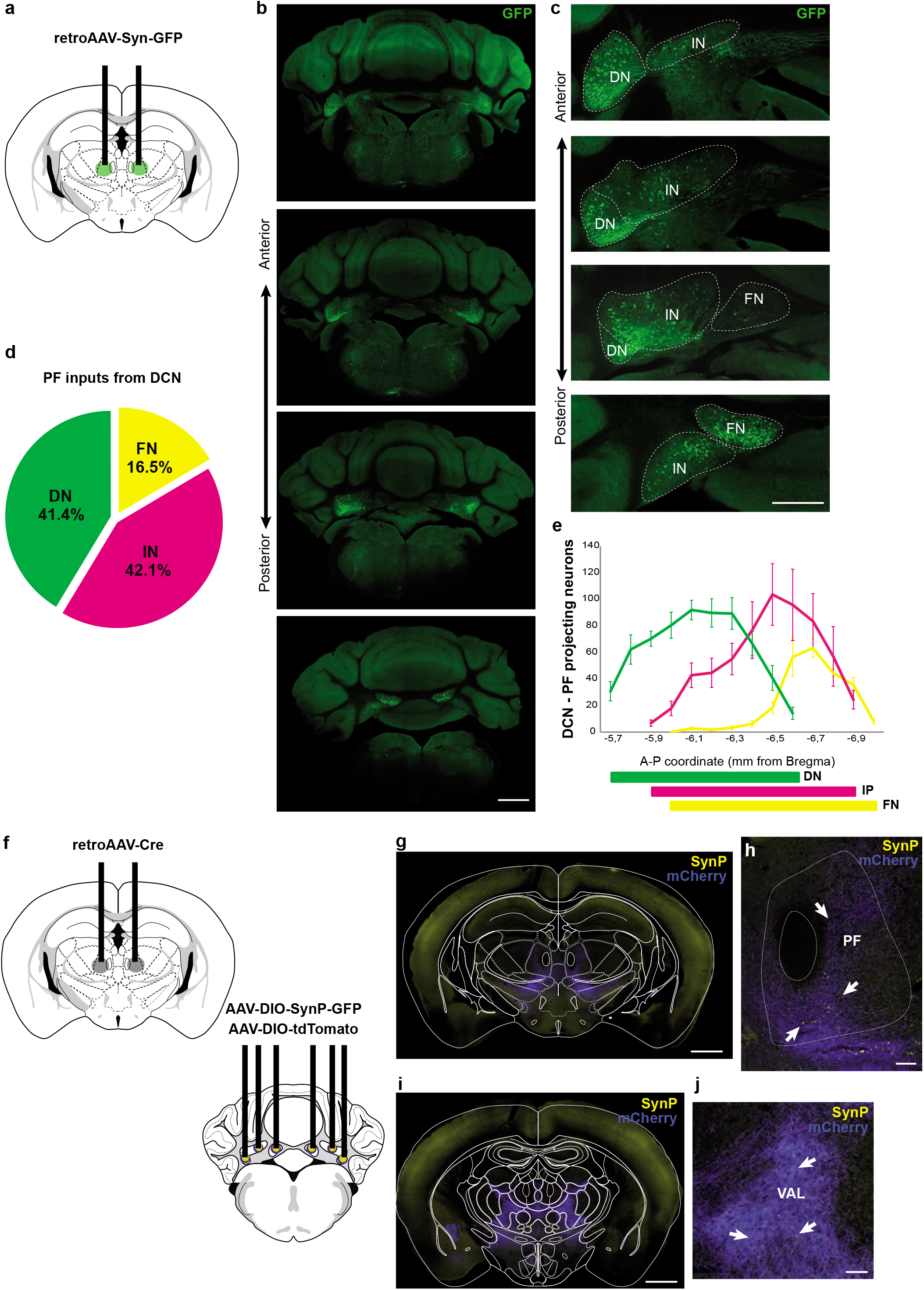
DCN monosynaptic inputs to PF and collaterals. **a** Retrograde labeling strategy by viral injection of retrograde AAV-syn-GFP in the parafascicular nucleus of the thalamus (PF). **b** Anterior to posterior cerebellar sections showing retrograde labeled neurons in the three deep cerebellar nuclei (DCN): dentate (DN), interposed (IN), and fastigial (FN). Scale bar: 1 mm. **c** High-magnification on DCN from **b**. Scale bar: 0.5 mm. **d** Quantification of the cell fraction (%) of retrograde labeled DN, IN and FN neurons projecting to PF. **e** Distribution of retrograde labeled DCN neurons projecting to PF in each nucleus. **f** Tracing of axon collaterals from DCN-PF projecting neurons by expression of retrograde AAV-Cre in PF, AAV-DIO-SynP-GFP and AAV-DIO-tdTomato in IN, FN, and DN. **g** Posterior thalamic section showing DCN neurons projecting to PF expressing Cre-dependent Synaptophysin (SynP)-GFP and tdTomato. Scale bar: 1mm. **h** Zoom-in of PF section exhibiting synaptic boutons (arrows) in DCN inputs Scale bar: 100 µm. **i-j** Anterior thalamic section showing axon collaterals within the thalamus. Scale bar: 1mm. **j** Zoom-in from **i** of VAL section exhibiting some synaptic boutons (arrows) from DCN-PF axon collaterals. Scale bar: 1mm.

To confirm the presence of DCN synaptic terminals in PF neurons, we localized the synaptic terminals of DCN neurons projecting to PF using Cre-dependent viral expression of the presynaptic marker synaptophysin (SynP)-GFP in combination with the expression of Cre-recombinase obtained by retrograde viral injections in PF (Figure 4f). Large amount of DCN terminals expressing SynP-GFP were found in PF (Figure 4g,h), and much less in other thalamic nuclei as VAL (Figure 4i,j). Overall, these results confirm the presence of cerebellar projections to PF and demonstrate that they originate from a population distinct from the one projecting to VAL.

### DCN-PF pathway inhibition counteracts the effects of Purkinje cell stimulations on oral dyskinesia in preventive mice

Since DCN may entrain PF, and since PF stimulations have been used to avoid LID ^46, 47^, we examined the involvement of the DCN-PF pathway in the effect of cerebellar stimulations in LID. We therefore examined the effect of transiently inactivating the specific projections from DCN to PF during the repeated sessions of PC opto-stimulations. For this purpose, we expressed inhibitory hM4Di DREADD receptors in DCN neurons that target PF, by injecting retrograde CAV2-Cre-GFP in PF in complementation with a Cre-dependent AAV-DIO-hM4Di-mCherry in DCN (Figure 5a-d) in mice receiving 4 weeks of PC stimulations.

**Fig. 5.**
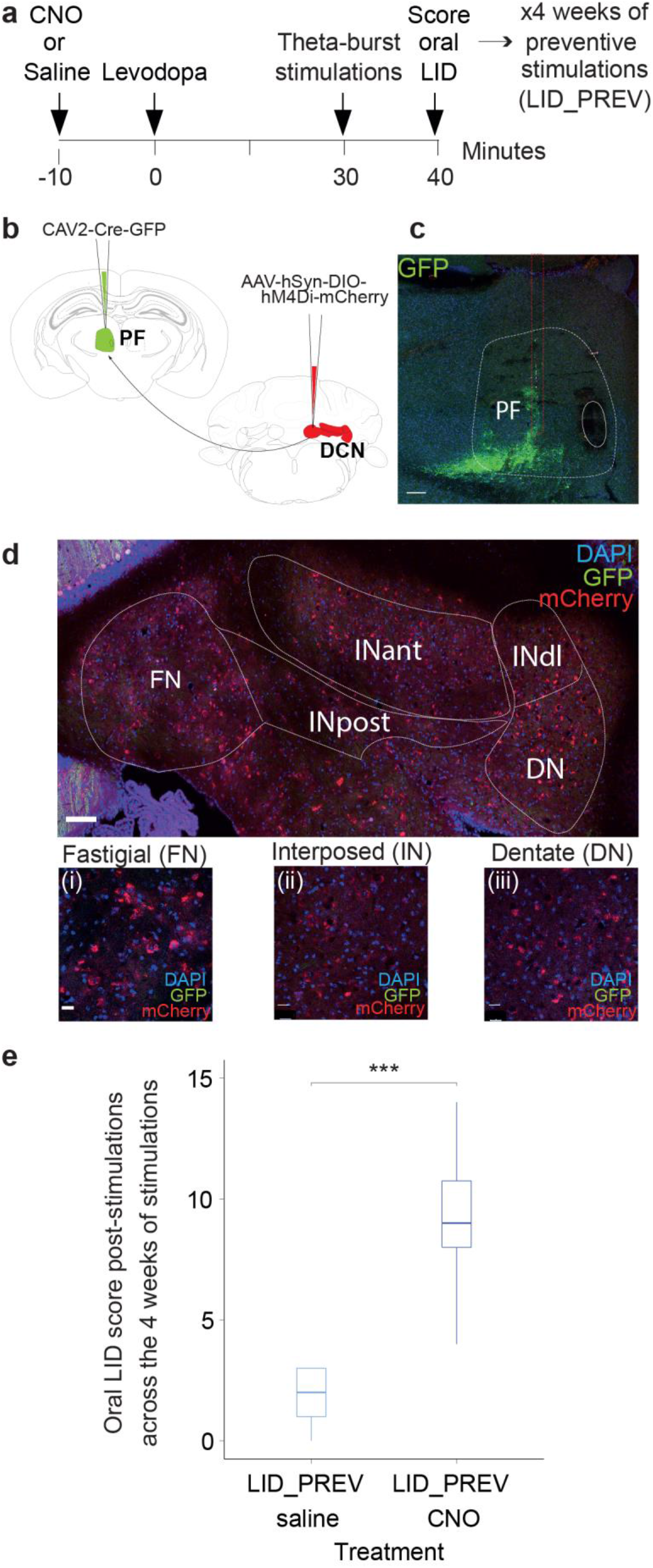
DCN to PF pathway inhibition counteracts the effects of Purkinje cells stimulation on the severity of oral dyskinesia in preventive mice. **a** Experimental timeline. **b** Schematic of mouse coronal sections showing the injection site of the retrograde CAV2-Cre-GFP in the parafascicular nucleus of the thalamus (PF, green) ipsilateral-to-the-lesion (top, left) and the injection site of the anterograde Cre-dependent AAV-hSyn-DIO-hM4Di-mCherry in the three deep cerebellar nuclei (DCN, red) contralateral-to-the-lesion. **c** Coronal section from L7-ChR2-YFP mouse showing the injection site of retrograde CAV2-Cre-GFP in PF. Red dotted lines: needle’s trajectory. White dotted lines highlight the limits of PF within the thalamus. Scale bar: 100 μm. **d** Coronal section from L7-ChR2-YFP mouse showing the expression of anterograde pAAV-hSyn-DIO-hM4Di-mCherry in neurons (red) of DCN. Med: medial cerebellar nucleus; IntP: interposed cerebellar nucleus, posterior part; IntA: interposed cerebellar nucleus, anterior part; IntDL: interposed cerebellar nucleus, dorsolateral hump; Lat: lateral cerebellar nucleus. Blue = DAPI; green = GFP; Red = mCherry. Scale bar: 100 μm. Insert: Postmortem histology showing hM4Di-mCherry-expressing neurons in the fastigial nucleus (i, FN), the interposed nucleus (ii, IN), and the dentate nucleus (iii, DN). Scale bars: 20µm. **e** Boxplots showing the average score of oral LID severity at the point time corresponding to 40 minutes after levodopa injection, 50 minutes after CNO injection and right after chronic theta-burst stimulations of Purkinje cells. Average score comprises the 4 weeks of preventive cerebellar stimulations in two groups: light blue represents preventive animals receiving 4 weeks of cerebellar stimulations + daily injections of both levodopa and saline (LID_PREV saline, N=5), dark blue represents preventive animals receiving 4 weeks of cerebellar stimulations + daily injections of both levodopa and CNO (LID_PREV CNO, N=6). Horizontal bars in boxplots represent the median score. Boxplots represents the lower and the upper quartiles. Non-parametric Kruskal-Wallis test with pairwise Wilcoxon test and a Benjamini & Hochberg correction were used. ***p<0.001; **p < 0.01; *p < 0.05; ns: p > 0.05.

As oral LID peaked around 30 minutes after levodopa injection, we scored oral LID severity in preventive mice at this time point, and after chronic cerebellar stimulations were applied, to highlight the difference of effects between chemogenetic inhibition and optogenetic activation. Mice injected with inhibitory DREADDs, stimulated for 4 weeks and receiving daily CNO injections before the stimulations, presented significantly more severe oral dyskinesia than control mice also injected with inhibitory DREADDs, stimulated for 4 weeks but receiving only daily saline injections (p<0.001; Figure 5e). These results indicate that DCN to PF inputs are involved in the preventive effect of PC stimulations.

### Purkinje cell stimulations alter corticostriatal transmission and plasticity in brain slices

As PF projects to the striatum ^72,73,74^, it may relay a cerebellar control over corticostriatal synaptic plasticity ^49^. Indeed, alterations in corticostriatal plasticity are found in hyperkinetic disorders such as LID ^14, 43^. LID have notably been associated with an excessive corticostriatal long-term potentiation (LTP) in the direct pathway medium spiny neurons (MSN) without prominent change in the indirect pathway, resulting in an increased motor activity ^43^. Direct and indirect pathway MSNs express different dopaminergic receptors, the dopamine receptor subtype-1 (D1R) or subtype-2 (D2R) respectively for the direct and indirect pathways ^75^. Therefore, we examined *ex vivo* the corticostriatal synaptic plasticity in MSNs of the dorsolateral striatum belonging either to the direct or indirect striatal pathways using brain slices from L7-ChR2xDrd2-GFP mice subjected to 4 conditions: SHAM, SHAM_PREV, LID or LID_PREV (Figure 6a). We previously reported that using a spike-timing dependent plasticity (STDP) paradigm, paired pre-synaptic activations preceded by post-synaptic activations induced LTP in both direct and indirect pathway MSN ^76, 77^. LTP was induced in MSNs of SHAM, SHAM_PREV, LID or LID_PREV mice except in direct pathway MSNs issued from LID_PREV mice, where a clear long-term depression (LTD) was found (Figure 6b-e **and** Table S12). Interestingly, in direct pathway MSNs, LTP induced in LID mice exhibited greater magnitude (p<0.0001) than in SHAM mice, whereas preventive PC stimulations either reduced LTP in SHAM mice (p<0.0001) or even reversed LTP into LTD in LID mice. The magnitude of corticostriatal LTP induced in indirect pathway MSNs did not show significant variation in SHAM and LID mice with or without preventive PC stimulations (ANOVA: F=0.3044, 38 degree of freedom, p=0.8220).

**Fig. 6.**
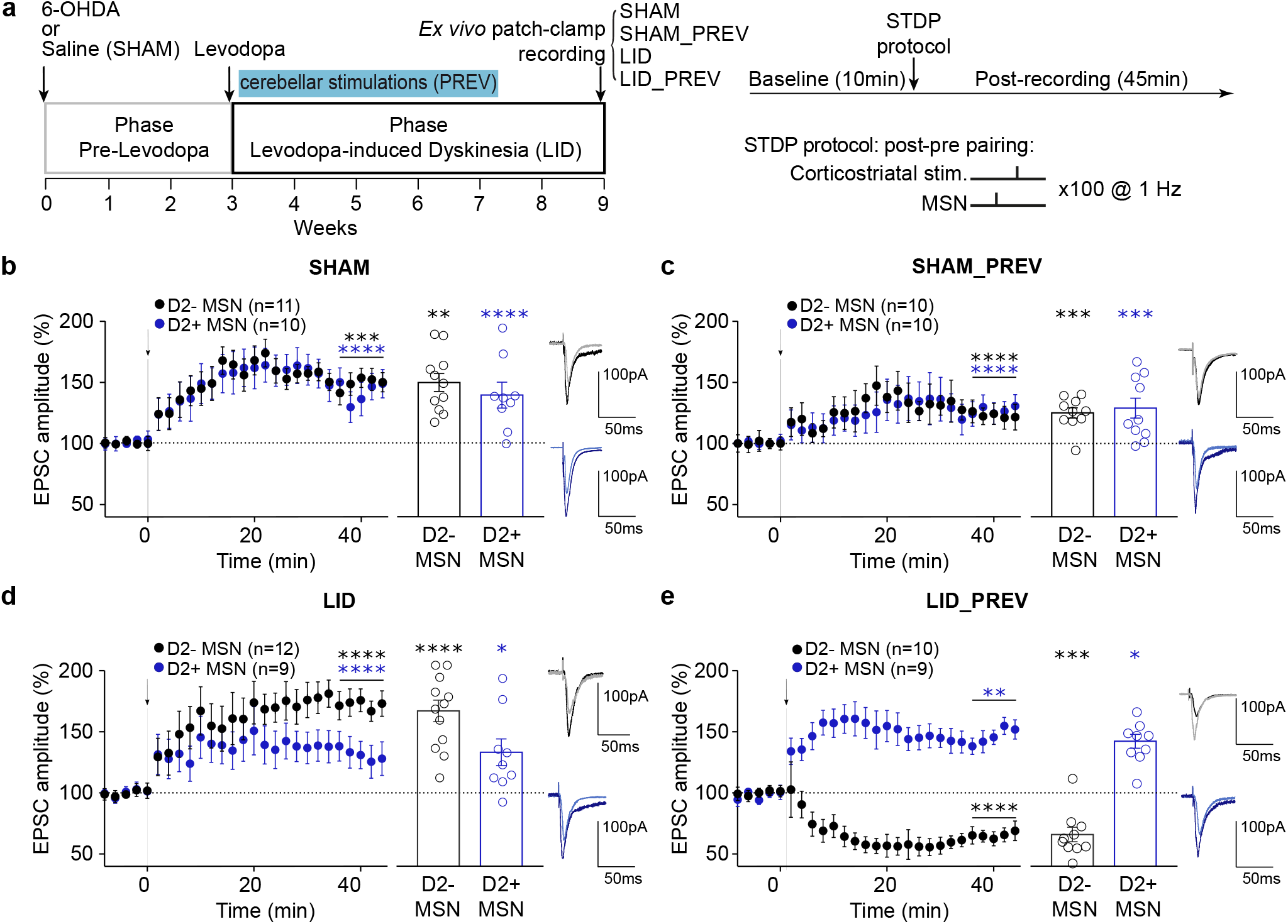
Spike-timing dependent plasticity produce LTD instead of LTP in D1-expressing neurons following Purkinje cell stimulations. **a** *Left*: Experimental timeline. Control mice (SHAM): 6 weeks of levodopa treatment. Preventive control mice (SHAM_PREV): 6 weeks of levodopa treatment + 4 weeks of cerebellar stimulations. Dyskinetic mice (LID): 6 weeks of levodopa treatment. Preventive mice (LID_PREV): 6 weeks of levodopa treatment + 4 weeks of cerebellar stimulations. *Ex vivo* experiments were realized on mice subjected to 4 conditions, i.e. SHAM, SHAM_PREV, LID and LID_PREV. *Right*: STDP pairings: A single spike evoked in the recorded MSN (post) was paired with a single cortical stimulation (pre); pairings were repeated 100 times at 1 Hz. **b-e** Averaged time courses of corticostriatal STDP in D1-MSNs and D2-MSNs induced by 100 post–pre pairings. **b** In SHAM, LTP induced by 100 post–pre pairings in D1-MSNs and D2-MSNs (n=21). **c** In SHAM_PREV, LTP induced by 100 post–pre pairings in D1-MSNs and D2-MSNs (n=20). **d** In LID, LTP induced by 100 post–pre pairings in D1-MSNs (n=12) and D2-MSNs (n=9). **e** In LID_PREV, LTP induced by 100 post–pre pairings in D1-MSNs (n=10) and the same protocol induced LTD in D2-MSNs (n=9). Synaptic strength was determined 34-44 min after pairings. Error bars represent the SEM. **p < 0.01; ***p < 0.001; ****p < 0.0001 by one sample t test.

Therefore, we found that cerebellar preventive stimulations reverse striatal pathological LTP into LTD in direct pathway neurons, an effect which may then prevent the consolidation of an abnormal motor activity in the direct pathway and help reinstating normal motor functions.

### Purkinje cell stimulations normalize the expression of the dyskinetic marker FosB/ΔFosB in the dorsolateral striatum

The expression of the transcription factors FosB/ΔFosB, from the immediate early gene *fosb*, has been used as a marker of dyskinesia ^38^. Alterations in its expression within the dorsolateral striatum affects LID, as both its inactivation ^41^ and its upregulation ^7, 40^ can respectively reduce and increase the severity of dyskinesia. We then examined whether PC stimulations also normalize the expression of the dyskinetic marker FosB/ΔFosB in the dorsolateral striatum (Figure 7a).

**Fig. 7.**
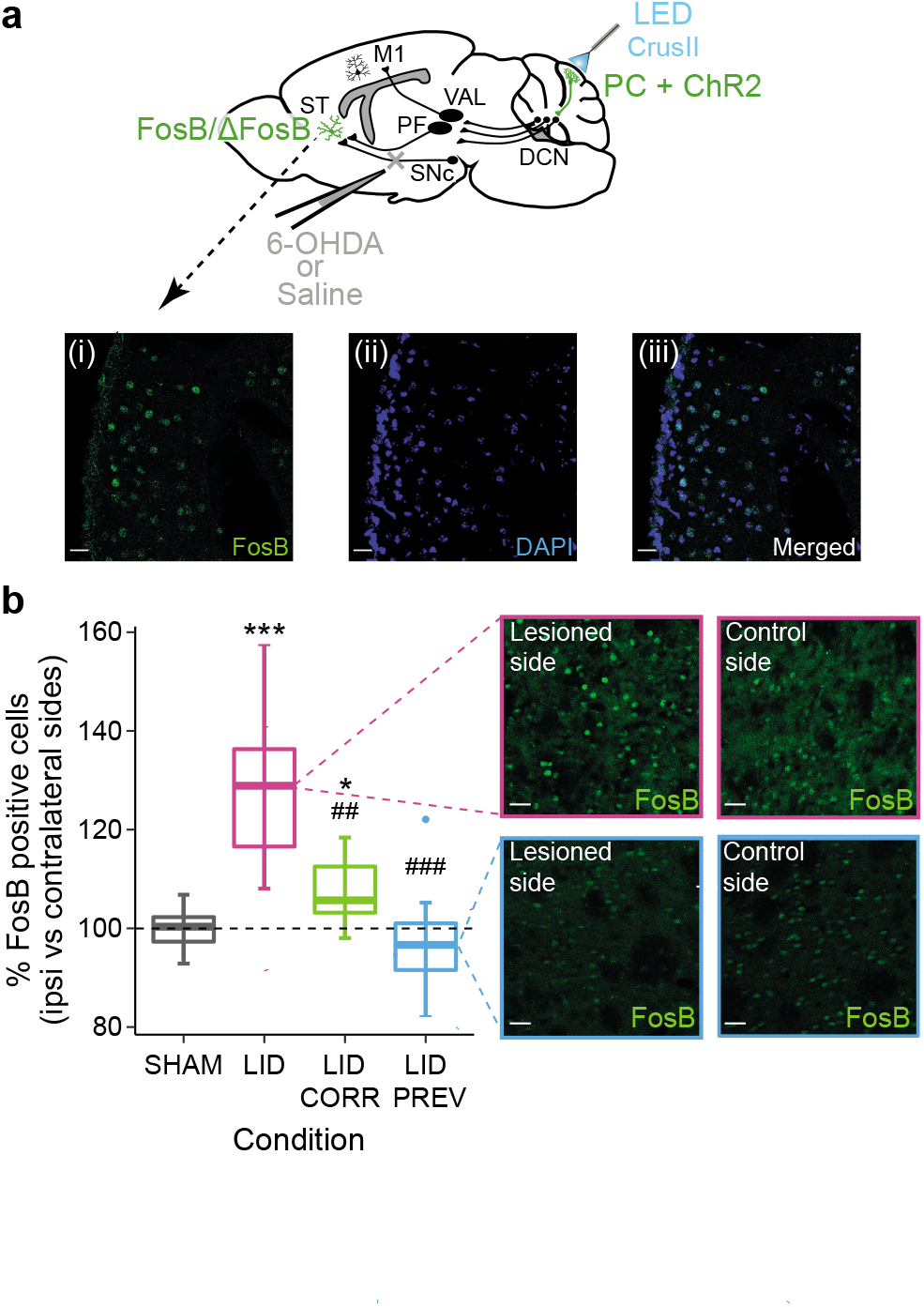
Striatal overexpression of the dyskinetic marker FosB/ΔFosB is restored by Purkinje cell stimulations. **a** Sagittal schematic showing neurons in the striatum expressing the dyskinetic marker FosB/ΔFosB (green). The cerebello-thalamo-cortical and cerebello-thalamo-striatal pathways are represented in mice expressing ChR2-YFP in Purkinje cells (PC+ChR2, green) as well as the injection site 6-OHDA or saline (grey). ST: Striatum; SNc: substantia nigra *pars compacta*; M1: Primary motor cortex, VAL: Ventroanterior-ventrolateral complex of the thalamus, PF: Parafascicular nucleus of the thalamus, DCN: deep cerebellar nuclei, CrusII: Crus2 of the ansiform lobule. Insets: Postmortem histology showing FosB-expressing neurons (i), DAPI (ii), and merged (iii). Scale bars: 20µm. **b** Boxplots showing the ratio of cells expressing FosB between the striatum ipsilateral to the lesion and the striatum contralateral to the lesion in percentage (%) in the 4 different conditions (magenta = LID, N=10; grey = SHAM, N=10; green = LID_CORR, N=7; and blue = LID_PREV, N=10). Horizontal bars in boxplots represent the median. Magenta inset: Postmortem histology showing FosB-expressing neurons in dyskinetic animals in the striatum ipsilateral to the lesion (left box) and in the striatum contralateral to the lesion (right box). Scale bars: 20µm. Blue inset: Postmortem histology showing FosB-expressing neurons in preventive animals in the striatum ipsilateral to the lesion (left box) and in the striatum contralateral to the lesion (right box). Scale bars: 20µm. Boxplots represents the lower and the upper quartiles. Student t-test. ***p < 0.001; **p < 0.01; *p < 0.05. * Compared to SHAM; # compared to LID. See also Table S7.

We compared FosB/ΔFosB expression in the dorsolateral striatum ipsilateral to the lesion with the dorsolateral striatum contralateral to the lesion, in all the different conditions. As expected, neither levodopa nor cerebellar stimulations impacted the expression of FosB/ΔFosB in SHAM animals and no asymmetry was found between the two striatum (SHAM, N=10, 99.6 ± 1.3 %, Figure 7b, Table S8). As previously demonstrated in other studies, LID mice presented an overexpression of FosB/ΔFosB in the dorsolateral striatum ipsilateral to the lesion (LID, N=10, 128.9 ± 5.0 %, Figure 7b, Table S8), significantly different from the control group. 2 weeks of cerebellar stimulations decreased the asymmetric expression of FosB/ΔFosB in the dorsolateral striatum of corrective mice compared to dyskinetic animals (LID_CORR, N=7, 107 ± 2.8 %, Figure 7b, Table S8), although still significantly different from the control animals. No significant asymmetry of striatal FosB/ΔFosB expression was found in mice receiving 4 weeks of cerebellar stimulations (LID_PREV, N=10, 94.2 ± 3.4 %, Figure 7b, Table S8), which did not significantly differ from SHAM mice. Moreover, striatal FosB/ΔFosB expression in LID_PREV animals was significantly different from the one observed in LID mice. Thus, our results suggest that PC stimulations are able to restore normal expression of striatal FosB/ΔFosB, which is accompanied by an anti-dyskinetic effect.

In conclusion, repeated sessions of PC stimulations over CrusII can both normalize the aberrant activity of major motor structures involved in LID, including DCN, PF and M1, but also the overexpression of the dyskinetic marker FosB/ΔFosB within the dorsolateral striatum, tightly linked to the development of LID. These effects were associated with the advent of LTD in the striatal direct pathway MSNs, which may prevent the overactivation of this pathway in LID. Altogether, these results indicate a widespread normalization in the cerebello-striato-cortical motor system and suggest that the cerebellar stimulations act on core mechanisms of LID.

## DISCUSSION

We used optogenetic stimulations, extracellular recordings, and chemogenetic inhibition to investigate the role of PC in CrusII, the orolingual region of the cerebellum, in the alleviation of orolingual levodopa-induced dyskinesia (LID). Previous clinical studies found a reduction of LID severity using cerebellar rTMS in PD patients ^17, 19, 20, 59^. However, the precise mechanisms, pathways and cell-types responsible for this beneficial effect remained unknown. In the present study, we first show that CrusII PC opto-stimulations correct, or even prevent, severe orolingual dyskinesia exhibited by chronically levodopa-treated PD mice. These results are the first to demonstrate a direct involvement of PC in the anti-dyskinetic effect of the cerebellum. Strikingly, this beneficial effect led to complete alleviation of orolingual dyskinesia and thus was stronger than observed in patients where rTMS was applied bilaterally over the hemispheres of the cerebellum ^18, 19^. However, the effect of rTMS on cerebellum is not yet well understood: rTMS may only indirectly activate PC and its efficacy may be constrained by the difficulty to target the optimal depths of the cerebellar cortex ^78^. Interestingly, we found that the beneficial effect of cerebellar stimulations in CrusII, which hosts dense projections from the orolingual area ^54^, is mainly observed on the orolingual LID, suggesting a correspondence between the cerebellar somatotopy and functional impact of cerebellar stimulation. Similarly, different subtypes of LID are associated with different patterns of striatal FosB/ΔFosB expression levels, consistent with striatal somatotopy ^38^. The efficacy of cerebellar rTMS in patients should thus strongly depend on the site of stimulation. Cerebellar rTMS have been reported to induce changes within the cerebellar cortex ^79^. However, our work demonstrates a normalization of the neuronal activity in a wide motor network following stimulations, reveals a contribution of the cerebello-thalamo-striatal pathway in mediating the effect of PC stimulations, and shows that it normalizes the expression of striatal FosB/ΔFosB causally linked to LID. Overall, our results indicate that PC stimulations exert long-range effects and act on core mechanisms of LID outside of the cerebellum

Our study further characterizes the alterations occurring in the cerebello-thalamo-cortical and cerebello-thalamo-striatal pathways during dyskinesia. The increased activity of the primary motor cortex in LID observed in our study is consistent with previous findings in rodents and humans ^21, 24^. M1 (and also the subthalamic nucleus which is overactive in PD) reaches DCN and the cerebellar cortex through the pontine nuclei ^48, 80, 81^. This observation contrasts with the decreased activity observed in IN and DN during LID, and may thus reflect compensatory adaptation in the cerebellum ^60^. However, cerebellar nuclei neurons exhibited increased irregularity discharge, and such cerebellar anomalies have been implicated in other motor disorders, such as tremor ^66^, ataxia ^82^ and dystonia ^64, 65^. The irregular activity found in cerebellar nuclei neurons could thus contribute to LID. The PF neurons exhibited a decreased activity as in the cerebellar nuclei; this change could result from a decreased cerebellar entrainment of PF through the cerebello-parafascicular connections ^51, 83,84,85^. Interestingly, our results failed to evidence consistent modulations in the motor thalamus in LID or following cerebellar stimulations. Overall, the changes in activity observed in LID are likely inter-dependent since they were all corrected, or prevented, by the cerebellar stimulations.

The cerebellar stimulations produce alternate periods of silence and increased “rebound” activity in DCN ^86^. Previous evidence demonstrated that this rebound activation is propagated in the forebrain motor network ^71^. The chemogenetic inhibition of the cerebello-parafascicular neurons reduced the beneficial impact of cerebellar stimulations suggesting an important contribution of the cerebello-thalamo-striatal pathway. Examination of the collaterals of these neurons revealed only sparse collaterals to the motor thalamus suggesting a primary role of the thalamostriatal over thalamocortical projections. The striatum is indeed playing a core role in LID generation notably through the overactivity of the direct pathway within the striatum^11, 13, 15^. This overactivity could result from an excessive potentiation at the corticostriatal synapses of direct pathway neurons ^14, 43^. Interestingly, we found that preventive cerebellar stimulations converted the corticostriatal LTP into LTD in direct pathway MSNs in LID mice. This suggests that these stimulations promoted corticostriatal LTD over LTP in the direct pathway and may therefore circumvent the excessive potentiation occurring in LID ^43^. We also found upregulation of striatal FosB/ΔFosB in LID, consistent with previous studies ^35, 36, 38, 40^. Since FosB/ΔFosB overexpression suffices to trigger LID, the normalization of striatal FosB/ΔFosB levels by cerebellar stimulations may explain the suppression of LID. FosB/ΔFosB has been shown to be mainly expressed in D1-expressing striatal neurons ^30, 36, 38^, suggesting that cerebellar stimulations can modulate transcriptional activity in D1-MSNs, probably through projections of IN to PF and the striatum ^85^. These changes of transcriptional activities might also be responsible for the change in corticostriatal plasticity observed in our study. Overall, these results show that our protocol of cerebellar stimulations induce profound changes in the striatal function. Moreover, the persistence of the beneficial effects of this protocol after its end indicates that it recruits a long-term plasticity that could be harnessed for the improvement of LID in PD patients.

Consistent with our finding that cerebellar stimulations may exert a transient therapeutic effect, stimulations of the output pathway of the cerebellum have been recently shown to reduce tremor and ataxia ^66, 87^. Therefore, improving the experimental approaches aimed at stimulating cerebellar Purkinje cells or cerebellar nuclei neurons may benefit to multiple motor disorders. Finally, our work confirms the necessity to study LID as a network disorder involving abnormal signaling between the basal ganglia, cerebral cortex, thalamus and cerebellum ^16, 44^.

## METHODS

### Animals and protocol

L7-ChR2;WT mice ^52^ were used for *in vivo* experiments, L7-ChR2;Drd2-GFP mice were used for *ex vivo* experiments. Animals were housed 1-3 per cage on a standard 12-hour light/dark cycle with *ad libitum* access to water and food. All behavioral manipulations took place during the light phase. All experiments were performed on mice aged 6-9 weeks, of either sex (35-45g), from the Institut de Biologie de l’Ecole Normale Supérieure, Paris, France and in accordance with the recommendations contained in the European Community Council Directives. All animals followed a 9 to 10-weeks experimental protocol. After surgical intervention, mice were carefully monitored during 1-1.5 weeks following a nursing protocol adapted from ^88^ to reduce post-surgery lethality. After 3 weeks, animals received daily intraperitoneal (I.P) injections of L3,4-dihydroxyphenylalanine methyl (L-DOPA, 3 day at 3mg/kg, then 6mg/kg, Sigma-Aldrich) and the peripheral DOPA decarboxylase inhibitor bensezaride hydrochloride (12mg/kg, Sigma-Aldrich) for 6 weeks (0.1mL/10g body weight). Animals were separated into 4 groups: LID_PREV, received daily theta-rhythm cerebellar “preventive” stimulations from the first day of levodopa administration, LID_CORR, received daily theta-rhythm cerebellar “corrective” stimulations after 2 weeks of L-DOPA injections, LID, received daily L-DOPA injections alone, and control SHAM mice, received daily L-DOPA and either no stimulations or preventive or corrective theta-rhythm cerebellar stimulations. Cerebellar stimulations were stopped simultaneously in all the groups and the last two weeks were assessed for long-term anti-dyskinetic effects. Every week, mice were behaviorally monitored (see below).

### Surgical procedures

All surgical procedures were performed at 6-9 weeks of age. After subcutaneous (S.C) administration of buprenorphine (0.06mg/kg), animals were anesthetized with isoflurane (3%) and maintained with 0.5%-1.0% inhaled isoflurane. Mice were placed in a stereotaxic frame (David Kopf Instruments, USA) and pretreated with desipramine (25 mg/kg, I.P) (Sigma-Aldrich). Small holes were drilled over the left medial forebrain bundle (MFB: -1.2 AP, 1.3 ML, -3.75 mm DV) and either over the left oral motor cortex (M1: +2.2 AP, -2.2 ML, -1.5 mm DV), the left parafascicular nucleus (PF: -2.3 AP, -0.75 ML, -3.5 mm DV), and the left ventroanterior-ventrolateral complex of the thalamus (VAL: -1.40 AP, -1.40 ML, -3.50 mm DV) or over the right fastigial nucleus (FN: -6.4 AP, +0.85 ML, -3.25 mm DV), interposed nucleus (IN: -6.4 AP, +1.6 ML, -3.25 mm DV), and dentate nucleus (DN: -6.2 AP, +2.3 ML, -3.3 mm DV). The left MFB was injected with either 1μL of 6-Hydroxydopamine hydrochloride (6-OHDA, (Sigma-Aldrich) 3.2µg/µL free-base in a 0.02% ascorbic acid solution (Sigma-Aldrich)) (for the parkinsonian animals) or 1μL vehicle (ascorbic acid for control animals) at a rate of 0.1μL/min, after which the syringe was left in place for 10min. An additional hole was drilled over the left cerebellum for placement of a skull screw (INOX A2, Bossard) coupled to the ground wire. A final craniectomy centered over the left Crus II (-6.3 to -7.3 mm AP, +3.0 to +4.2 mm ML) was performed, without removing the dura to prevent damage to the cerebellar cortex. A 2.88 mm^2^ LED (SMD chip LED lamp, Kingbright, USA) was then cement to the skull over Crus II. Recording electrodes were slowly lowered through the craniectomy at the wanted coordinates. The ground wire was clipped to the recording board using pins (Small EIB pins, Neuralynx, Dublin, Ireland) and the entire recording device was secured with dental cement (Metabond) and dental acrylic (Pi-Ku-Plast HP 36, monomer and polymer, Bredent, Germany).

All animals were given anti-inflammatory Metacam S.C (Metacam 2mg/mL, Boehringer Ingelheim) for postoperative analgesia and sterile glucose-saline solutions S.C ^88^ (Glucose 5%, Osalia). Parkinsonian animals were closely monitored for 1-1.5 weeks following surgery, if needed mouse cages were kept on a heating pad, animals received several glucose-saline injections daily and were fed a mixt of bledine (Blédina, Danone, France) and concentrated milk.

### Behavior

#### Open field

Animals were monitored in a 38cm diameter open field (Noldus, The Netherlands) once a week for 5 min during 9 weeks. The mice were monitored with a camera (Allied Vision Prosilica GigE GC650, Stemmer Imaging) placed directly above to assess periods of inactivity/activity. The video acquisition was made at 25 Hz frequency and at a resolution of 640 by 480 pixels. DeepLabCut method was used for the analysis of locomotor activity. Seven points of interest were labeled: nose, left ear, right ear, basis of tail, end of tail, red led of the headstage, green led of the headstage. Labels were manually applied to the desired points on 500 frames. The position of the mouse head has been reconstructed from the barycenter of the points: nose, left ear, right ear, red led of the headstage, green led of the headstage weighted by their likelihood. Then, to this head’s trajectory, a cubic smoothing spline fit was applied. For the analysis of locomotion, the periods of activity were isolated from the periods of inactivity by thresholding the speed of the head point at 1cm/s.

#### AIMs assessment

After a 3-weeks pre-L-DOPA phase, daily IP injections of levodopa begun. In rodent models of LID, abnormal involuntary movements (AIMs) are considered the behavioral and mechanistic equivalent of LID in PD patients ^14, 53^. To assess AIMs in our mouse model, a modified scale was used ^53^. Each mouse was placed in a glass cylinder surrounded by 2 mirrors to detect AIMs in every angle and was recorded with a video camera (Allied Vision Prosilica GigE GC650, Stemmer Imaging) for 4 minutes every 20 minutes over a 2-hours period, starting 20 minutes before L-DOPA injection. *Post-hoc* scoring was performed for 2 minutes every 20 minutes. The mice were then evaluated a total of 8 times during the whole recording. Movements were identified as dyskinetic only when they were repetitive, affected the side of the body contralateral to the lesion, and could be clearly distinguished from naturally occurring behaviors such as grooming, sniffing, rearing, and gnawing. Specifically, AIMs were classified into three categories based on their topographic distribution: axial, forelimb, and orolingual. Forelimb dyskinesia are defined as hyperkinetic and/or dystonic movements of the contralateral forelimb on the sagittal or frontal plane. Axial AIMs are considered a twisted posture of the neck and upper body towards the side contralateral to the lesion. Finally, orolingual AIMs are defined as repetitive and empty chewing movements of the jaw, with or without tongue protrusion. Each subtype of AIMs were scored on a severity scale from 0 to 4 where: 0 = absent; 1 = occasional occurrence, less than half of the observation period; 2 = frequent occurrence, more than half of the observation time; 3 = continuous but interrupted by sensory stimuli; and 4 = continuous and not suppressible by sensory distraction ^53^. AIM scores were averaged per time point or per session. Non-parametric Kruskal-Wallis test with pairwise Wilcox test and a Benjamini & Hochberg correction were used to compared the average scores across time and between conditions for peak-dose and average off-period dyskinesia scores.

### Optogenetic stimulations

To stimulate PC on Crus II cerebellar cortex, 3 micro LEDs (1.6×0.6 mm, SMD chip LED lamp, Kingbright, USA) emitting blue light with a dominant 458nm wavelength, were soldered together to cover the full extent of the stimulated area. A small piece of coverslip glass was glued to the bottom of the LEDs to prevent heat brain damage. Two insulated power wires were connecting the LEDs to allow connection with a stimulating cable (New England Wire Technologies, Lisbon) coupled to a LED driver (Universal LED Controler, Mightex). Stimulated mice (LID_PREV, LID_CORR and appropriate controls) received daily train of 20ms theta-rhythm stimulations delivered at 8.33 Hz and at 16mW/mm^2^ irradiance for 2×40seconds separated by a 2 minutes period every day 30 minutes after L-DOPA injection as used in ^19, 89, 90^.

### *In vivo* freely moving chronic recordings

To record cell activity, we used bundles of electrodes consisting of nichrome wire (0.005 inches diameter, Kanthal RO-800) folded and twisted into bundles of 4-8 electrodes. Prior to surgical intervention, the bundles were pinned to an electrode interface board (EIB-16; Neuralynx, Dublin, Ireland) according to the appropriate coordinates of the targeted structures. The microwires of each bundle were connected to the EIB-16 with gold pins (Neuralynx, Dublin, Ireland). The entire recording device was secured by dental cement. The impedance of every electrode was set to 200-500 kΩ using gold-plating (Cyanure-free gold solution, Sifco, France). Electrical signals were acquired using a headstage and amplifier from TDT (RZ2, RV2, Tucker-Davis Technologies, Alachya, FL, USA), filtered, amplified, and recorded on Synapse System (Tucker-Davis Technologies, Alachya, FL, USA). Spike waveform were filtered at 3 Hz low-pass and 8000 Hz high-pass and digitized at 25 kHz. The experimenter manually set a threshold for storage and visualization of electrical events.

During recording sessions, after a 5-minutes open field and a 20-minutes recording baseline, L-DOPA (3-6 mg/kg) was injected IP and pontaneous activity was recorded for 1h30 and stimulated using a LED driver (Universal LED Controler, Mightex), an automatized commutator (ACO32 SYS3-32-CH motorized commutator, Tucker-Davis Technologies, Alachya, FL, USA), and controlled by TTL pulses from our behavioral monitoring system (RV2, Tucker-Davis Technologies, Alachya, FL, USA). At the end of recording sessions, animals were detached and returned to their home cages. Single units were identified offline by manual spike sorting performed on Matlab (Mathworks, Natick, MA, USA) scripts based on k-means clustering on principle component analysis (PCA) of the spike waveforms ^91^. One cluster was considered to represent a single-unit if the unit’s spike waveform was different from other units on the same wire, in 3D PCA space. The unit’s firing activity was analyzed from all structures. No statement is made whether the same cells were recorded across the 9-weeks session. For display purposes, the firing rate of single units was averaged per condition and per group of weeks using boxplots. Boxplots show the median rate in Hz, represented by horizontal bars, over 4 categories of weeks: first boxplot represents the 2^nd^ and 3^rd^ week of the protocol (pre-L-DOPA), second boxplot represents the 4^th^ and 5^th^ weeks when levodopa treatment has started as well as cerebellar stimulations for preventive mice, third boxplot represents the 6^th^ and 7^th^ weeks when corrective mice start receiving cerebellar stimulations, the last boxplot represents the 8^th^ and 9^th^ weeks of the protocol when stimulations stop and long-term effects of cerebellar stimulations are visible. Welch Anova with Games Howell post-hoc test or one-way Anova’s with Tukey post-hoc test were performed based the results of Levene test to compared the averaged firing rate between conditions in DCN whereas linear model ANOVA with the mice randomly distributed and Tukey’s Post-Hoc test were performed to compared the averaged firing rate between conditions in M1, PF, and VAL.

### Neuroanatomical tracing

Mice were injected with 100nL of retrograde AAV-Syn-eGFP (Titer: 1×10^13^ GC/mL, Lot#V16600, Addgene) in PF (N=6). For synaptophysin labeling and axon collaterals, 70nL of AAV8.2-hEF1a-DIO-synaptophysin-GFP (Titer: 2.19×10^13^ GC/mL, Massachusetts General Hospital) together with 50 nL of AAV1.CAG.Flex.tdTomato.WPRE.bGH (Titer: 7.8×10^13^ GC/mL, Lot #CS0923, Upenn Vector) were injected in IN, DN, and FN, in combination with 150nL of retrograde AAV-Cre-EBFP (Titer: 1×10^13^ GC/mL, Lot #V15413, Addgene) injection in PF. Mice were perfused (see below) 15-25 days after injections to allow the expression of the AAVs. Brains were sliced entirely at 90 µm using a vibratome (Leica VT 1000S), and mounted on gelatin-coated slides, dried and then coverslipped with Mowiol (Sigma). Slices were analyzed and imaged using a confocal microscope (SP8, Leica), and images were edited and analyzed using FIJI/ImageJ.

### Chemogenetic experiment

Mice were injected with inhibitory DREADDs pAAV8-hSyn-DIO-hM4DimCherry (Titer: 2.9×10^13^ GC/mL, Vol: 100μL, Lot: v54499; Addgene) in IN, DN, and FN (150nL per nucleus)contralaterally-to-the-lesion, in complementation with 300nL CAV2-Cre-GFP (Titer: 6.4 x 10^12^, dilution 1/10, Plateforme de Vectorologie de Montpellier) viral infusion in ipsilateral-to-the-lesion PF. CAV2-Cre-GFP injections’ surgeries were performed 1 week before the injections of pAAV-hSyn-DIO-hM4DimCherry and the MFB lesion. After 3 weeks, to allow good expression of the viruses and to reproduce our Parkinsonian model, the animals start the levodopa treatment for 6 weeks and the cerebellar stimulations for 4 weeks (LID_PREV). The severity of their orolingual LID was scored 40 minutes after levodopa injection (at the peak dose) and right after cerebellar stimulation. Scores were averaged across the 4 weeks of preventive stimulations. For neuronal modulation of animals expressing DREADDs, Clozapine N-oxide (1.25 mg/kg, Tocris Bioscience) was diluted in saline and injected I.P 10min prior L-DOPA. Control group is injected with saline.

### Perfusion, Immunohistochemistry, Microscopy, and Cell counting

Mice were anesthetized with ketamine/xylazine I.P and transcardially perfused with 4% paraformaldehyde in PBS (Formalin solution, neutral buffer 10%, Sigma-Aldrich). Brains were dissected and post-fixed for 24h in 4% PFA, 24h in 20% sucrose (Merck) and 24h in 30% sucrose. Coronal 20µm sections were cut using a freezing microtome (Leica) and mounted on Superfrost glass slides (Superfrost Plus, Thermo Fisher) for imaging.

For immunohistochemistry, the tissue was blocked with 3% normal donkey serum (NDS, JacksonImmunoResearch) or normal goat serum (NGS, JacksonImmunoResearch) and permeabilized with 0.1% Triton X-100 (Sigma-Aldrich) for 2 hours at room temperature on a shaker. Primary antibodies: Guinea pig anti-TH (Synaptic System, 1:500) and Rabbit anti-FosB (Santa Cruz, 1:100) were added to 1% NDS or NGS and incubated overnight at 4°C on a shaker. Secondary antibodies: donkey anti-Guinea Pig Cy3 (JacksonImmunoResearch, 1:400) and goat anti-Rabbit Alexa 488 (JacksonImmunoResearch, 1: 200) were added in 1% NDS/NGS for 2 hours at room temperature on a shaker. The slices were then washed, incubated with Hoechst (Invitrogen, ThermoFisher scientific, 1:10 000), and mounted onto slides for visualization and imaging. The color reaction was acquired under a dissecting microscope (Leica) and 5, 20, 40 or 64x images were taken. For FosB/ΔFosB quantification, images were taken using confocal microscope (SP8, Leica) with 40x objective. Exposure time were matched between images of the same type. Individual images were stitched together to produce an entire coronal image of both striatum. FIJI/ImageJ software was used to count cells using manual counting. Student t-test was used for statistics. The extend of the dopaminergic lesion was quantified using an optical density analysis on TH staining between the two striatum. Mean density of fluorescence of each striatum was normalized on the mean density of fluorescence of the ipsilateral corpus collasum. Only animals with >50% dopamine depletion were included in this study.

### Ex vivo whole-cell patch-clamp electrophysiology

SHAM, SHAM_PREV, LID, and LID_PREV mice were anaesthetized using isofluorane and their brain removed from the skull. Horizontal striatal slices (270 μm-thick), containing the dorsal lateral striatum, were cut using a VT1000S vibratome (VT1000S, Leica Microsystems, Nussloch, Germany) in ice-cold oxygenated solution (ACSF: 125 mM NaCl, 2.5 mM KCl, 25 mM glucose, 25 mM NaHCO3, 1.25 mM NaH2PO4, 2 mM CaCl2, 1 mM MgCl2, 1 mM pyruvic acid). Slices were then incubated at 32–34°C for 60 minutes before returning to room temperature in holding ACSF. For whole-cell recordings, borosilicate glass pipettes of 4–8 MΩ resistance were filled with a potassium gluconate-based internal solution consisting of (in mM): 122 K-gluconate, 13 KCl, 10 HEPES, 10 phosphocreatine, 4 Mg-ATP, 0.3 Na-GTP, 0.3 EGTA (adjusted to pH 7.35 with KOH, osmolarity 296 ± 3.8 mOsm). Signals were amplified using with EPC10–2 amplifiers (HEKA Elektronik, Lambrecht, Germany). All recordings were performed at 32–34°C, using a temperature control system (Bath-controller V, Luigs&Neumann, Ratingen, Germany) and slices were continuously superfused with extracellular solution at a rate of 2 ml/min. Recordings were sampled at 10 kHz, using the Patchmaster v2×32 program (HEKA Elektronik). D2^+^-MSNs were visualized under direct interference contrast with an upright BX51WI microscope (Olympus, Japan), with a 40x water immersion objective combined with an infra-red filter, a monochrome CCD camera (Roper Scientific, The Netherlands) and a compatible system for analysis of images as well as contrast enhancement. Spike-timing-dependent plasticity (STDP) protocols of stimulations were performed with one concentric bipolar electrode (Phymep, Paris, France; FHC, Bowdoin, ME) placed in the layer 5 of the somatosensory cortex while whole-cell recording MSN in the dorsolateral striatum. Electrical stimulations were monophasic, at constant current (ISO-Flex stimulators, AMPI, Jerusalem, Israel). Currents were adjusted to evoke 100–300 pA EPSCs. STDP protocols consisted of pairings of post-and presynaptic stimulations separated by a specific time interval (∼20 ms); pairings being repeated at 1 Hz. The postsynaptic stimulation of an action potential evoked by a depolarizing current step (30 ms duration) in the recorded MSN preceded the presynaptic cortical stimulation, in a post-pre pairing paradigm. Post–pre pairings was repeated 100 times at 1 Hz (Figure 6a1). Recordings on neurons were made over a period of 10 minutes at baseline, and for at least 40 minutes after the STDP protocols; long-term changes in synaptic efficacy were measured in the last 10 minutes. Experiments were excluded if the mean input resistance (Ri) varied by more than 20% through the experiment. Off-line analysis was performed with Fitmaster (HEKA Elektronik), IGOR Pro 6.0.3 (WaveMetrics, Lake Oswego, OR, USA). Statistical analyses were performed with Prism 7.00 software (San Diego, CA, USA). All results are expressed as mean ± SEM. Statistical significance was assessed in one-sample t tests, unpaired t tests as appropriate, using the indicated significance threshold (p).

## Acknowledgments

This work was supported by Agence Nationale de Recherche to D.P (ANR-12-JSV4-0004 Ceredystim, ANR-16-CE37-0003-02 Amedyst, ANR-19-CE37-0007-01 Multimod) and to C.L. (ANR-17-CE37-0009 Mopla, ANR-17-CE16-0019 Synpredict) and to B.C. (France Parkinson, Labex Memolife) and by the Institut National de la Santé et de la Recherche Médicale (France). The authors declare no competing financial interests. We are grateful to Sabine Meunier for the careful reading of the manuscript. We thank the Imaging Facility at IBENS.

## Author contributions

DP and CL acquired funding; DP and CL conceived and designed all experiments and analysis, except all patch clamp experiments designed by BD and LV; BC, DP, CL wrote the manuscript; BC, JLF, EP, AC, TT, CL analyzed the data; BC, JLF, EP, AC, FM, CMH, SP, DP performed the experiments. All authors interpreted results, revised the final manuscript, and approved the final manuscript.

## Data, code and materials availability

All data are available in the main text or in the supplementary materials. The code for electrophysiological analysis is available from the corresponding author upon reasonable request.

## SUPPLEMENTARY FIGURE LEGENDS

**Supplementary Fig. S1.**
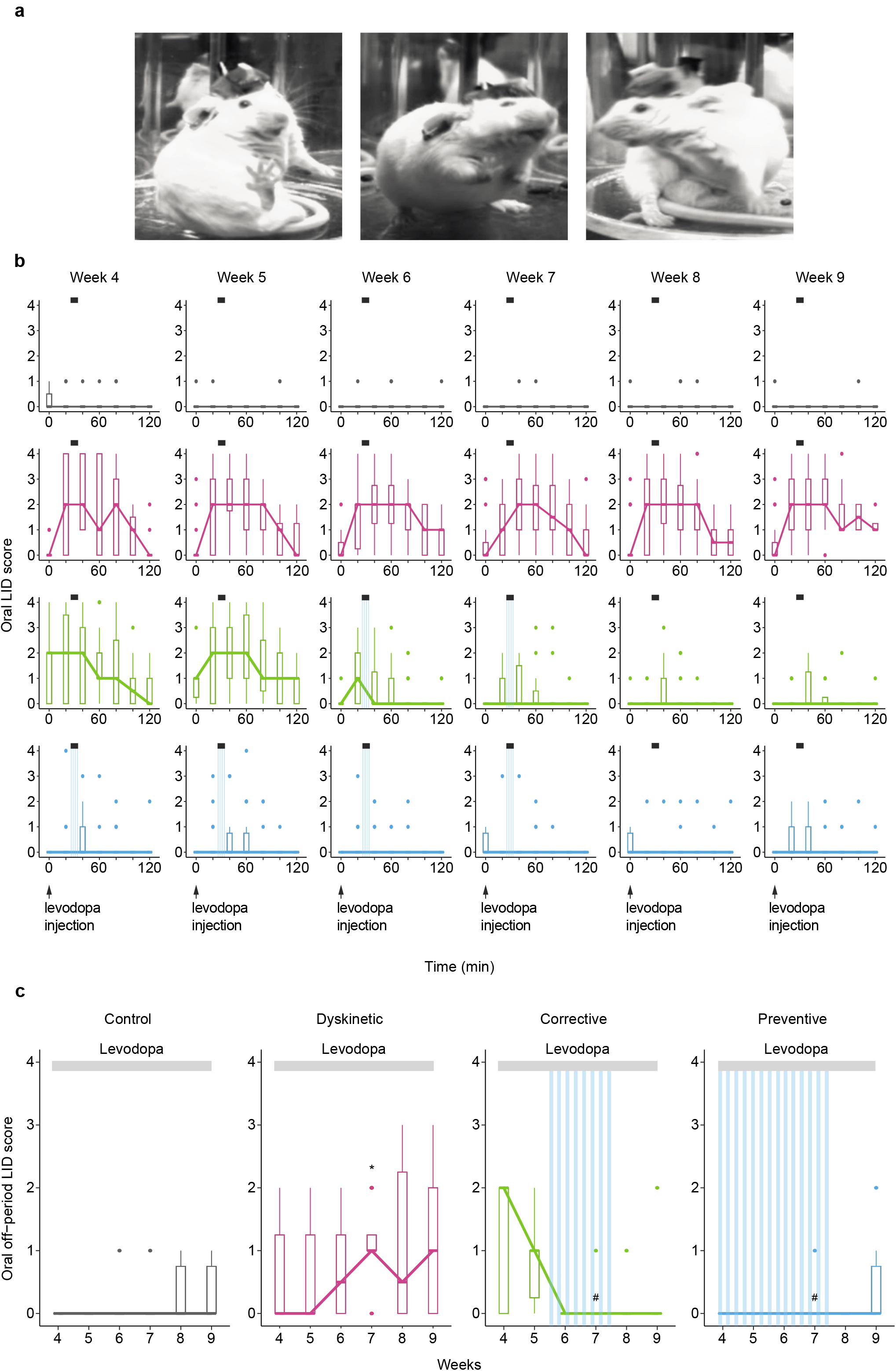
Optogenetic stimulations of CrusII Purkinje cells are sufficient to both reduce and prevent severe orolingual peak-dose dyskinesia. Related to Fig. 1. **a** Examples of orolingual peak-dose levodopa-induced dyskinesia in L7-ChR2-YFP LID parkinsonian mice chronically treated with levodopa. **b** Boxplot showing the average oral LID scores measured over 7 time points starting from the time of levodopa injection (black arrows) across the 6 weeks of treatment in SHAM (grey, N=17), LID (magenta, N=19), LID_CORR receiving 2 weeks of Purkinje cell stimulations (green, N=24), and LID_PREV mice receiving 4 weeks of Purkinje cell stimulations (blue, N=18). Boxplots represents the lower and the upper quartiles and horizontal bars in boxplots represent median score. Light grey lines: time of levodopa peak-dose effect. Striped blue lines: time of theta-burst stimulations. **c** Boxplot showing the average oral “off-period” LID scores measured 20 minutes before levodopa injection across the 6 weeks of treatment in SHAM (grey, N=8), LID (magenta, N=6), LID_CORR (green, N=6), and LID_PREV mice (blue, N=4). Boxplots represents the lower and the upper quartiles and horizontal bars in boxplots represent median score. Light grey lines: 6 weeks of levodopa treatment. Stripped blue lines: weeks of Purkinje cell stimulations. See also Table S1.

**Supplementary Fig. S2.**
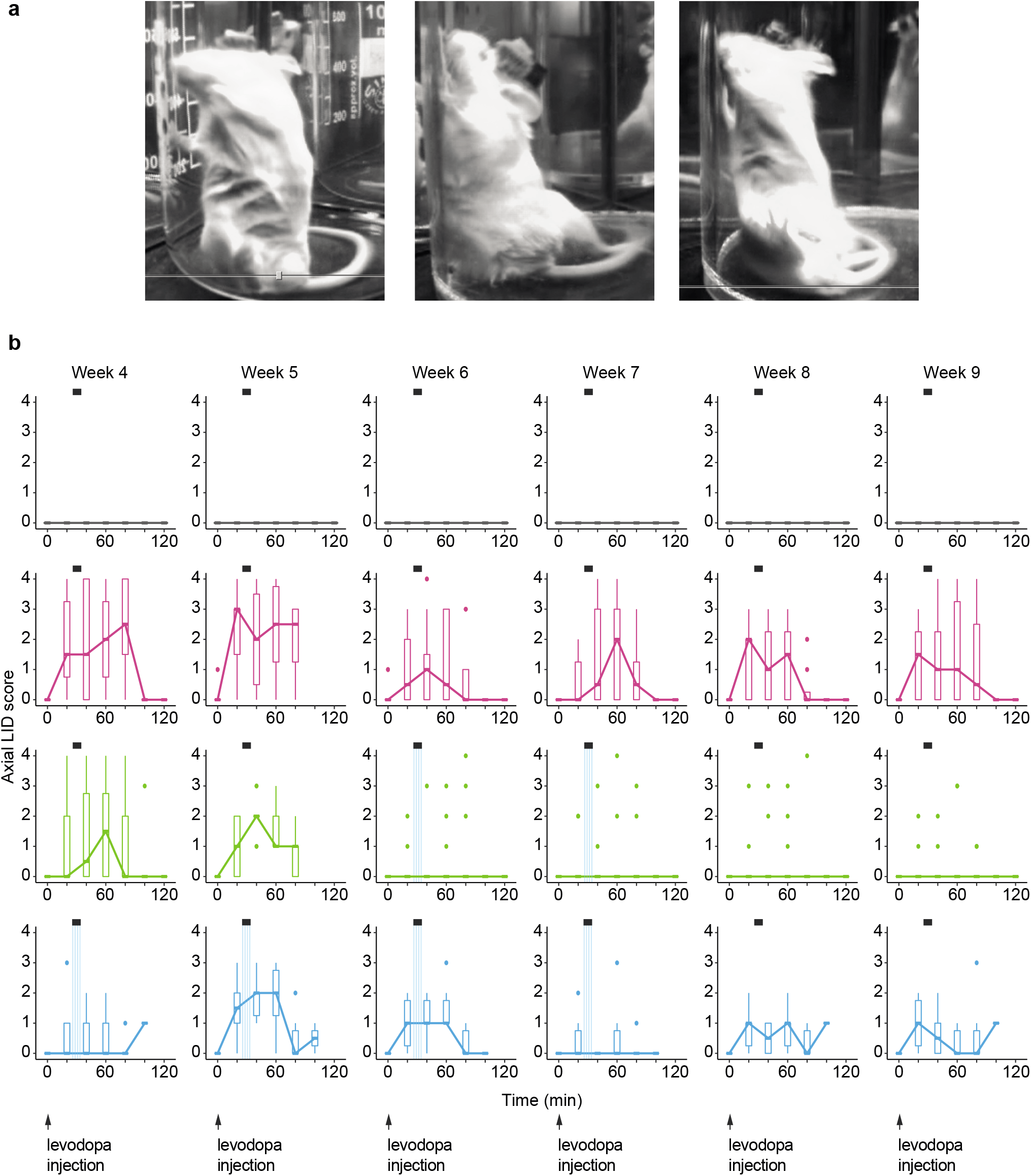
Optogenetic stimulations of CrusII Purkinje cells are not sufficient to completely reduce and prevent severe axial peak-dose dyskinesia. Related to Fig. 1. **a** Examples of axial peak-dose levodopa-induced dyskinesia in L7-ChR2-YFP LID parkinsonian mice chronically treated with levodopa. **b** Boxplot showing the average axial LID scores measured over 7 time points starting from the time of levodopa injection (black arrows) across the 6 weeks of treatment in SHAM (grey, N=12), LID (magenta, N=8), LID_CORR receiving 2 weeks of Purkinje cell stimulations (green, N=14), and LID_PREV mice receiving 4 weeks of Purkinje cell stimulations (blue, N=6). Boxplots represents the lower and the upper quartiles and horizontal bars in boxplots represent median score. Light grey lines: time of levodopa peak-dose effect. Striped blue lines: time of theta-burst stimulations.

**Supplementary Fig. S3.**
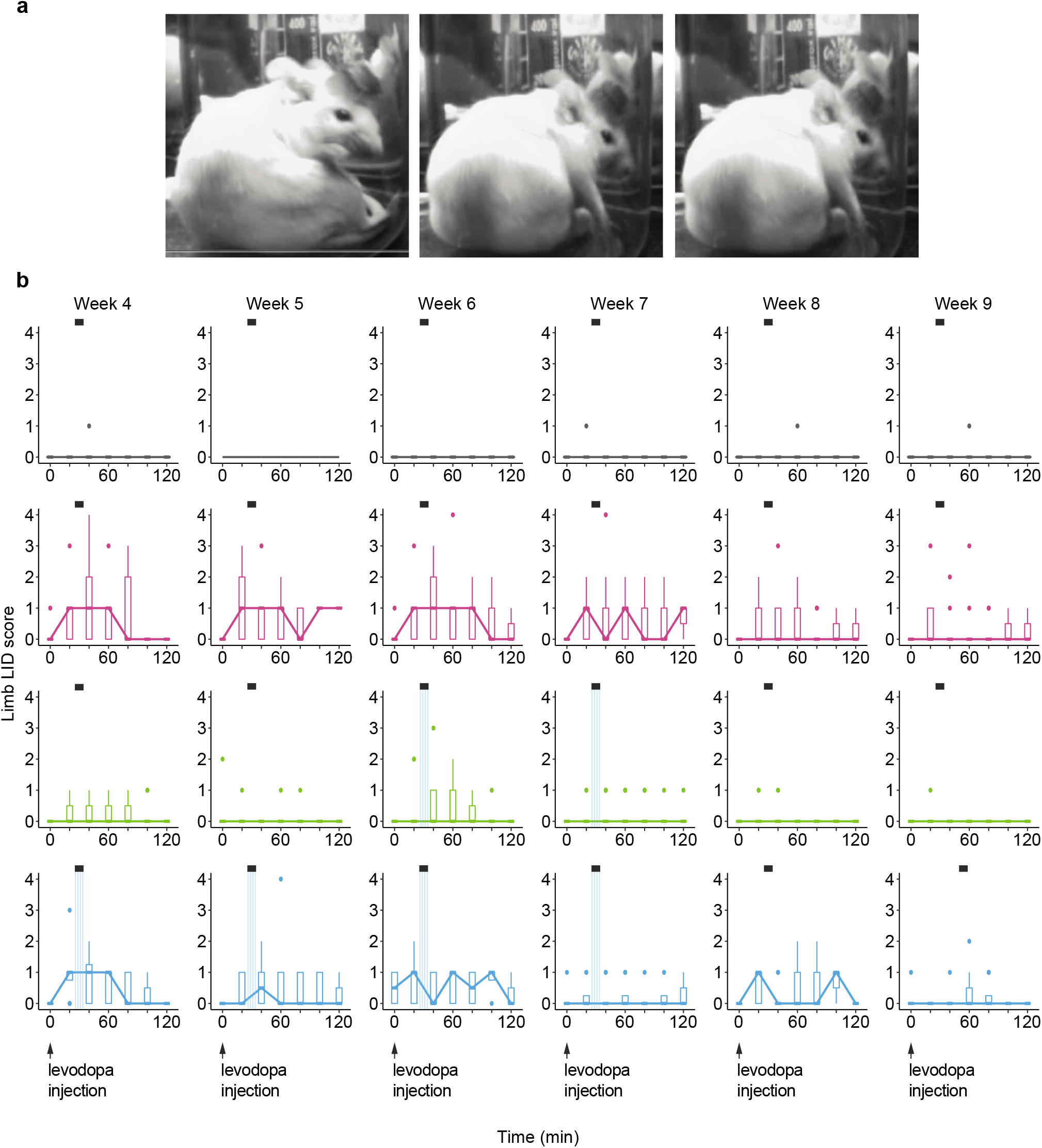
Optogenetic stimulations of CrusII Purkinje cells are not sufficient to completely reduce and prevent severe limb peak-dose dyskinesia. Related to Fig. 1. **a** Examples of limb peak-dose levodopa-induced dyskinesia in L7-ChR2-YFP LID parkinsonian mice chronically treated with levodopa. **b** Boxplot showing the average limb LID scores measured over 7 time points starting from the time of levodopa injection (black arrows) across the 6 weeks of treatment in SHAM (grey, N=14), LID (magenta, N=9), LID_CORR receiving 2 weeks of Purkinje cell stimulations (green, N=15), and LID_PREV mice receiving 4 weeks of Purkinje cell stimulations (blue, N=9). Boxplots represents the lower and the upper quartiles and horizontal bars in boxplots represent median score. Light grey lines: time of levodopa peak-dose effect. Striped blue lines: time of theta-burst stimulations.

**Supplementary Fig. S4.**
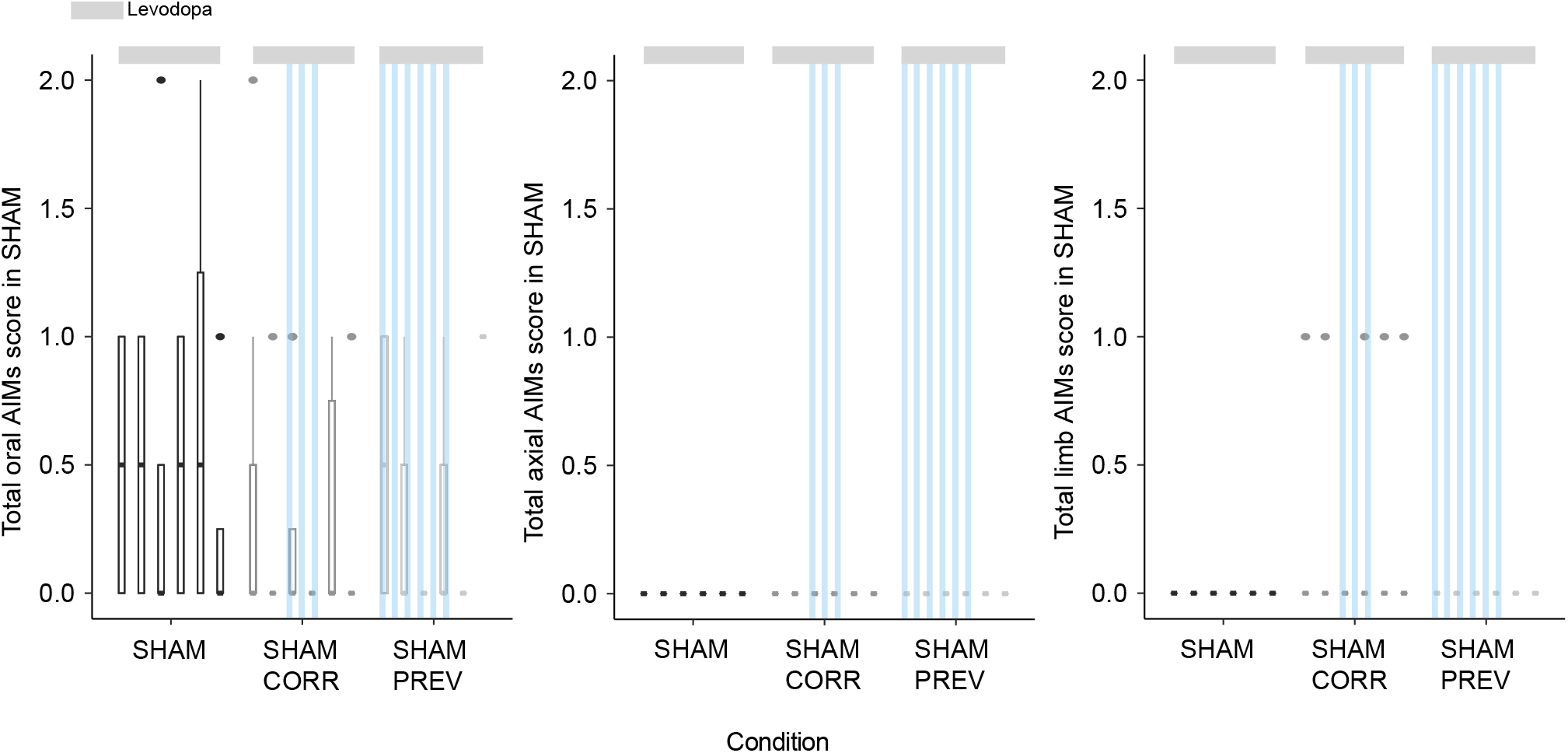
Optogenetic stimulations of CrusII Purkinje cells does not affect severe peak-dose oral, axial and limb dyskinesia in SHAM. Related to Fig. 1. Boxplot showing the sum of oral (left), axial (middle), and limb (right) LID scores measured over 8 time points starting 20 minutes before levodopa injection to 120 minutes after across the 6 weeks of treatment in SHAM mice treated with levodopa (dark grey, SHAM, N=4), SHAM mice treated with levodopa and receiving 2 weeks of Purkinje cells stimulations (grey, SHAM_CORR, N=5), SHAM mice treated with levodopa and receiving 4 weeks of Purkinje cells stimulations (light grey, SHAM_PREV, N=3). Horizontal bars in boxplots represent median score. Light grey lines: 6 weeks of levodopa treatment. Stripped blue lines: time of theta-burst stimulations. Boxplots represents the lower and the upper quartiles. Non-parametric Kruskal-Wallis test with pairwise Wilcox test and a Benjamini & Hochberg correction. ***p < 0.001; **p < 0.01; *p < 0.05; ns: p > 0.5. See also Table S8.

**Supplementary Fig. S5.**
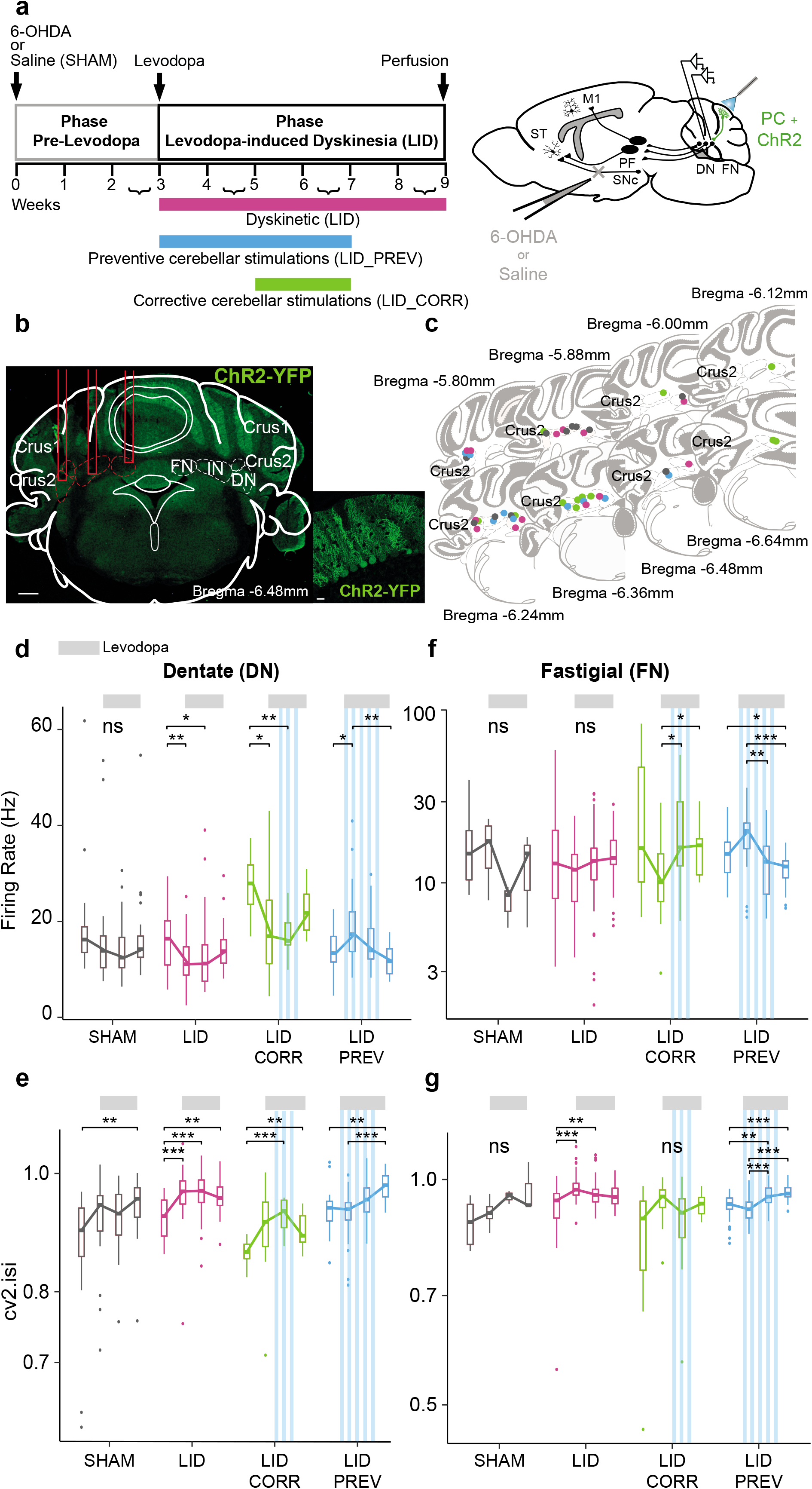
Purkinje cell stimulations do not completely normalize the aberrant activity in the dentate and fastigial cerebellar nuclei. Related to Fig. 2. **a** *Left*: Experimental timeline. *Right*: Schematic of electrode implantation in the dentate (DN) and fastigial nuclei (FN), ChR2-YFP expression in the Purkinje cells (PC+ChR2, green) and injection site with 6-OHDA or saline in sagittal mouse brain. ST: Striatum; SNc: substantia nigra *pars compacta*; M1: Primary motor cortex; PF: Parafascicular nucleus of the thalamus. **b** Coronal section from L7-ChR2-YFP mouse showing electrode’s trajectory (red lines). Dotted red lines: IN, FN, and DN. Green: ChR2-YFP expression. Scale bars: 0.5 mm (left); 20µm (right). Crus1: Crus1 of the ansiform lobule, Crus2: Crus2 of the ansiform lobule. **c** Schematic of verified recording sites in IN, FN, and DN (grey = SHAM; magenta = LID; green = LID_CORR; blue = LID_PREV). Crus2: Crus2 of the ansiform lobule. **d** Firing rate (Hz) across 9 weeks in DN. Boxplots show the median rate (horizontal bars), over 4 categories of weeks. First boxplot: 2^nd^ and 3^rd^ week of the protocol, second boxplot: 4^th^ and 5^th^ weeks when levodopa treatment started, third boxplot: 6^th^ and 7^th^ weeks, last boxplot: 8^th^ and 9^th^ weeks when stimulations stopped. Grey = SHAM (N=3); Magenta = LID (N=3); Green = LID_CORR (N=3); Blue = LID_PREV (N=3). Light grey lines: 6 weeks of levodopa (3 boxplots). Stripped blue lines: weeks of theta-burst PC stimulations. **e** Coefficient of variation 2 (cv2.isi) across 9 weeks in DN. The order of boxplots is identical to panel **d**. Grey = SHAM (N=3); Magenta = LID (N=3); Green = LID_CORR (N=3); Blue = LID_PREV (N=3). Light grey lines: 6 weeks of levodopa (3 boxplots). Stripped blue lines: weeks of theta-burst PC stimulations. **f** Firing rate (Hz) across 9 weeks in FN. The order of boxplots is identical to panel **d**. Grey = SHAM (N=3); Magenta = LID (N=6); Green = LID_CORR (N=4); Blue = LID_PREV (N=3). Light grey lines: 6 weeks of levodopa (3 boxplots). Stripped blue lines: weeks of theta-burst PC stimulations. **g** Coefficient of variation 2 (cv2.isi) across 9 weeks in FN. The order of boxplots is identical to panel **d**. Grey = SHAM (N=3); Magenta = LID (N=6); Green = LID_CORR (N=4); Blue = LID_PREV (N=3). Light grey lines: 6 weeks of levodopa (3 boxplots). Stripped blue lines: weeks of theta-burst PC stimulations. Boxplots represents the lower and the upper quartiles. Welch Anova with Games Howell post-hoc test and one-way Anova’s with Tukey post-hoc test based on Levene test. ***p < 0.001; **p < 0.01; *p < 0.05; ns: p > 0.5. See also Tables S2, S3, and S4.

### Influence of locomotion on DCN discharge

As DCN activity is modulated by locomotion in normal animals (Sarnaik and Raman, 2018), we quantified the locomotor activity using DeepLabCut (Mathis et al., 2018) (Supplementary Figure 6) and tested if the activity observed in DCN correlated with a change in locomotor activity in control animals (Supplementary Figure 7). No differences were observed in the DCN during levodopa treatment in control animals, whether the animals were moving or not (Supplementary Figure 7).

### Increased cv2.isi does not simply reflect increased bursting

Since greater cv2.isi in LID mice may reflect irregular burst firing under levodopa treatment in DCN neurons, we analyzed the burst rate in periods of locomotor activity and inactivity. Surprisingly, the burst rate of dyskinetic mice decreased during levodopa treatment in periods of activity in the IN, DN, and FN (Supplementary Figure 8a). However, the burst rate of LID mice in periods of locomotor inactivity increased during levodopa treatment in the DN and FN (Supplementary Figure 8b).

PC stimulations during 4 weeks prevented the decreased of the burst rate in the IN during periods of locomotor activity (Supplementary Figure 8a**, left panel**). Changes, although less consistent, were also observed as a function of the motor state in the DN and FN (Supplementary Figure 8a**, middle and right panels)**. Only mild differences were observed in the burst rate in the IN, DN or FN in the different groups during periods of locomotor inactivity (Supplementary Figure 8b).

---

**Supplementary Fig. S6.**
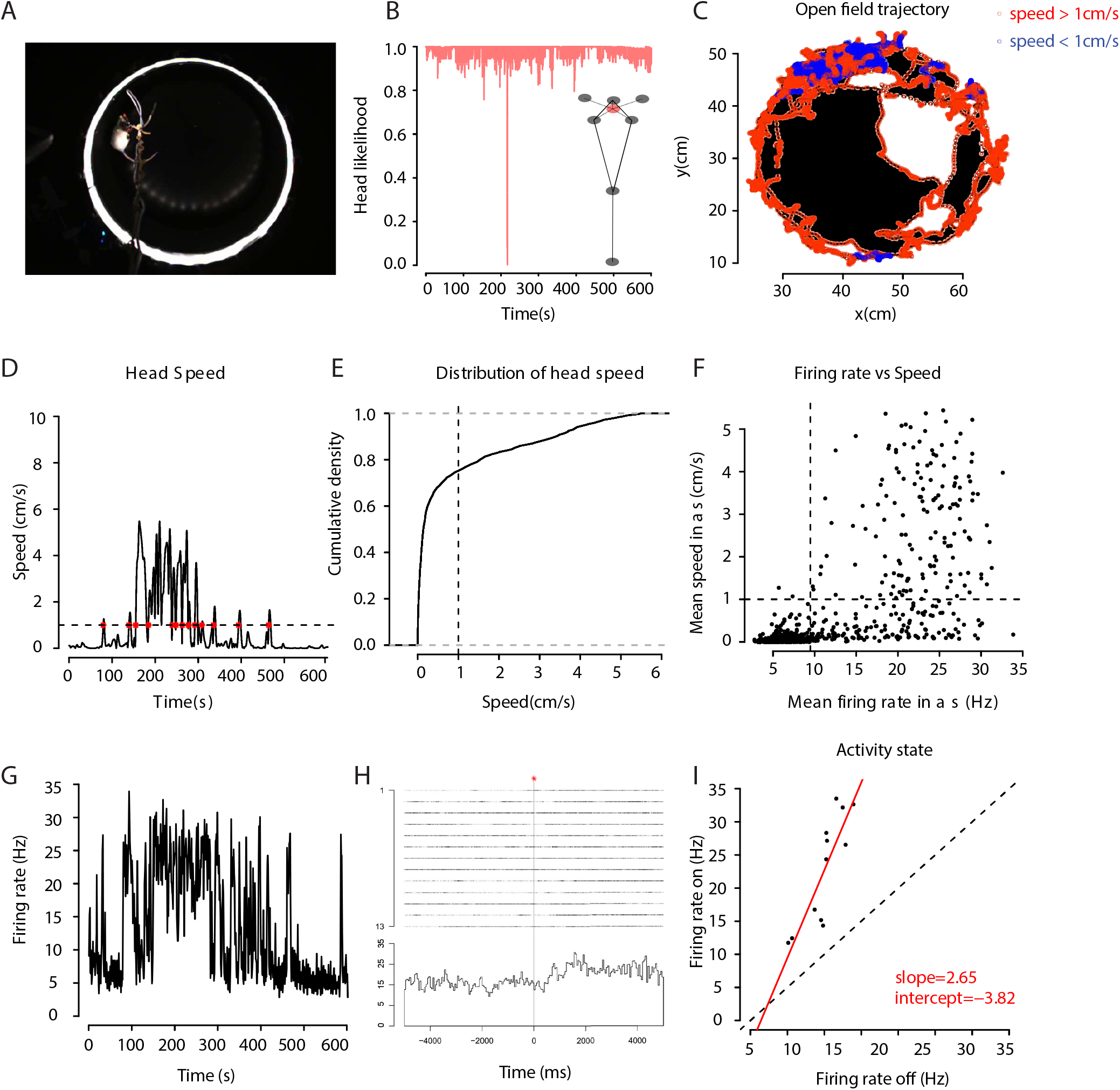
Modulation of the firing rate of the deep cerebellar nuclei induced by locomotor activity. Related to Fig. 2. **a** Image of the tracking points of interest, represented by different colors (tip of the tail: dark blue; base of the tail: light blue; right ear: dark green; left ear: light green; nose: yellow; right led: dark red; left led: light red) during a recording session in the openfield using Deeplabcut. **b** Evolution of the probability of the head point during the recording. Diagram of the reconstruction of the head point (light red) from the barycenter weighted by the probabilities of the points: nose; right ear; left ear; right led; left led. **c** Path of the head’s point as a function of the movement speed. The movement speed is discretized in “off” and “on” periods which are represented by blue and red circles, respectively when the speed is lower and higher than the threshold of 1cm/s. **d** Evolution of the speed of the head’s point. The moments when the speed of the head’s point crosses the threshold of 1cm/s (dotted line) are reported by a red star. **e** Cumulative distribution of the speed of the head’s point. Threshold of 1cm/s: vertical dotted line. **f** Mean firing rate (Hz) of the deep cerebellar nuclei (DCN) activity as a function of the average velocity defined per second (cm/s). Mean firing rate: vertical dotted line, velocity threshold: horizontal dotted line. **g** Evolution of the average firing rate (Hz) of the DCN activity over time. **h** Raster plot of mean DCN activity around the transitions from “off” to “on” state of locomotor activity (red stars in D). **i** Relation of the firing rate (Hz) between the locomotor “on” and “off” periods defined per cell (n=12). Dotted line (y=x), straight red line of linear regression of the firing rate of the cells during the “on” periods as a function of the “off” periods.

**Supplementary Fig. S7.**
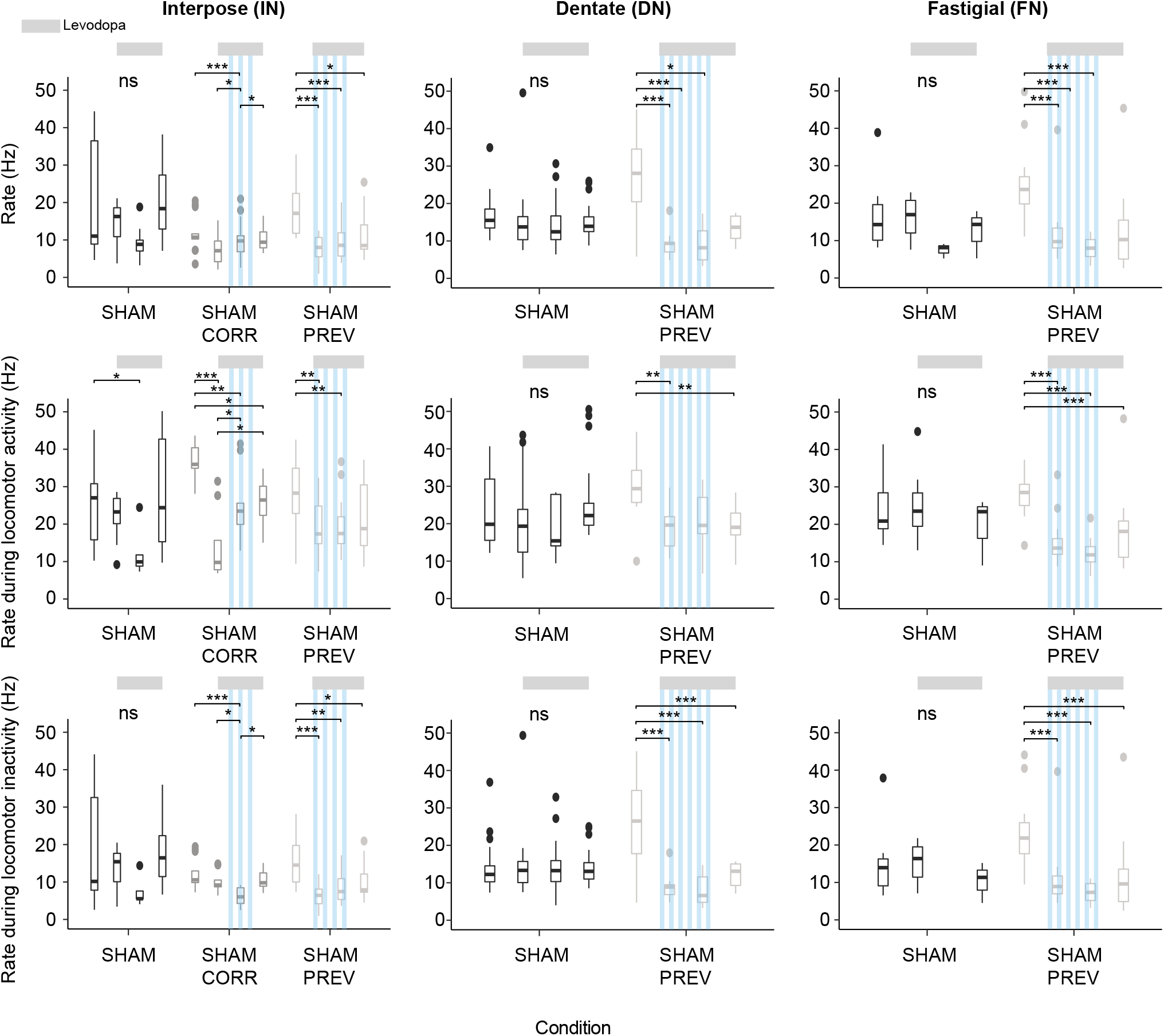
Locomotor activity does not impact the firing rate of the three deep cerebellar nuclei in SHAM. Related to Fig. 2. *Top*: Firing rate (Hz) over 9 weeks in the three deep cerebellar nuclei (DCN) in all control mice: the interposed nucleus (IN, left), the dentate nucleus (DN, middle), and the fastigial nucleus (FN, right). *Middle*: Firing rate (Hz) during periods of locomotor activity (“on”) over 9 weeks in DCN in all control mice: IN (left), DN (middle), and FN (right). *Bottom*: Firing rate (Hz) during periods of locomotor inactivity (“off”) over 9 weeks in DCN in all control mice: IN (left), DN (middle), and FN (right). Boxplots represents the lower and the upper quartiles and show the median rate (horizontal bars), over 4 categories of weeks. First boxplot: 2^nd^ and 3^rd^ week of the protocol, second boxplot: 4^th^ and 5^th^ weeks when levodopa treatment started as well as cerebellar stimulations for preventive sham mice, third boxplot: 6^th^ and 7^th^ weeks when corrective sham mice start receiving cerebellar stimulations, last boxplot: 8^th^ and 9^th^ weeks of the protocol when stimulations stop and long-term effects of cerebellar stimulations are visible. Control mice only treated with levodopa represented in dark grey (SHAM, IN: N=1, FN: N=1, DN: N=2); control mice treated with levodopa and receiving 2 weeks of Purkinje cell stimulations represented in grey (SHAM_CORR, IN: N=2); control mice treated with levodopa and receiving 4 weeks of Purkinje cell stimulations represented in grey (SHAM_PREV, IN: N=2, FN: N=2, DN: N=1). Light grey lines: 6 weeks of levodopa (3 boxplots). Stripped blue lines: weeks of theta-burst Purkinje cell stimulations. Welch Anova with Games Howell post-hoc test and one-way Anova’s with Tukey post-hoc test based on Levene test. ***p < 0.001; **p < 0.01; *p < 0.05; ns: p > 0.5. See also Table S9.

**Supplementary Fig. S8.**
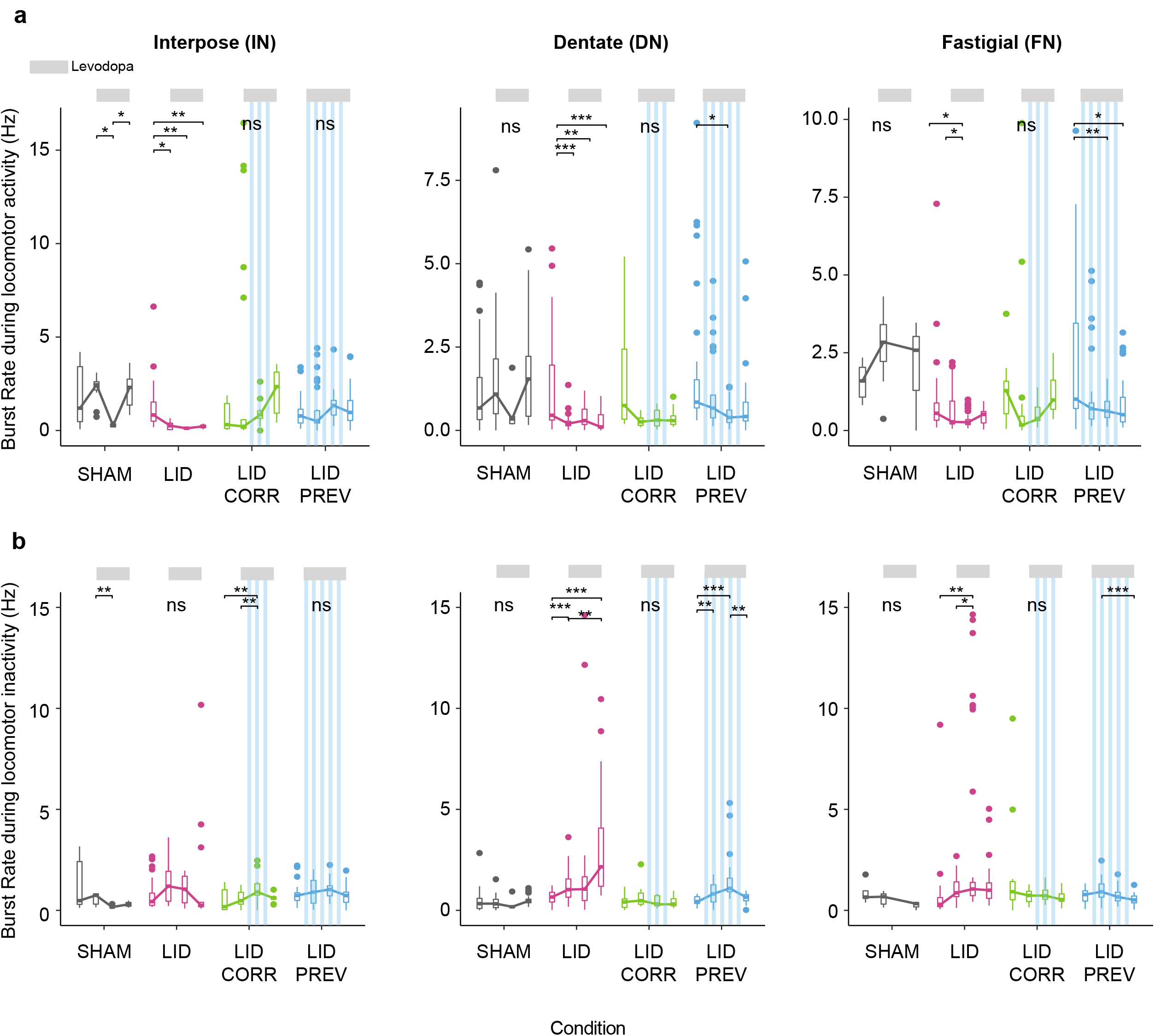
In dyskinetic mice, the burst rate decreases during periods of activity and increases during periods of inactivity. These effects are prevented by cerebellar stimulations in the interposed nucleus. Related to Fig. 2. **a** Burst rate (Hz) during periods of locomotor activity (“on”) over 9 weeks in the interposed nucleus (IN, left), the dentate nucleus (DN, middle), and the fastigial nucleus (FN, right). Boxplots show the median burst rate (horizontal bars), over 4 categories of weeks. First boxplot: 2^nd^ and 3^rd^ week of the protocol, second boxplot: 4^th^ and 5^th^ weeks when levodopa treatment started, third boxplot: 6^th^ and 7^th^, last boxplot: 8^th^ and 9^th^ weeks when stimulations stopped. Grey = SHAM (IN: N=4, DN: N=3, FN: N=3); Magenta = LID (IN: N=3, DN: N=4, FN: N=6); Green = LID_CORR (IN: N=4, DN: N=3, FN: N=4); Blue = LID_PREV (IN: N=3, DN: N=3, FN: N=3). Light grey lines: 6 weeks of levodopa (3 boxplots). Stripped blue lines: weeks of theta-burst Purkinje cell stimulations. **b** Burst rate (Hz) during periods of locomotor inactivity (“off”) over 9 weeks in IN (left), DN (middle), and FN (right). Same order of boxplot as panel A. Grey = SHAM (IN: N=4, DN: N=3, FN: N=3); Magenta = LID (IN: N=3, DN: N=4, FN: N=6); Green = LID_CORR (IN: N=4, DN: N=3, FN: N=4); Blue = LID_PREV (IN: N=3, DN: N=3, FN: N=3). Light grey lines: 6 weeks of levodopa (3 boxplots). Stripped blue lines: weeks of theta-burst Purkinje cell stimulations. Boxplots represents the lower and the upper quartiles. Welch Anova with Games Howell post-hoc test and one-way Anova’s with Tukey post-hoc test based on Levene test. ***p < 0.001; **p < 0.01; *p < 0.05; ns: p > 0.5. See also Table S10 and S11.

**Supplementary Fig. S9.**
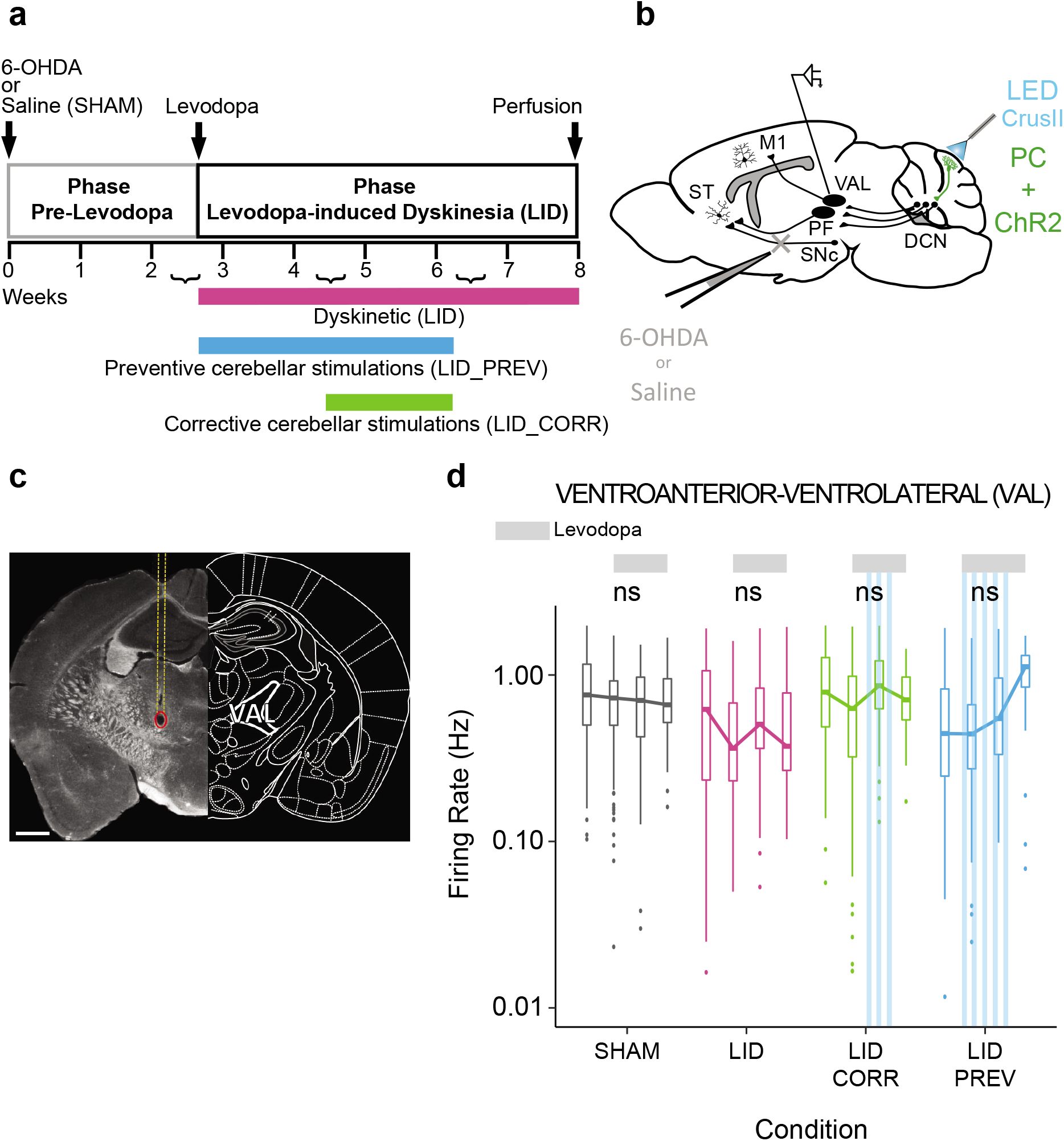
Neither levodopa nor Purkinje cell stimulations affect the firing rate in the ventroanterior-ventrolateral complex of the thalamus. Related to Fig. 3. **a** Experimental timeline. Dyskinetic mice (LID, red): 5 weeks of levodopa treatment. Preventive mice LID_PREV, blue): 5 weeks of levodopa treatment + 4 weeks of cerebellar stimulations. Corrective mice (LID_CORR, green): 5 weeks of levodopa treatment + 2 weeks of cerebellar stimulations. **b** Schematic of electrode implantation in ventroanterior-ventrolateral complex of the thalamus (VAL), ChR2-YFP expression in the Purkinje cells (PC+ChR2, green) and injection site with 6-OHDA or saline in sagittal mouse brain. ST: Striatum; SNc: substantia nigra *pars compacta*; M1: Primary motor cortex, PF: Parafascicular nucleus of the thalamus, DCN: deep cerebellar nuclei, CrusII: Crus2 of the ansiform lobule. **c** Coronal section from L7-ChR2-YFP mouse showing the electrode’s trajectory (yellow dotted line) and the electrolytic lesion (red circle) at the recording site in VAL. Scale bar: 0.5 mm. **d** Firing rate (Hz) across 9 weeks in VAL. Boxplots show the median rate (horizontal bars), over 4 categories of weeks. First boxplot: 2^nd^ and 3^rd^ week of the protocol, second boxplot: 4^th^ and 5^th^ weeks when levodopa treatment started, third boxplot: 6^th^ and 7^th^ weeks, last boxplot represents the 8^th^ week when stimulations stopped. Grey = SHAM (N=5); Magenta = LID (N=4); Green = LID_CORR (N=6); Blue = LID_PREV (N=8). Light grey lines: 6 weeks of levodopa (3 boxplots). Stripped blue lines: weeks of theta-burst PC stimulations. Boxplots represents the lower and the upper quartiles. One-way Anova with Tukey HSD post-hoc test. ***p < 0.001; **p < 0.01; *p < 0.05; ns: p > 0.5. See also Table S6.

**Table S1:** Precision measures, exact p-values, and replicate data relevant to Figure 1, and Supplementary Figure 1c.

| Fig | Param | Week | Group1 | Group2 | Mean1 | Mean2 | n1 | n2 | Sum | Adjusted<br>p Value |
| --- | --- | --- | --- | --- | --- | --- | --- | --- | --- | --- |
| Fig. 1c | 6OHDA<br>lesion | / | 6-OHDA | SHAM | 80.93 | 11.05 | 40 | 17 | *** | < 0.001 |
| Fig. 1e | Oral | 4 | LID | SHAM | 8.84 | 0.42 | 19 | 19 | *** | < 0.001 |
|  |  |  | LID_CORR | SHAM | 7.96 | 0.42 | 23 | 19 | *** | < 0.001 |
|  |  |  | LID_PREV | SHAM | 1.59 | 0.42 | 17 | 19 | * | 0.035 |
|  |  |  | LID_CORR | LID | 7.96 | 8.84 | 23 | 19 | ns | 0.711 |
|  |  |  | LID_PREV | LID | 1.59 | 8.84 | 17 | 19 | ### | < 0.001 |
|  |  |  | LID_PREV | LID_CORR | 1.59 | 7.96 | 17 | 23 | &&& | < 0.001 |
| Fig. 1e | Oral | 5 | LID | SHAM | 7.79 | 0.23 | 19 | 17 | *** | < 0.001 |
|  |  |  | LID_CORR | SHAM | 9.20 | 0.23 | 15 | 17 | *** | < 0.001 |
|  |  |  | LID_PREV | SHAM | 1.83 | 0.23 | 18 | 17 | ** | 0.002 |
|  |  |  | LID_CORR | LID | 9.20 | 7.79 | 15 | 19 | ns | 0.425 |
|  |  |  | LID_PREV | LID | 1.83 | 7.79 | 18 | 19 | ### | < 0.001 |
|  |  |  | LID_PREV | LID_CORR | 1.83 | 9.20 | 18 | 15 | &&& | < 0.001 |
| Fig. 1e | Oral | 6 | LID | SHAM | 8.31 | 0.23 | 19 | 19 | *** | < 0.001 |
|  |  |  | LID_CORR | SHAM | 2.71 | 0.23 | 24 | 19 | *** | < 0.001 |
|  |  |  | LID_PREV | SHAM | 0.94 | 0.23 | 18 | 19 | * | 0.048 |
|  |  |  | LID_CORR | LID | 2.71 | 8.31 | 24 | 19 | ### | < 0.001 |
|  |  |  | LID_PREV | LID | 0.94 | 8.31 | 18 | 19 | ### | < 0.001 |
|  |  |  | LID_PREV | LID_CORR | 0.94 | 2.71 | 18 | 24 | && | 0.002 |
| Fig. 1e | Oral | 7 | LID | SHAM | 8.37 | 0.20 | 19 | 15 | *** | < 0.001 |
|  |  |  | LID_CORR | SHAM | 1.87 | 0.20 | 23 | 15 | *** | < 0.001 |
|  |  |  | LID_PREV | SHAM | 0.78 | 0.20 | 18 | 15 | ns | 0.127 |
|  |  |  | LID_CORR | LID | 1.87 | 8.37 | 23 | 19 | ### | < 0.001 |
|  |  |  | LID_PREV | LID | 0.78 | 8.37 | 18 | 19 | ### | < 0.001 |
|  |  |  | LID_PREV | LID_CORR | 0.78 | 1.87 | 18 | 23 | & | 0.010 |
| Fig. 1e | Oral | 8 | LID | SHAM | 8.74 | 0.35 | 19 | 17 | *** | < 0.001 |
|  |  |  | LID_CORR | SHAM | 0.91 | 0.35 | 23 | 17 | ns | 0.16 |
|  |  |  | LID_PREV | SHAM | 0.78 | 0.35 | 18 | 17 | ns | 0.43 |
|  |  |  | LID_CORR | LID | 0.91 | 8.74 | 23 | 19 | ### | < 0.001 |
|  |  |  | LID_PREV | LID | 0.78 | 8.74 | 18 | 19 | ### | < 0.001 |
|  |  |  | LID_PREV | LID_CORR | 0.78 | 0.91 | 18 | 23 | ns | 0.43 |
| Fig. 1e | Oral | 9 | LID | SHAM | 6.42 | 0.36 | 19 | 11 | *** | < 0.001 |
|  |  |  | LID_CORR | SHAM | 1.31 | 0.36 | 16 | 11 | ns | 0.053 |
|  |  |  | LID_PREV | SHAM | 1.92 | 0.36 | 13 | 11 | ** | 0.004 |
|  |  |  | LID_CORR | LID | 1.31 | 6.42 | 16 | 19 | ### | < 0.001 |
|  |  |  | LID_PREV | LID | 1.92 | 6.42 | 13 | 19 | ### | < 0.001 |
|  |  |  | LID_PREV | LID_CORR | 1.92 | 1.31 | 13 | 16 | ns | 0.219 |
| Fig. 1f | Axial | 4 | LID | SHAM | 8.12 | 0 | 8 | 12 | *** | <0.001 |
|  |  |  | LID_CORR | SHAM | 5.07 | 0 | 14 | 12 | *** | <0.001 |
|  |  |  | LID_PREV | SHAM | 2.40 | 0 | 5 | 12 | *** | <0.001 |
|  |  |  | LID_CORR | LID | 5.07 | 8.12 | 14 | 8 | ns | 0.077 |
|  |  |  | LID_PREV | LID | 2.40 | 8.12 | 5 | 8 | # | 0.018 |
|  |  |  | LID_PREV | LID_CORR | 2.40 | 5.07 | 5 | 14 | ns | 0.219 |
| Fig. 1f | Axial | 5 | LID | SHAM | 9.00 | 0 | 6 | 9 | *** | <0.001 |
|  |  |  | LID_CORR | SHAM | 5.40 | 0 | 5 | 9 | *** | <0.001 |
|  |  |  | LID_PREV | SHAM | 5.83 | 0 | 6 | 9 | *** | <0.001 |
|  |  |  | LID_CORR | LID | 5.40 | 9.00 | 5 | 6 | ns | 0.190 |
|  |  |  | LID_PREV | LID | 5.83 | 9.00 | 6 | 6 | ns | 0.190 |
|  |  |  | LID_PREV | LID_CORR | 5.83 | 5.40 | 6 | 5 | ns | 0.980 |
| Fig. 1f | Axial | 6 | LID | SHAM | 4.25 | 0 | 8 | 11 | *** | <0.001 |
|  |  |  | LID_CORR | SHAM | 1.50 | 0 | 14 | 11 | ** | 0.006 |
|  |  |  | LID_PREV | SHAM | 4.00 | 0 | 6 | 11 | *** | <0.001 |
|  |  |  | LID_CORR | LID | 1.50 | 4.25 | 14 | 8 | ## | 0.002 |
|  |  |  | LID_PREV | LID | 4.00 | 4.25 | 6 | 8 | ns | 0.610 |
|  |  |  | LID_PREV | LID_CORR | 4.00 | 1.50 | 6 | 14 | &&& | <0.001 |
| Fig. 1f | Axial | 7 | LID | SHAM | 4.62 | 0 | 8 | 11 | *** | <0.001 |
|  |  |  | LID_CORR | SHAM | 1.36 | 0 | 14 | 11 | * | 0.014 |
|  |  |  | LID_PREV | SHAM | 1.33 | 0 | 6 | 11 | ** | 0.004 |
|  |  |  | LID_CORR | LID | 1.36 | 4.62 | 14 | 8 | ## | 0.002 |
|  |  |  | LID_PREV | LID | 1.33 | 4.62 | 6 | 8 | # | 0.030 |
|  |  |  | LID_PREV | LID_CORR | 1.33 | 1.36 | 6 | 14 | ns | 0.586 |
| Fig. 1f | Axial | 8 | LID | SHAM | 4.37 | 0 | 8 | 11 | *** | <0.001 |
|  |  |  | LID_CORR | SHAM | 1.36 | 0 | 14 | 11 | * | 0.011 |
|  |  |  | LID_PREV | SHAM | 2.67 | 0 | 6 | 11 | *** | <0.001 |
|  |  |  | LID_CORR | LID | 1.36 | 4.37 | 14 | 8 | ## | 0.001 |
|  |  |  | LID_PREV | LID | 2.67 | 4.37 | 6 | 8 | ns | 0.628 |
|  |  |  | LID_PREV | LID_CORR | 2.67 | 1.36 | 6 | 14 | && | 0.001 |
| Fig. 1f | Axial | 9 | LID | SHAM | 5.85 | 0 | 8 | 11 | *** | <0.001 |
|  |  |  | LID_CORR | SHAM | 1.27 | 0 | 11 | 11 | ** | 0.005 |
|  |  |  | LID_PREV | SHAM | 2.83 | 0 | 6 | 11 | *** | <0.001 |
|  |  |  | LID_CORR | LID | 1.27 | 5.85 | 11 | 8 | ## | 0.001 |
|  |  |  | LID_PREV | LID | 2.83 | 5.85 | 6 | 8 | ns | 0.246 |
|  |  |  | LID_PREV | LID_CORR | 2.83 | 1.27 | 6 | 11 | & | 0.021 |
| Fig. 1g | Limb | 4 | LID | SHAM | 3.89 | 0.07 | 9 | 14 | *** | <0.001 |
|  |  |  | LID_CORR | SHAM | 1.20 | 0.07 | 15 | 14 | *** | <0.001 |
|  |  |  | LID_PREV | SHAM | 3.12 | 0.07 | 8 | 14 | *** | <0.001 |
|  |  |  | LID_CORR | LID | 1.20 | 3.89 | 15 | 9 | ## | 0.003 |
|  |  |  | LID_PREV | LID | 3.12 | 3.89 | 8 | 9 | ns | 0.981 |
|  |  |  | LID_PREV | LID_CORR | 3.12 | 1.20 | 8 | 15 | && | 0.002 |
| Fig. 1g | Limb | 5 | LID | SHAM | 3.57 | 0.09 | 7 | 11 | *** | <0.001 |
|  |  |  | LID_CORR | SHAM | 0.83 | 0.09 | 6 | 11 | * | 0.034 |
|  |  |  | LID_PREV | SHAM | 2.67 | 0.09 | 9 | 11 | *** | <0.001 |
|  |  |  | LID_CORR | LID | 0.83 | 3.57 | 6 | 7 | ## | 0.004 |
|  |  |  | LID_PREV | LID | 2.67 | 3.57 | 9 | 7 | ns | 0.375 |
|  |  |  | LID_PREV | LID_CORR | 2.67 | 0.83 | 9 | 6 | & | 0.015 |
| Fig. 1g | Limb | 6 | LID | SHAM | 4.22 | 0 | 9 | 13 | *** | <0.001 |
|  |  |  | LID_CORR | SHAM | 1.53 | 0 | 15 | 13 | *** | <0.001 |
|  |  |  | LID_PREV | SHAM | 2.89 | 0 | 9 | 13 | *** | <0.001 |
|  |  |  | LID_CORR | LID | 1.53 | 4.22 | 15 | 9 | ### | <0.001 |
|  |  |  | LID_PREV | LID | 2.89 | 4.22 | 9 | 9 | ns | 0.476 |
|  |  |  | LID_PREV | LID_CORR | 2.89 | 1.53 | 9 | 15 | && | 0.003 |
| Fig. 1g | Limb | 7 | LID | SHAM | 3.44 | 0.08 | 9 | 12 | *** | <0.001 |
|  |  |  | LID_CORR | SHAM | 0.86 | 0.08 | 14 | 12 | ** | 0.004 |
|  |  |  | LID_PREV | SHAM | 1.22 | 0.08 | 9 | 12 | *** | <0.001 |
|  |  |  | LID_CORR | LID | 0.86 | 3.44 | 14 | 9 | ### | <0.001 |
|  |  |  | LID_PREV | LID | 1.22 | 3.44 | 9 | 9 | # | 0.018 |
|  |  |  | LID_PREV | LID_CORR | 1.22 | 0.86 | 9 | 14 | ns | 0.302 |
| Fig. 1g | Limb | 8 | LID | SHAM | 2.22 | 0.08 | 9 | 12 | *** | <0.001 |
|  |  |  | LID_CORR | SHAM | 0.28 | 0.08 | 14 | 12 | ns | 0.264 |
|  |  |  | LID_PREV | SHAM | 2.00 | 0.08 | 9 | 12 | *** | <0.001 |
|  |  |  | LID_CORR | LID | 0.28 | 2.22 | 14 | 9 | ### | <0.001 |
|  |  |  | LID_PREV | LID | 2.00 | 2.22 | 9 | 9 | ns | 0.852 |
|  |  |  | LID_PREV | LID_CORR | 2.00 | 0.28 | 9 | 14 | &&& | <0.001 |
| Fig. 1g | Limb | 9 | LID | SHAM | 1.89 | 0.08 | 9 | 12 | *** | <0.001 |
|  |  |  | LID_CORR | SHAM | 0.09 | 0.08 | 11 | 12 | ns | 0.936 |
|  |  |  | LID_PREV | SHAM | 0.87 | 0.08 | 8 | 12 | * | 0.014 |
|  |  |  | LID_CORR | LID | 0.09 | 1.89 | 11 | 9 | ### | <0.001 |
|  |  |  | LID_PREV | LID | 0.87 | 1.89 | 8 | 9 | ns | 0.291 |
|  |  |  | LID_PREV | LID_CORR | 0.87 | 0.09 | 8 | 11 | & | 0.018 |
| Supp.<br>Fig1c | Oral | 4 | LID | SHAM | 0.62 | 0 | 8 | 7 | ns | 0.160 |
|  |  |  | LID_CORR | SHAM | 1.20 | 0 | 5 | 7 | ns | 0.150 |
|  |  |  | LID_PREV | SHAM | 0 | 0 | 5 | 7 | - | - |
|  |  |  | LID_CORR | LID | 1.20 | 0.62 | 5 | 8 | ns | 0.370 |
|  |  |  | LID_PREV | LID | 0 | 0.62 | 5 | 8 | ns | 0.210 |
|  |  |  | LID_PREV | LID_CORR | 0 | 1.20 | 5 | 5 | ns | 0.160 |
| Supp.<br>Fig1c | Oral | 5 | LID | SHAM | 0.62 | 0 | 8 | 6 | ns | 0.157 |
|  |  |  | LID_CORR | SHAM | 0.83 | 0 | 6 | 6 | ns | 0.067 |
|  |  |  | LID_PREV | SHAM | 0 | 0 | 6 | 6 | - | - |
|  |  |  | LID_CORR | LID | 0.83 | 0.62 | 6 | 8 | ns | 0.575 |
|  |  |  | LID_PREV | LID | 0 | 0.62 | 6 | 8 | ns | 0.157 |
|  |  |  | LID_PREV | LID_CORR | 0 | 0.83 | 6 | 6 | ns | 0.067 |
| Supp.<br>Fig1c | Oral | 6 | LID | SHAM | 0.75 | 0.17 | 8 | 6 | ns | 0.330 |
|  |  |  | LID_CORR | SHAM | 0 | 0.17 | 6 | 6 | ns | 0.400 |
|  |  |  | LID_PREV | SHAM | 0 | 0.17 | 6 | 6 | ns | 0.400 |
|  |  |  | LID_CORR | LID | 0 | 0.75 | 6 | 8 | ns | 0.160 |
|  |  |  | LID_PREV | LID | 0 | 0.75 | 6 | 8 | ns | 0.160 |
|  |  |  | LID_PREV | LID_CORR | 0 | 0 | 6 | 6 | - | - |
| Supp.<br>Fig1c | Oral | 7 | LID | SHAM | 1.12 | 0.17 | 8 | 6 | * | 0.028 |
|  |  |  | LID_CORR | SHAM | 0.17 | 0.17 | 6 | 6 | ns | 1 |
|  |  |  | LID_PREV | SHAM | 0.17 | 0.17 | 6 | 6 | ns | 1 |
|  |  |  | LID_CORR | LID | 0.17 | 1.12 | 6 | 8 | # | 0.028 |
|  |  |  | LID_PREV | LID | 0.17 | 1.12 | 6 | 8 | # | 0.028 |
|  |  |  | LID_PREV | LID_CORR | 0.17 | 0.17 | 6 | 6 | ns | 1 |
| Supp.<br>Fig1c | Oral | 8 | LID | SHAM | 1.12 | 0.33 | 8 | 6 | ns | 0.490 |
|  |  |  | LID_CORR | SHAM | 0.17 | 0.33 | 6 | 6 | ns | 0.590 |
|  |  |  | LID_PREV | SHAM | 0 | 0.33 | 6 | 6 | ns | 0.350 |
|  |  |  | LID_CORR | LID | 0.17 | 1.12 | 6 | 8 | ns | 0.350 |
|  |  |  | LID_PREV | LID | 0 | 1.12 | 6 | 8 | ns | 0.350 |
|  |  |  | LID_PREV | LID_CORR | 0 | 0.17 | 6 | 6 | ns | 0.49 |
| Supp.<br>Fig1c | Oral | 9 | LID | SHAM | 1.12 | 0.33 | 8 | 6 | ns | 0.550 |
|  |  |  | LID_CORR | SHAM | 0.33 | 0.33 | 6 | 6 | ns | 0.900 |
|  |  |  | LID_PREV | SHAM | 0.50 | 0.33 | 6 | 6 | ns | 0.920 |
|  |  |  | LID_CORR | LID | 0.33 | 1.12 | 6 | 8 | ns | 0.550 |
|  |  |  | LID_PREV | LID | 0.50 | 1.12 | 6 | 8 | ns | 0.590 |
|  |  |  | LID_PREV | LID_CORR | 0.50 | 0.33 | 6 | 6 | ns | 0.900 |

**Table S2:** Number of cells and mice in each condition per week for the three deep cerebellar nuclei. Related to Figure 2 and Supplementary Figure 5.

| All cell (DCN) | Region | Interposed |  | Dentate |  | Fastigial |  |
| --- | --- | --- | --- | --- | --- | --- | --- |
| Groupe | Weeks | Cells | Mice | Cells | Mice | Cells | Mice |
| SHAM | W2-W3 | n=10 | N=1 | n=28 | N=2 | n=7 | N=1 |
|  | W4-W5 | n=11 | N=1 | n=30 | N=2 | n=8 | N=1 |
|  | W6-W7 | n=10 | N=1 | n=24 | N=2 | n=3 | N=1 |
|  | W8-W9 | n=9 | N=1 | n=28 | N=2 | n=3 | N=1 |
| LID | W2-W3 | n=25 | N=3 | n=41 | N=4 | n=41 | N=5 |
|  | W4-W5 | n=8 | N=2 | n=33 | N=4 | n=42 | N=5 |
|  | W6-W7 | n=10 | N=3 | n=24 | N=3 | n=49 | N=6 |
|  | W8-W9 | n=13 | N=3 | n=30 | N=4 | n=27 | N=4 |
| LID_CORR | W2-W3 | n=19 | N=2 | n=12 | N=1 | n=15 | N=3 |
|  | W4-W5 | n=24 | N=3 | n=17 | N=2 | n=22 | N=3 |
|  | W6-W7 | n=12 | N=2 | n=10 | N=1 | n=20 | N=3 |
|  | W8-W9 | n=11 | N=2 | n=12 | N=1 | n=17 | N=3 |
| LID_PREV | W2-W3 | n=31 | N=3 | n=34 | N=3 | n=31 | N=3 |
|  | W4-W5 | n=32 | N=3 | n=26 | N=3 | n=32 | N=3 |
|  | W6-W7 | n=26 | N=3 | n=25 | N=3 | n=32 | N=3 |
|  | W8-W9 | n=27 | N=3 | n=16 | N=3 | n=29 | N=3 |

**Table S3:**
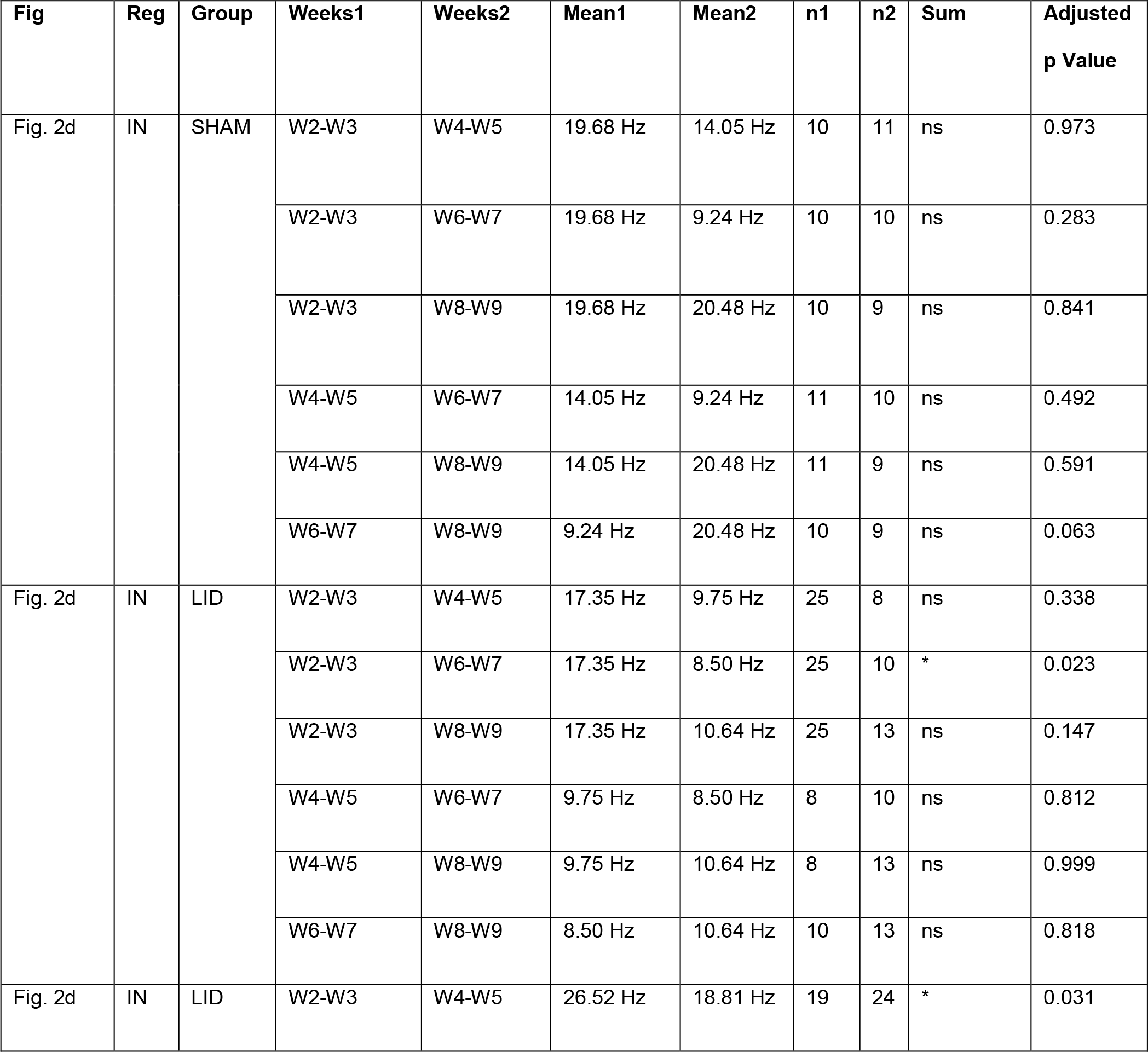
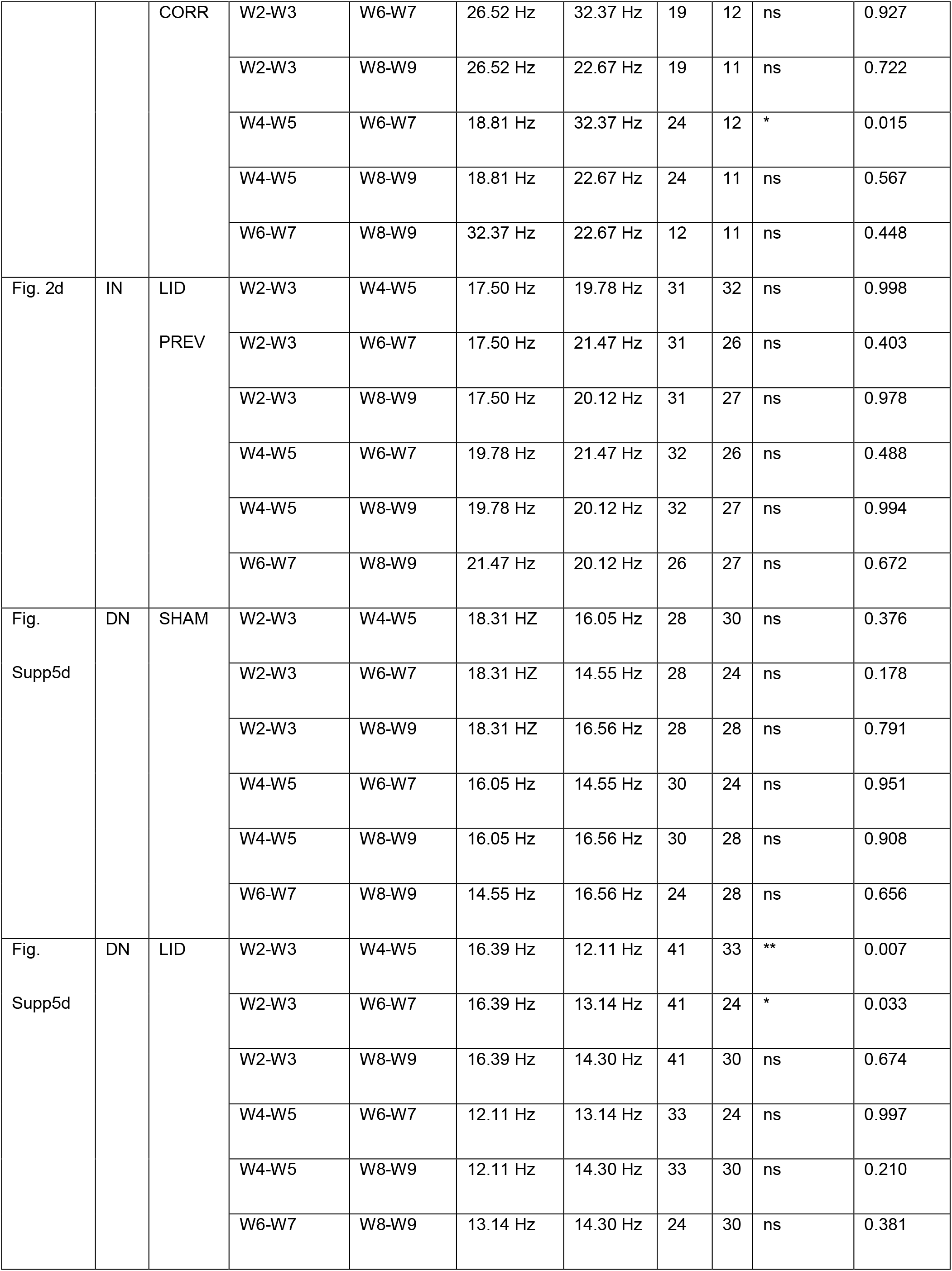
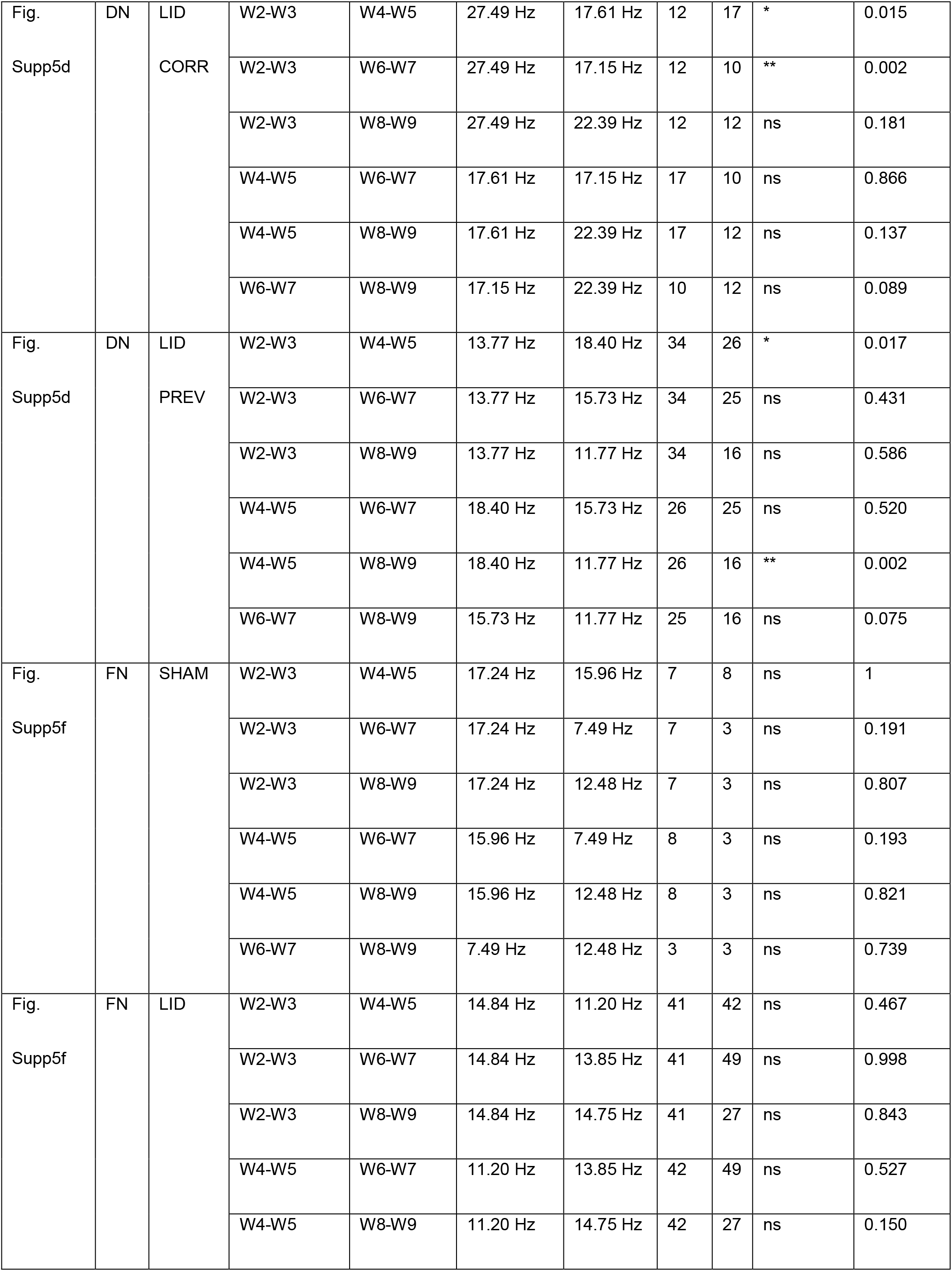
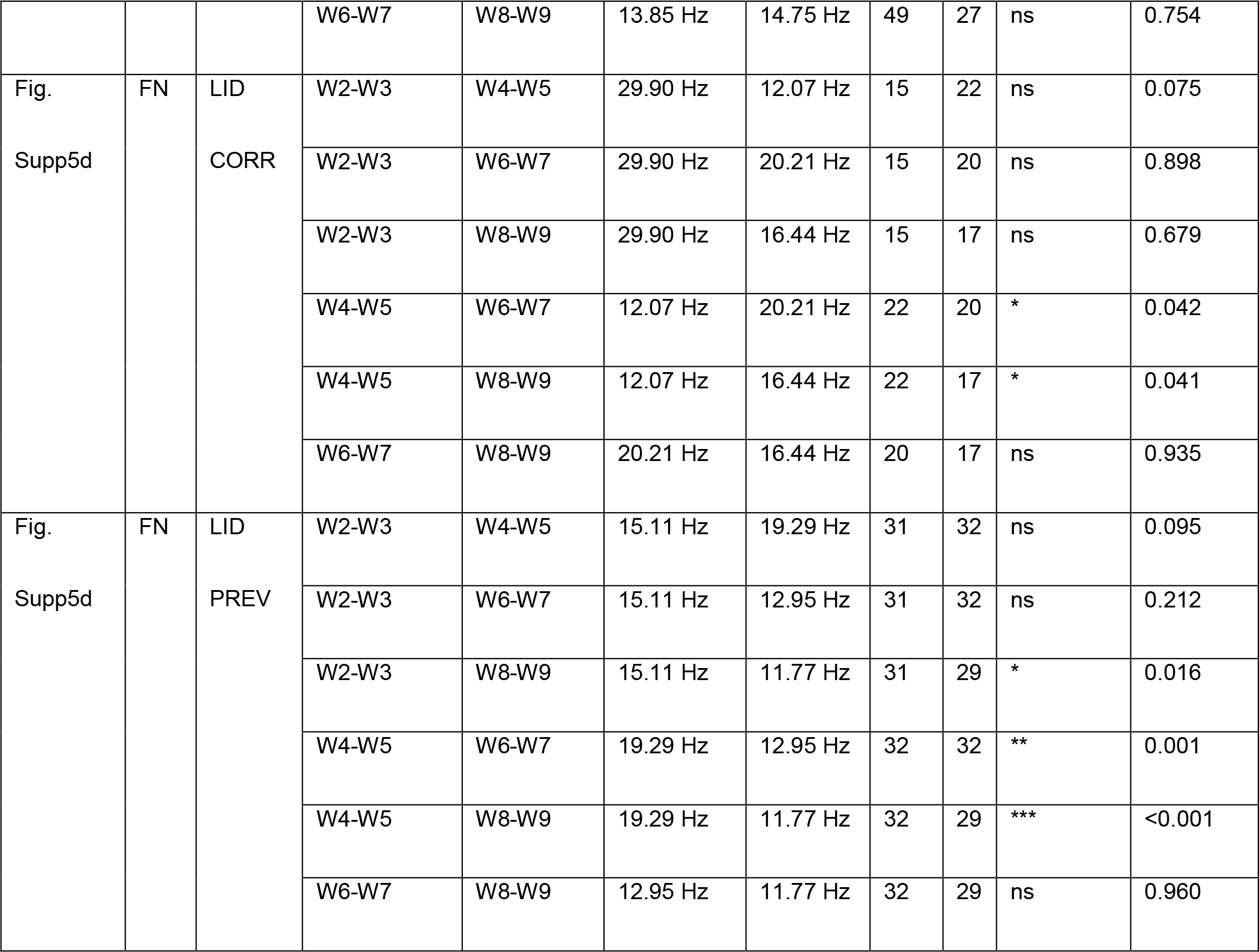
Precision measures, exact p-values, and replicate data relevant to Figure 2d and Supplementary Figures 5d and 5f.

**Table S4:**
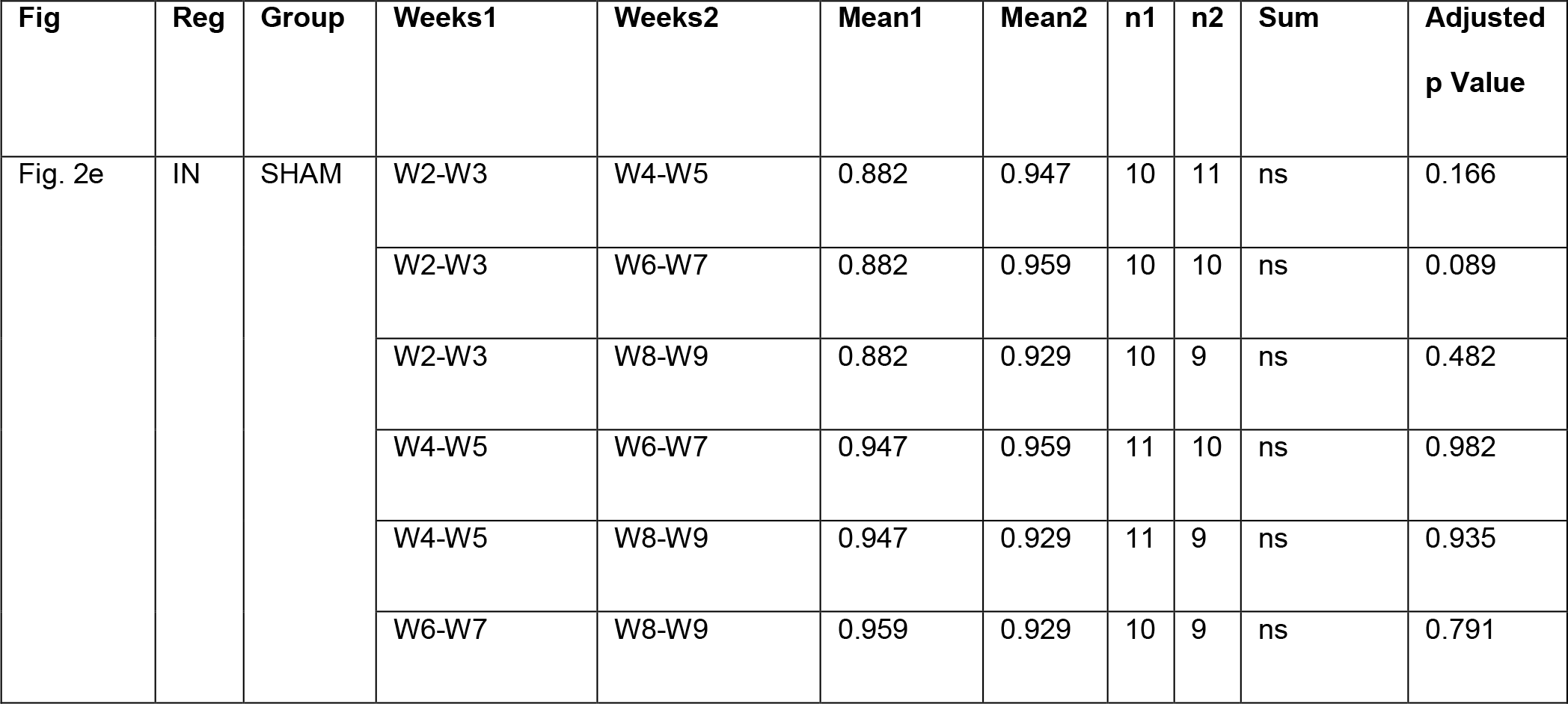
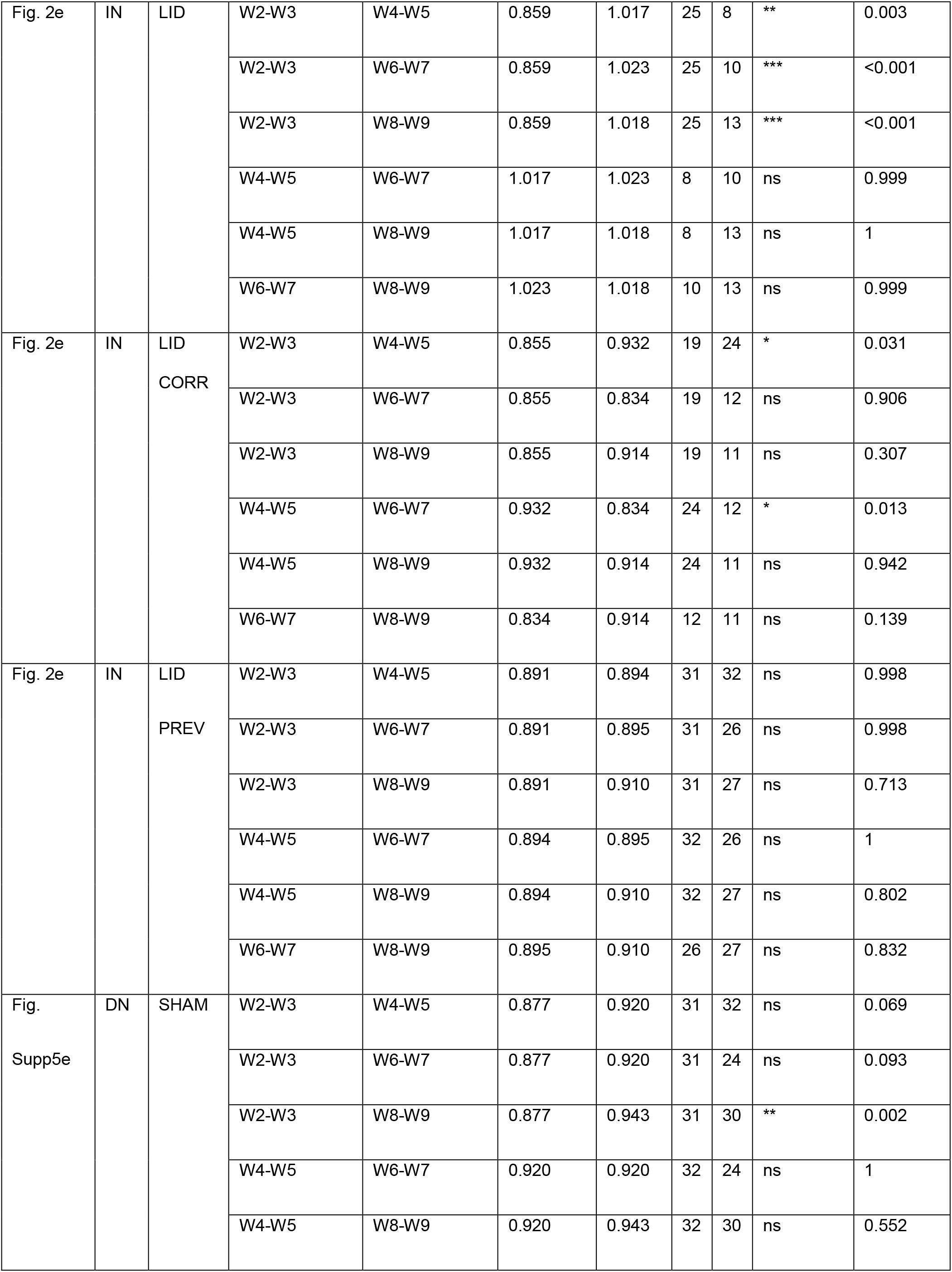
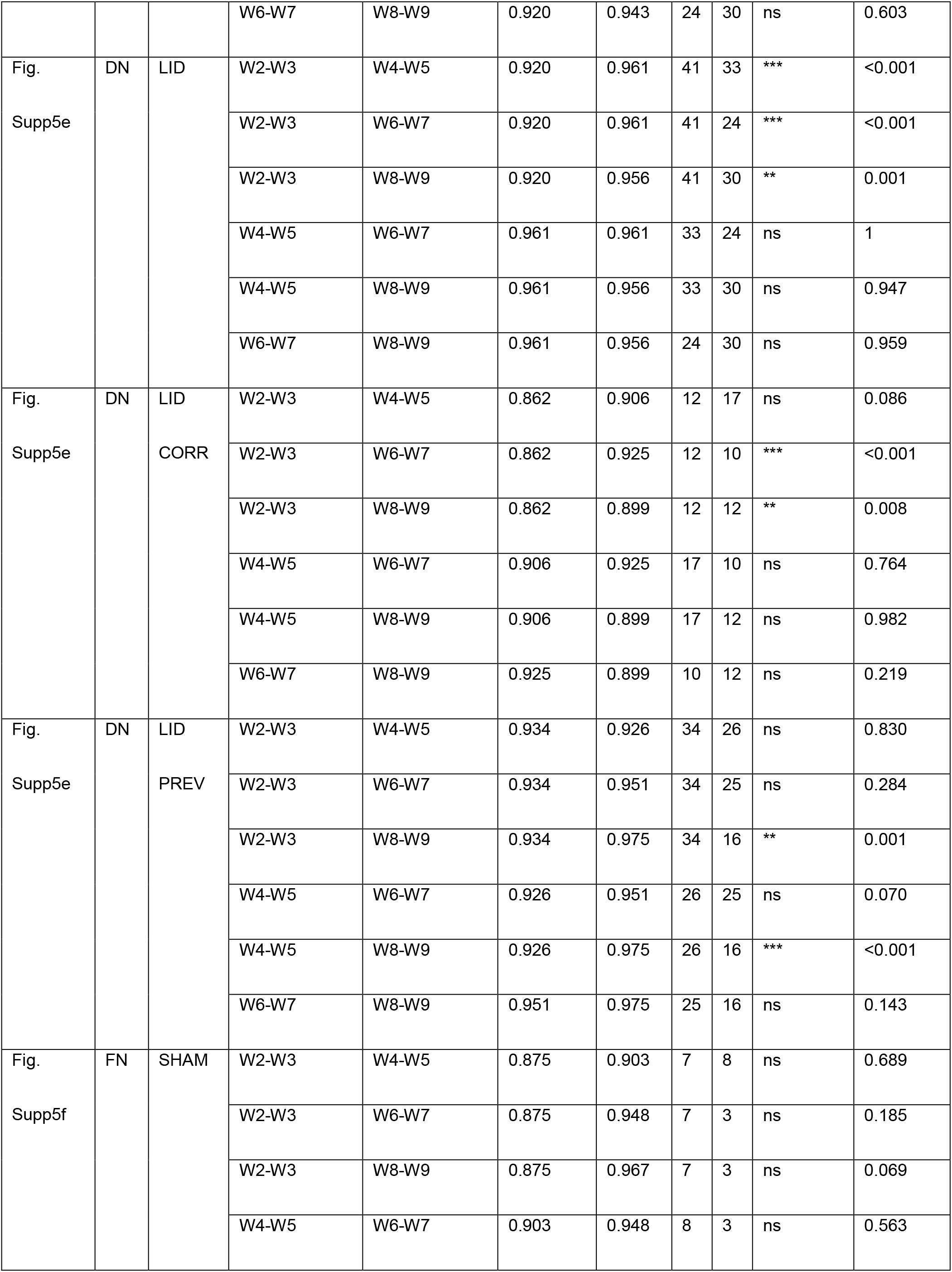
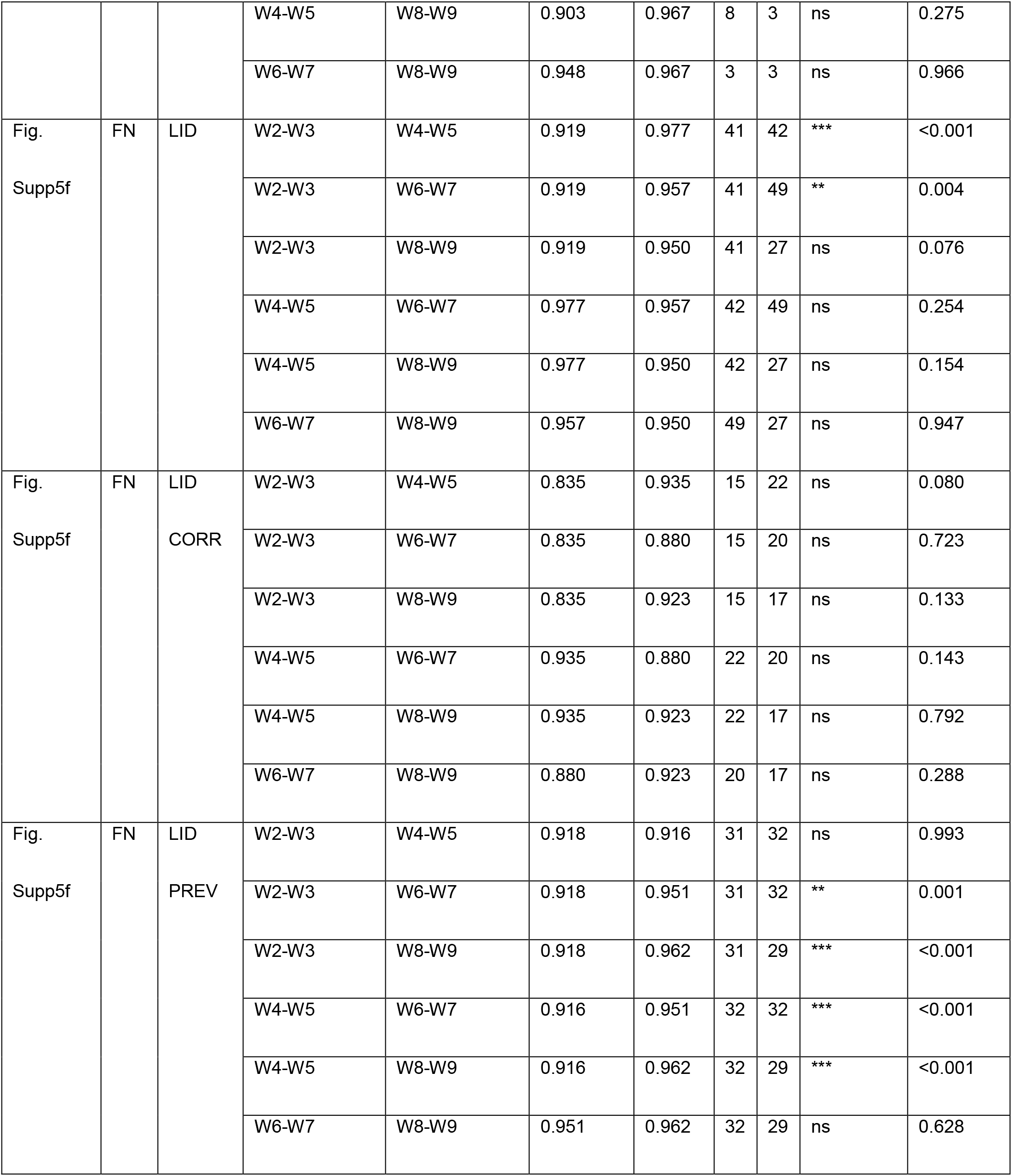
Precision measures, exact p-values, and replicate date relevant to Figure 2e and Supplementary Figures 5e and 5g.

**Table S5:**
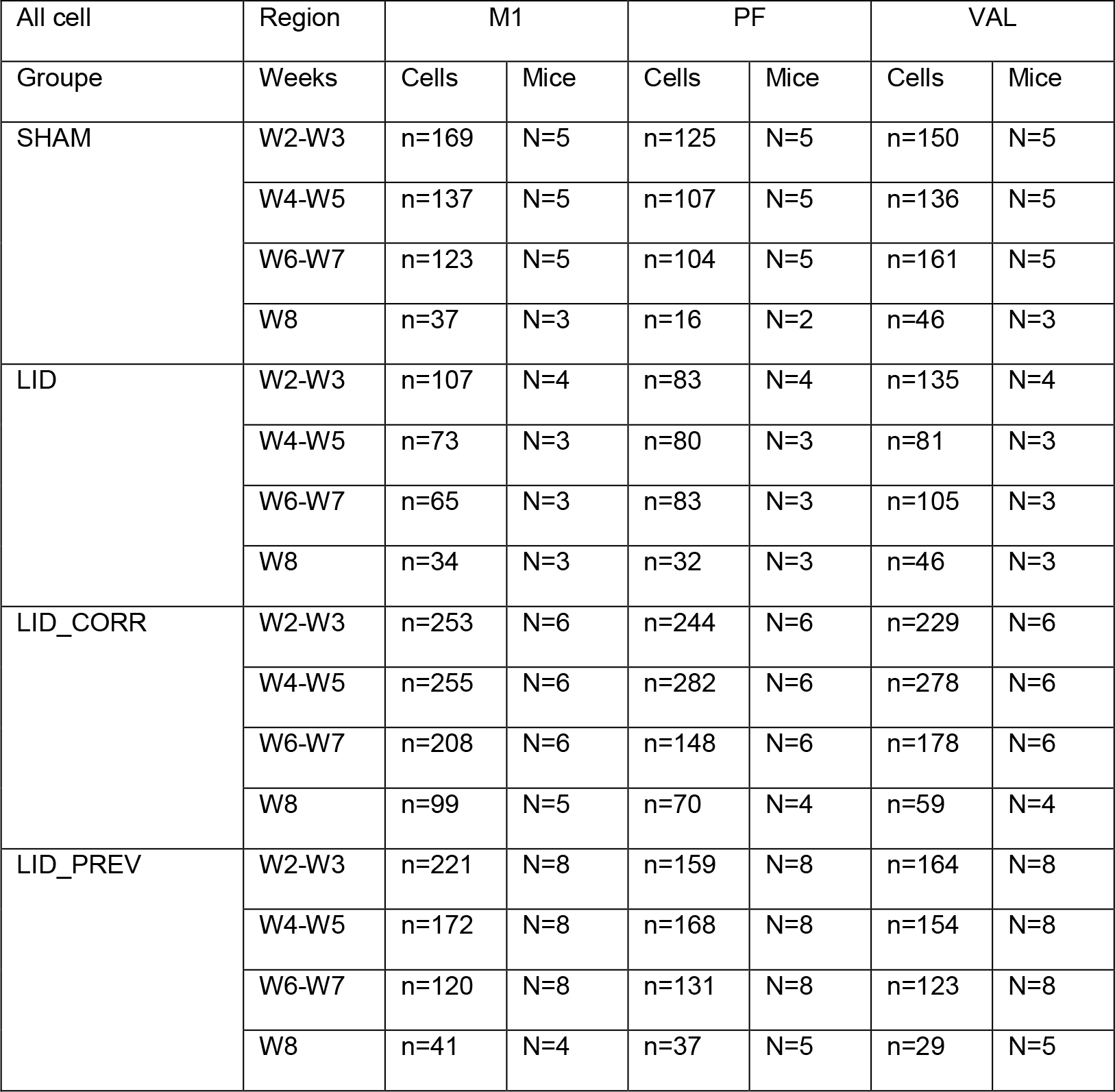
Number of cells and mice in each condition per week in the motor cortex (M1), the parafascicular nucleus (PF) of the thalamus and the ventroanterior-ventrolateral complex (VAL) of the thalamus. Related to Figures 3c, 3d and Supplementary Figure 7d.

| All cell | Region | M1 |  | PF |  | VAL |  |
| --- | --- | --- | --- | --- | --- | --- | --- |
| Groupe | Weeks | Cells | Mice | Cells | Mice | Cells | Mice |
| SHAM | W2-W3 | n=169 | N=5 | n=125 | N=5 | n=150 | N=5 |
|  | W4-W5 | n=137 | N=5 | n=107 | N=5 | n=136 | N=5 |
|  | W6-W7 | n=123 | N=5 | n=104 | N=5 | n=161 | N=5 |
|  | W8 | n=37 | N=3 | n=16 | N=2 | n=46 | N=3 |
| LID | W2-W3 | n=107 | N=4 | n=83 | N=4 | n=135 | N=4 |
|  | W4-W5 | n=73 | N=3 | n=80 | N=3 | n=81 | N=3 |
|  | W6-W7 | n=65 | N=3 | n=83 | N=3 | n=105 | N=3 |
|  | W8 | n=34 | N=3 | n=32 | N=3 | n=46 | N=3 |
| LID_CORR | W2-W3 | n=253 | N=6 | n=244 | N=6 | n=229 | N=6 |
|  | W4-W5 | n=255 | N=6 | n=282 | N=6 | n=278 | N=6 |
|  | W6-W7 | n=208 | N=6 | n=148 | N=6 | n=178 | N=6 |
|  | W8 | n=99 | N=5 | n=70 | N=4 | n=59 | N=4 |
| LID_PREV | W2-W3 | n=221 | N=8 | n=159 | N=8 | n=164 | N=8 |
|  | W4-W5 | n=172 | N=8 | n=168 | N=8 | n=154 | N=8 |
|  | W6-W7 | n=120 | N=8 | n=131 | N=8 | n=123 | N=8 |
|  | W8 | n=41 | N=4 | n=37 | N=5 | n=29 | N=5 |

**Table S6:**
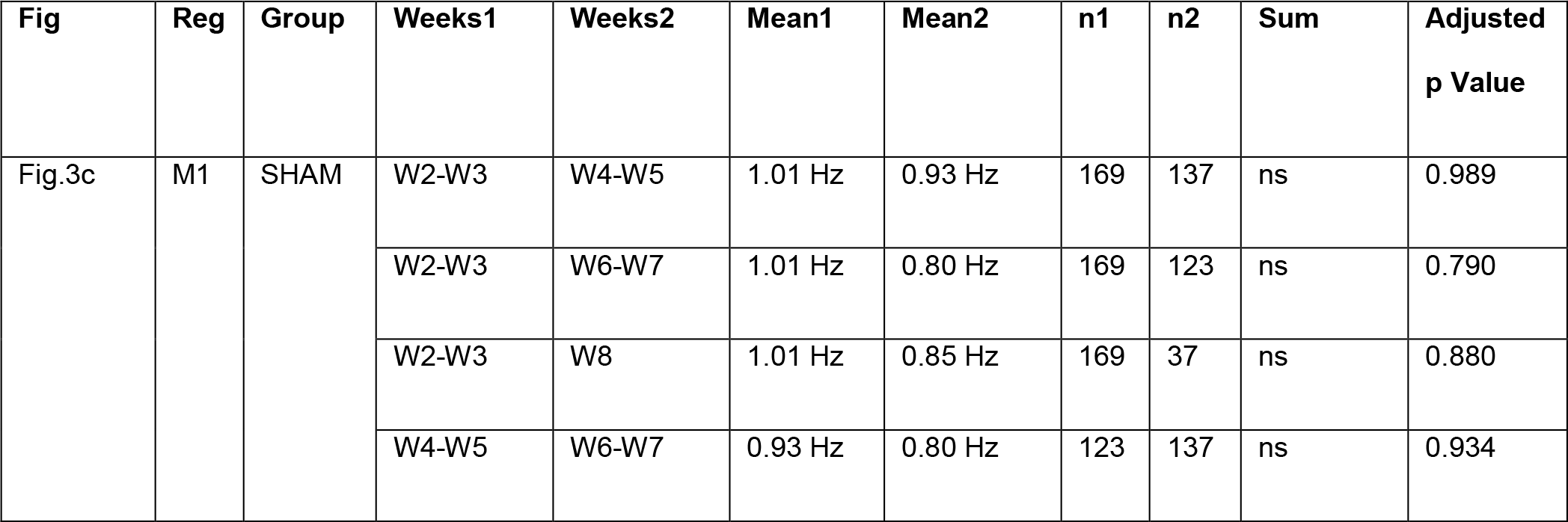
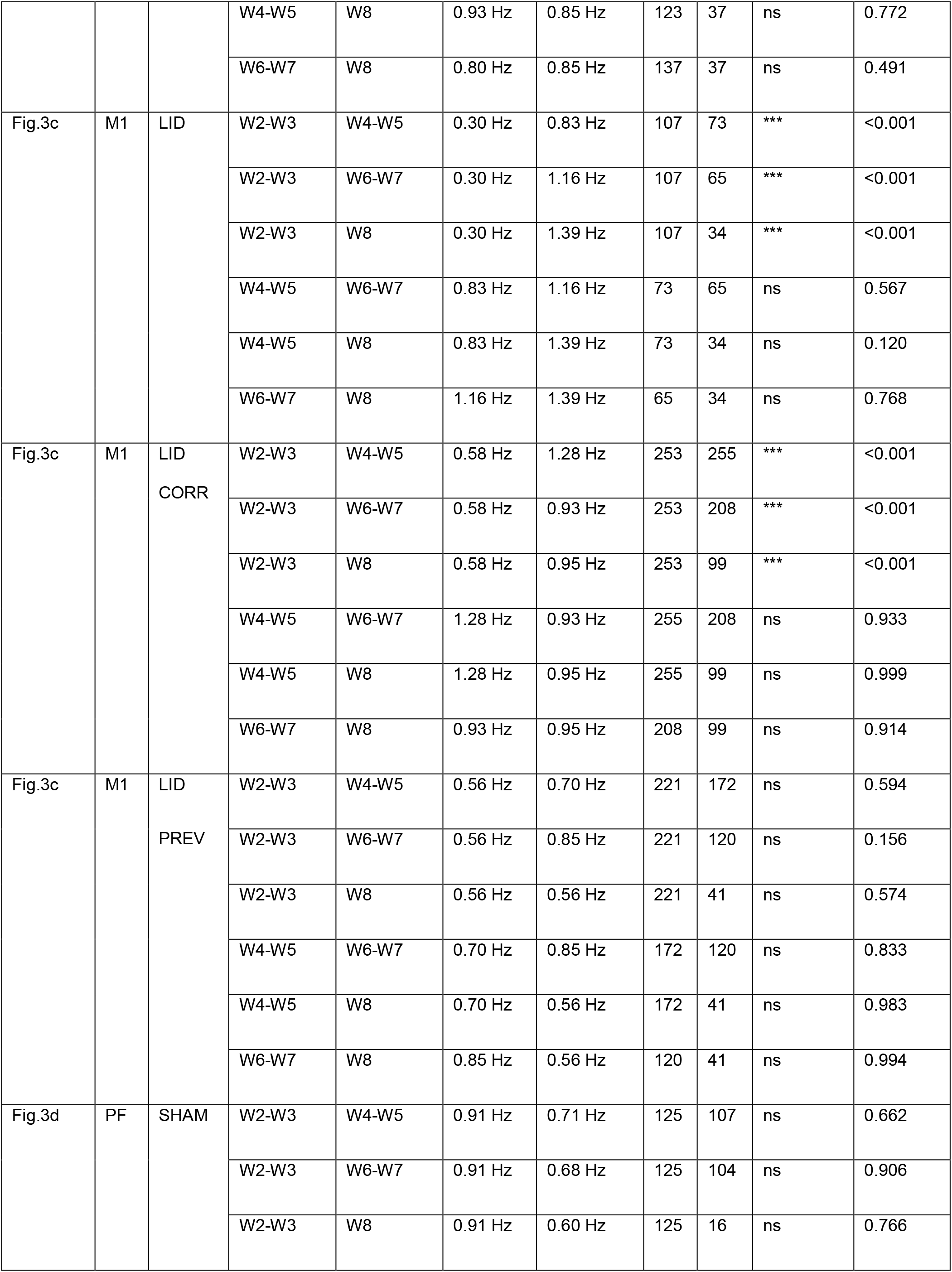
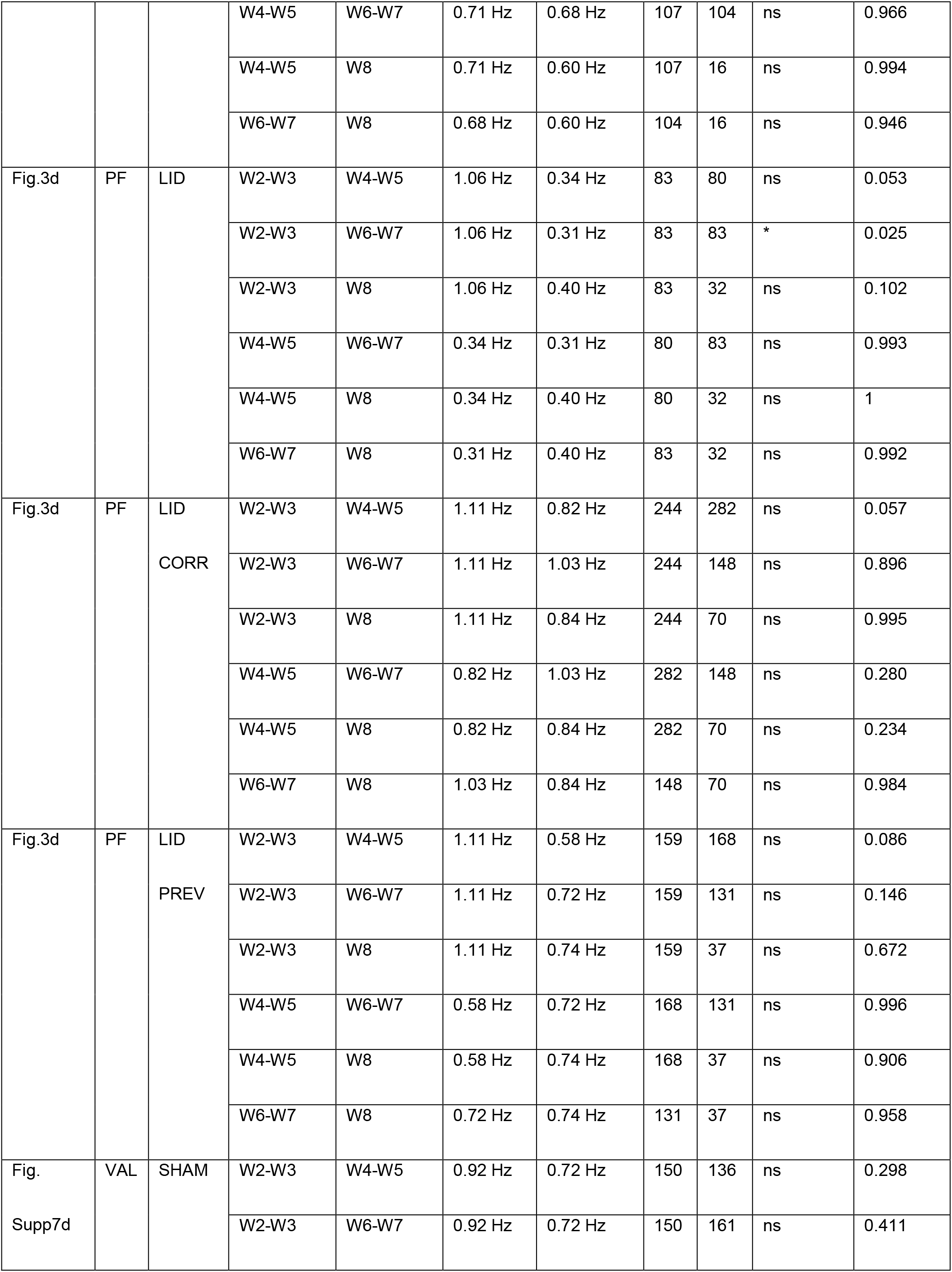
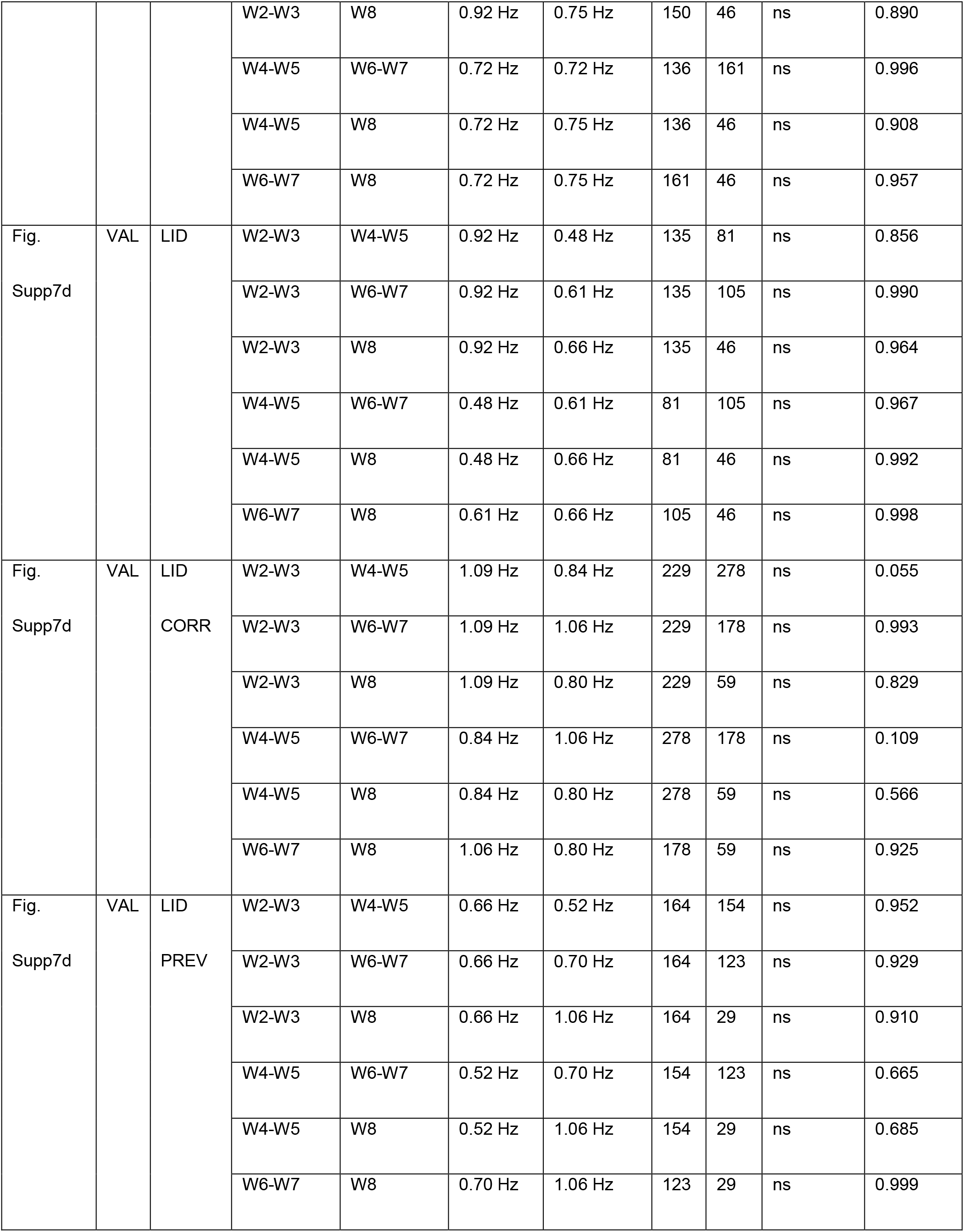
Precision measures, exact p-values, and replicate date relevant to Figures 3c, 3d and Supplementary Figure 7d.

**Table S7:**
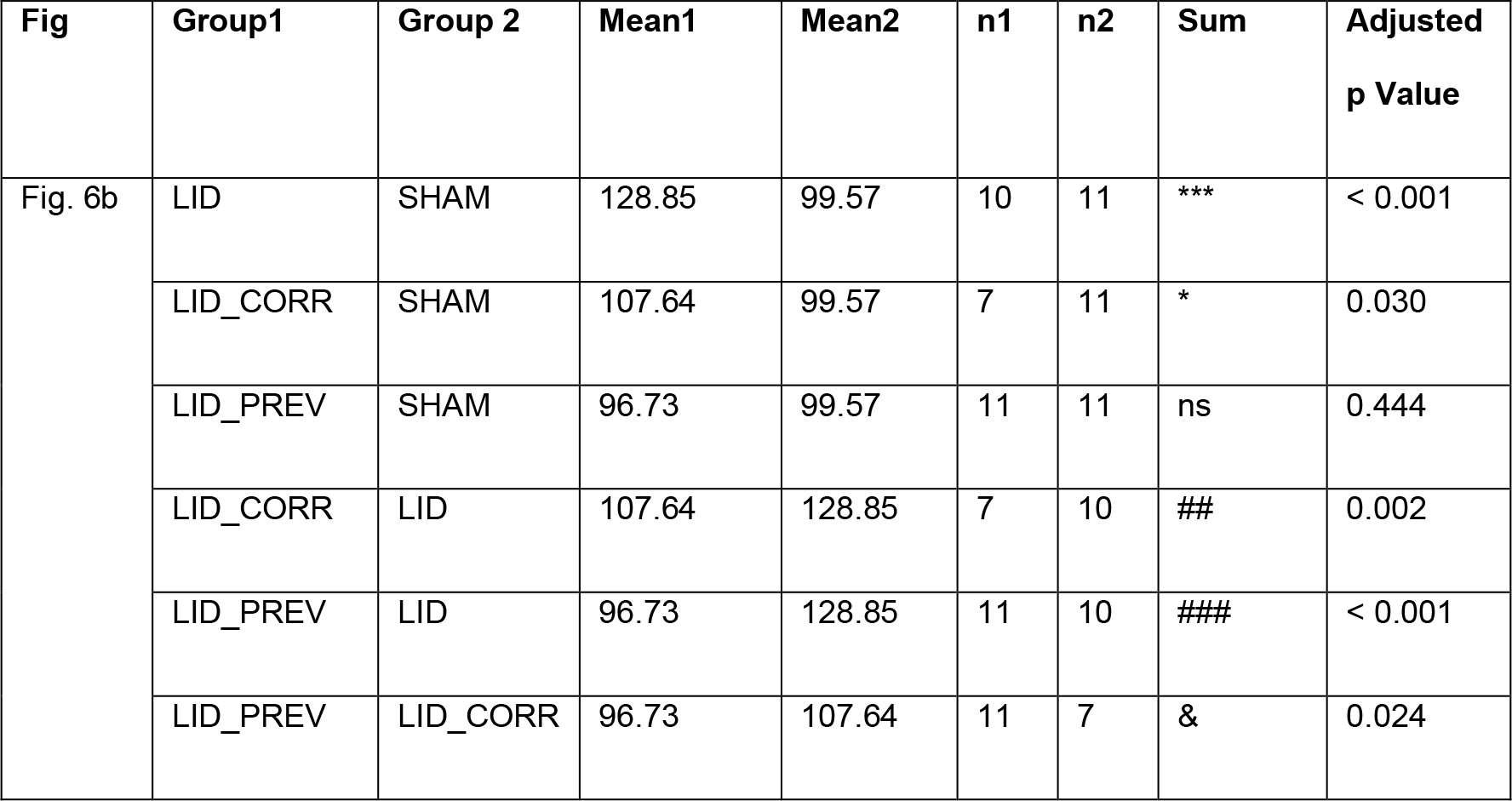
Precision measures, exact p-values, and replicate date relevant to Figure 6b.

**Table S8:** Precision measures, exact p-values, and replicate date relevant to Supplementary Figure 4.

| Fig | Param | Week | Group1 | Group2 | Mean1 | Mean2 | n1 | n2 | Sum | Adjusted p Value |
| --- | --- | --- | --- | --- | --- | --- | --- | --- | --- | --- |
| Fig. Supp4 | Oral | 4 | SHAM | SHAM_CORR | 0.50 | 0.36 | 4 | 11 | ns | 1 |
|  |  |  | SHAM | SHAM_PREV | 0.50 | 0.50 | 4 | 4 | ns | 1 |
|  |  |  | SHAM_CORR | SHAM_PREV | 0.36 | 0.50 | 11 | 4 | ns | 1 |
| Fig. Supp4 | Oral | 5 | SHAM | SHAM_CORR | 0.50 | 0.1 | 4 | 10 | ns | 0.51 |
|  |  |  | SHAM | SHAM_PREV | 0.50 | 0.33 | 4 | 3 | ns | 0.77 |
|  |  |  | SHAM_CORR | SHAM_PREV | 0.1 | 0.33 | 10 | 3 | ns | 0.60 |
| Fig. Supp4 | Oral | 6 | SHAM | SHAM_CORR | 0.50 | 0.25 | 4 | 12 | ns | 0.49 |
|  |  |  | SHAM | SHAM_PREV | 0.50 | 0 | 4 | 3 | ns | 0.49 |
|  |  |  | SHAM_CORR | SHAM_PREV | 0.25 | 0 | 12 | 3 | ns | 0.49 |
| Fig.<br>Supp4 | Oral | 7 | SHAM | SHAM_CORR | 0.50 | 0 | 4 | 8 | ns | 0.17 |
|  |  |  | SHAM | SHAM_PREV | 0.50 | 0.33 | 4 | 3 | ns | 0.77 |
|  |  |  | SHAM_CORR | SHAM_PREV | 0 | 0.33 | 8 | 3 | ns | 0.18 |
| Fig.<br>Supp4 | Oral | 8 | SHAM | SHAM_CORR | 0.75 | 0.30 | 4 | 10 | ns | 0.33 |
|  |  |  | SHAM | SHAM_PREV | 0.75 | 0 | 4 | 3 | ns | 0.33 |
|  |  |  | SHAM_CORR | SHAM_PREV | 0.30 | 0 | 10 | 3 | ns | 0.33 |
| Fig.<br>Supp4 | Oral | 9 | SHAM | SHAM_CORR | 0.25 | 0.20 | 4 | 5 | ns | 0.94 |
|  |  |  | SHAM | SHAM_PREV | 0.25 | 1 | 4 | 2 | ns | 0.33 |
|  |  |  | SHAM_CORR | SHAM_PREV | 0.20 | 1 | 5 | 2 | ns | 0.33 |
| Fig.<br>Supp4 | Axial | 4 | SHAM | SHAM_CORR | 0 | 0 | 4 | 5 | ns | - |
|  |  |  | SHAM | SHAM_PREV | 0 | 0 | 4 | 3 | ns | - |
|  |  |  | SHAM_CORR | SHAM_PREV | 0 | 0 | 5 | 3 | ns | - |
| Fig.<br>Supp4 | Axial | 5 | SHAM | SHAM_CORR | 0 | 0 | 4 | 3 | ns | - |
|  |  |  | SHAM | SHAM_PREV | 0 | 0 | 4 | 2 | ns | - |
|  |  |  | SHAM_CORR | SHAM_PREV | 0 | 0 | 3 | 2 | ns | - |
| Fig.<br>Supp4 | Axial | 6 | SHAM | SHAM_CORR | 0 | 0 | 4 | 5 | ns | - |
|  |  |  | SHAM | SHAM_PREV | 0 | 0 | 4 | 2 | ns | - |
|  |  |  | SHAM_CORR | SHAM_PREV | 0 | 0 | 5 | 2 | ns | - |
| Fig.<br>Supp4 | Axial | 7 | SHAM | SHAM_CORR | 0 | 0 | 4 | 5 | ns | - |
|  |  |  | SHAM | SHAM_PREV | 0 | 0 | 4 | 2 | ns | - |
|  |  |  | SHAM_CORR | SHAM_PREV | 0 | 0 | 5 | 2 | ns | - |
| Fig. | Axial | 8 | SHAM | SHAM_CORR | 0 | 0 | 4 | 5 | ns | - |
| Supp4 |  |  | SHAM | SHAM_PREV | 0 | 0 | 4 | 2 | ns | - |
|  |  |  | SHAM_CORR | SHAM_PREV | 0 | 0 | 5 | 2 | ns | - |
| Fig.<br>Supp4 | Axial | 9 | SHAM | SHAM_CORR | 0 | 0 | 4 | 5 | ns | - |
|  |  |  | SHAM | SHAM_PREV | 0 | 0 | 4 | 2 | ns | - |
|  |  |  | SHAM_CORR | SHAM_PREV | 0 | 0 | 5 | 2 | ns | - |
| Fig.<br>Supp4 | Limb | 4 | SHAM | SHAM_CORR | 0 | 0.14 | 4 | 7 | ns | 0.51 |
|  |  |  | SHAM | SHAM_PREV | 0 | 0 | 4 | 3 | ns | - |
|  |  |  | SHAM_CORR | SHAM_PREV | 0.14 | 0 | 7 | 3 | ns | 0.51 |
| Fig.<br>Supp4 | Limb | 5 | SHAM | SHAM_CORR | 0 | 0.20 | 4 | 5 | ns | 0.54 |
|  |  |  | SHAM | SHAM_PREV | 0 | 0 | 4 | 2 | ns | - |
|  |  |  | SHAM_CORR | SHAM_PREV | 0.20 | 0 | 5 | 2 | ns | 0.54 |
| Fig.<br>Supp4 | Limb | 6 | SHAM | SHAM_CORR | 0 | 0 | 4 | 7 | ns | - |
|  |  |  | SHAM | SHAM_PREV | 0 | 0 | 4 | 2 | ns | - |
|  |  |  | SHAM_CORR | SHAM_PREV | 0 | 0 | 7 | 2 | ns | - |
| Fig.<br>Supp4 | Limb | 7 | SHAM | SHAM_CORR | 0 | 0.17 | 4 | 6 | ns | 0.58 |
|  |  |  | SHAM | SHAM_PREV | 0 | 0 | 4 | 2 | ns | - |
|  |  |  | SHAM_CORR | SHAM_PREV | 0.17 | 0 | 6 | 2 | ns | 0.58 |
| Fig.<br>Supp4 | Limb | 8 | SHAM | SHAM_CORR | 0 | 0.17 | 4 | 6 | ns | 0.58 |
|  |  |  | SHAM | SHAM_PREV | 0 | 0 | 4 | 2 | ns | - |
|  |  |  | SHAM_CORR | SHAM_PREV | 0.17 | 0 | 6 | 2 | ns | 0.58 |
| Fig.<br>Supp4 | Limb | 9 | SHAM | SHAM_CORR | 0 | 0.17 | 4 | 6 | ns | 0.58 |
|  |  |  | SHAM | SHAM_PREV | 0 | 0 | 4 | 2 | ns | - |
|  |  |  | SHAM_CORR | SHAM_PREV | 0.17 | 0 | 6 | 2 | ns | 0.58 |

**Table S9:** Precision measures, exact p-values, and replicate date relevant to Supplementary Figure 7.

| Fig | Reg | Group | Weeks1 | Weeks2 | Mean1 | Mean2 | n1 | n2 | Sum | Adjusted<br>p Value |
| --- | --- | --- | --- | --- | --- | --- | --- | --- | --- | --- |
| Fig.<br>Supp7<br>(upper) | IN | SHAM | W2-W3 | W4-W5 | 2.64 Hz | 2.52 Hz | 10 | 11 | ns | 0.975 |
|  |  |  | W2-W3 | W6-W7 | 2.64 Hz | 2.12 Hz | 10 | 10 | ns | 0.551 |
|  |  |  | W2-W3 | W8-W9 | 2.64 Hz | 2.88 Hz | 10 | 9 | ns | 0.854 |
|  |  |  | W4-W5 | W6-W7 | 2.52 Hz | 2.12 Hz | 11 | 10 | ns | 0.743 |
|  |  |  | W4-W5 | W8-W9 | 2.52 Hz | 2.88 Hz | 11 | 9 | ns | 0.617 |
|  |  |  | W6-W7 | W8-W9 | 2.12 Hz | 2.88 Hz | 10 | 9 | ns | 0.223 |
| Fig.<br>Supp7<br>(upper) | IN | SHAM<br>CORR | W2-W3 | W4-W5 | 2.37 Hz | 1.89 Hz | 13 | 21 | ns | 0.228 |
|  |  |  | W2-W3 | W6-W7 | 2.37 Hz | 2.27 Hz | 13 | 21 | *** | <0.001 |
|  |  |  | W2-W3 | W8-W9 | 2.37 Hz | 2.28 Hz | 13 | 24 | ns | 0.472 |
|  |  |  | W4-W5 | W6-W7 | 1.89 Hz | 2.27 Hz | 21 | 21 | * | 0.018 |
|  |  |  | W4-W5 | W8-W9 | 1.89 Hz | 2.28 Hz | 21 | 24 | ns | 0.914 |
|  |  |  | W6-W7 | W8-W9 | 2.27 Hz | 2.28 Hz | 21 | 24 | ** | 0.002 |
| Fig.<br>Supp7<br>(upper) | IN | SHAM<br>PREV | W2-W3 | W4-W5 | 2.84 Hz | 1.92 Hz | 18 | 23 | *** | < 0.001 |
|  |  |  | W2-W3 | W6-W7 | 2.84 Hz | 2.14 Hz | 18 | 19 | *** | < 0.001 |
|  |  |  | W2-W3 | W8-W9 | 2.84 Hz | 2.32 Hz | 18 | 16 | * | 0.016 |
|  |  |  | W4-W5 | W6-W7 | 1.92 Hz | 2.14 Hz | 23 | 19 | ns | 0.492 |
|  |  |  | W4-W5 | W8-W9 | 1.92 Hz | 2.32 Hz | 23 | 16 | ns | 0.076 |
|  |  |  | W6-W7 | W8-W9 | 2.14 Hz | 2.32 Hz | 19 | 16 | ns | 0.716 |
| Fig.<br>Supp7<br>(upper) | DN | SHAM | W2-W3 | W4-W5 | 2.82 Hz | 2.66 Hz | 28 | 30 | ns | 0.406 |
|  |  |  | W2-W3 | W6-W7 | 2.82 Hz | 2.60 Hz | 28 | 24 | ns | 0.742 |
|  |  |  | W2-W3 | W8-W9 | 2.82 Hz | 2.73 Hz | 28 | 28 | ns | 0.806 |
|  |  |  | W4-W5 | W6-W7 | 2.66 Hz | 2.60 Hz | 30 | 24 | ns | 0.998 |
|  |  |  | W4-W5 | W8-W9 | 2.66 Hz | 2.73 Hz | 30 | 28 | ns | 0.916 |
|  |  |  | W6-W7 | W8-W9 | 2.60 Hz | 2.73 Hz | 24 | 28 | ns | 0.954 |
| Fig.<br>Supp7<br>(upper) | DN | SHAM<br>PREV | W2-W3 | W4-W5 | 3.14 Hz | 2.15 Hz | 10 | 12 | *** | < 0.001 |
|  |  |  | W2-W3 | W6-W7 | 3.14 Hz | 2.08 Hz | 10 | 11 | *** | < 0.001 |
|  |  |  | W2-W3 | W8-W9 | 3.14 Hz | 2.57 Hz | 10 | 13 | * | 0.024 |
|  |  |  | W4-W5 | W6-W7 | 2.15 Hz | 2.08 Hz | 12 | 11 | ns | 0.982 |
|  |  |  | W4-W5 | W8-W9 | 2.15 Hz | 2.57 Hz | 12 | 13 | ns | 0.126 |
|  |  |  | W6-W7 | W8-W9 | 2.08 Hz | 2.57 Hz | 11 | 13 | ns | 0.062 |
| Fig.<br>Supp7<br>(upper) | FN | SHAM | W2-W3 | W4-W5 | 2.71 Hz | 2.69 Hz | 7 | 8 | ns | 1 |
|  |  |  | W2-W3 | W6-W7 | 2.71 Hz | 1.99 Hz | 7 | 3 | ns | 0.191 |
|  |  |  | W2-W3 | W8-W9 | 2.71 Hz | 2.40 Hz | 7 | 3 | ns | 0.808 |
|  |  |  | W4-W5 | W6-W7 | 2.69 Hz | 1.99 Hz | 8 | 3 | ns | 0.193 |
|  |  |  | W4-W5 | W8-W9 | 2.69 Hz | 2.40 Hz | 8 | 3 | ns | 0.821 |
|  |  |  | W6-W7 | W8-W9 | 1.99 Hz | 2.40 Hz | 3 | 3 | ns | 0.739 |
| Fig.<br>Supp7<br>(upper) | FN | SHAM<br>PREV | W2-W3 | W4-W5 | 3.25 Hz | 2.34 Hz | 18 | 14 | *** | <0.001 |
|  |  |  | W2-W3 | W6-W7 | 3.25 Hz | 1.99 Hz | 18 | 15 | *** | <0.001 |
|  |  |  | W2-W3 | W8-W9 | 3.25 Hz | 2.28 Hz | 18 | 16 | *** | <0.001 |
|  |  |  | W4-W5 | W6-W7 | 2.34 Hz | 1.99 Hz | 14 | 15 | ns | 0.290 |
|  |  |  | W4-W5 | W8-W9 | 2.34 Hz | 2.28 Hz | 14 | 16 | ns | 0.990 |
|  |  |  | W6-W7 | W8-W9 | 1.99 Hz | 2.28 Hz | 15 | 16 | ns | 0.423 |
| Fig.<br>Supp7<br>(middle) | IN | SHAM | W2-W3 | W4-W5 | 28.58 Hz | 22.26 Hz | 10 | 11 | ns | 0.582 |
|  |  |  | W2-W3 | W6-W7 | 28.58 Hz | 12.35 Hz | 10 | 5 | * | 0.047 |
|  |  |  | W2-W3 | W8-W9 | 28.58 Hz | 27.84 Hz | 10 | 9 | ns | 0.999 |
|  |  |  | W4-W5 | W6-W7 | 22.26 Hz | 12.35 Hz | 11 | 5 | ns | 0.104 |
|  |  |  | W4-W5 | W8-W9 | 22.26 Hz | 27.84 Hz | 11 | 9 | ns | 0.725 |
|  |  |  | W6-W7 | W8-W9 | 12.35 Hz | 27.84 Hz | 5 | 9 | ns | 0.087 |
| Fig.<br>Supp7<br>(middle) | IN | SHAM<br>CORR | W2-W3 | W4-W5 | 38.92 Hz | 14.03 Hz | 7 | 9 | *** | <0.001 |
|  |  |  | W2-W3 | W6-W7 | 38.92 Hz | 24.58 Hz | 7 | 9 | ** | 0.008 |
|  |  |  | W2-W3 | W8-W9 | 38.92 Hz | 26.18 Hz | 7 | 12 | * | 0.013 |
|  |  |  | W4-W5 | W6-W7 | 14.03 Hz | 24.58 Hz | 9 | 9 | * | 0.048 |
|  |  |  | W4-W5 | W8-W9 | 14.03 Hz | 26.18 Hz | 9 | 12 | * | 0.010 |
|  |  |  | W6-W7 | W8-W9 | 24.58 Hz | 26.18 Hz | 9 | 12 | ns | 0.970 |
| Fig.<br>Supp7<br>(middle) | IN | SHAM<br>PREV | W2-W3 | W4-W5 | 28.28 Hz | 18.96 Hz | 18 | 23 | ** | 0.001 |
|  |  |  | W2-W3 | W6-W7 | 28.28 Hz | 19.39 Hz | 18 | 19 | ** | 0.004 |
|  |  |  | W2-W3 | W8-W9 | 28.28 Hz | 21.53 Hz | 18 | 16 | ns | 0.060 |
|  |  |  | W4-W5 | W6-W7 | 18.96 Hz | 19.39 Hz | 23 | 19 | ns | 0.998 |
|  |  |  | W4-W5 | W8-W9 | 18.96 Hz | 21.53 Hz | 23 | 16 | ns | 0.735 |
|  |  |  | W6-W7 | W8-W9 | 19.39 Hz | 21.53 Hz | 19 | 16 | ns | 0.846 |
| Fig.<br>Supp7<br>(middle) | DN | SHAM | W2-W3 | W4-W5 | 29.96 Hz | 22.64 Hz | 28 | 30 | ns | 0.191 |
|  |  |  | W2-W3 | W6-W7 | 29.96 Hz | 19.02 Hz | 28 | 5 | ns | 0.369 |
|  |  |  | W2-W3 | W8-W9 | 29.96 Hz | 26.47 Hz | 28 | 28 | ns | 0.782 |
|  |  |  | W4-W5 | W6-W7 | 22.64 Hz | 19.02 Hz | 30 | 5 | ns | 0.949 |
|  |  |  | W4-W5 | W8-W9 | 22.64 Hz | 26.47 Hz | 30 | 28 | ns | 0.718 |
|  |  |  | W6-W7 | W8-W9 | 19.02 Hz | 26.47 Hz | 5 | 28 | ns | 0.685 |
| Fig.<br>Supp7<br>(middle) | DN | SHAM<br>PREV | W2-W3 | W4-W5 | 29.52 Hz | 18.95 Hz | 10 | 12 | ** | 0.006 |
|  |  |  | W2-W3 | W6-W7 | 29.52 Hz | 21.48 Hz | 10 | 11 | ns | 0.060 |
|  |  |  | W2-W3 | W8-W9 | 29.52 Hz | 19.48 Hz | 10 | 13 | ** | 0.008 |
|  |  |  | W4-W5 | W6-W7 | 18.95 Hz | 21.48 Hz | 12 | 11 | ns | 0.828 |
|  |  |  | W4-W5 | W8-W9 | 18.95 Hz | 19.48 Hz | 12 | 13 | ns | 0.997 |
|  |  |  | W6-W7 | W8-W9 | 21.48 Hz | 19.48 Hz | 11 | 13 | ns | 0.902 |
| Fig.<br>Supp7<br>(middle) | FN | SHAM | W2-W3 | W4-W5 | 24.46 Hz | 25.18 Hz | 7 | 8 | ns | 0.988 |
|  |  |  | W2-W3 | W8-W9 | 24.46 Hz | 19.50 Hz | 7 | 3 | ns | 0.737 |
|  |  |  | W4-W5 | W8-W9 | 25.18 Hz | 19.50 Hz | 8 | 3 | ns | 0.662 |
| Fig.<br>Supp7<br>(middle) | FN | SHAM<br>PREV | W2-W3 | W4-W5 | 34.79 Hz | 15.57 Hz | 18 | 14 | *** | <0.001 |
|  |  |  | W2-W3 | W6-W7 | 34.79 Hz | 12.11 Hz | 18 | 15 | *** | <0.001 |
|  |  |  | W2-W3 | W8-W9 | 34.79 Hz | 18.13 Hz | 18 | 16 | *** | <0.001 |
|  |  |  | W4-W5 | W6-W7 | 15.57 Hz | 12.11 Hz | 14 | 15 | ns | 0.774 |
|  |  |  | W4-W5 | W8-W9 | 15.57 Hz | 18.13 Hz | 14 | 16 | ns | 0.888 |
|  |  |  | W6-W7 | W8-W9 | 12.11 Hz | 18.13 Hz | 15 | 16 | ns | 0.320 |
| Fig.<br>Supp7<br>(lower) | IN | SHAM | W2-W3 | W4-W5 | 18.12 Hz | 13.31 Hz | 10 | 11 | ns | 0.749 |
|  |  |  | W2-W3 | W6-W7 | 18.12 Hz | 7.41 Hz | 10 | 5 | ns | 0.303 |
|  |  |  | W2-W3 | W8-W9 | 18.12 Hz | 18.49 Hz | 10 | 9 | ns | 1 |
|  |  |  | W4-W5 | W6-W7 | 13.31 Hz | 7.41 Hz | 11 | 5 | ns | 0.754 |
|  |  |  | W4-W5 | W8-W9 | 13.31 Hz | 18.49 Hz | 11 | 9 | ns | 0.723 |
|  |  |  | W6-W7 | W8-W9 | 7.41 Hz | 18.49 Hz | 5 | 9 | ns | 0.290 |
| Fig.<br>Supp7<br>(lower) | IN | SHAM<br>CORR | W2-W3 | W4-W5 | 11.99 Hz | 10.13 Hz | 12 | 9 | ns | 0.561 |
|  |  |  | W2-W3 | W6-W7 | 11.99 Hz | 6.01 Hz | 12 | 9 | *** | < 0.001 |
|  |  |  | W2-W3 | W8-W9 | 11.99 Hz | 10.57 Hz | 12 | 12 | ns | 0.703 |
|  |  |  | W4-W5 | W6-W7 | 10.13 Hz | 6.01 Hz | 9 | 9 | * | 0.047 |
|  |  |  | W4-W5 | W8-W9 | 10.13 Hz | 10.57 Hz | 9 | 12 | ns | 0.989 |
|  |  |  | W6-W7 | W8-W9 | 6.01 Hz | 10.57 Hz | 9 | 12 | * | 0.013 |
| Fig.<br>Supp7<br>(lower) | IN | SHAM<br>PREV | W2-W3 | W4-W5 | 15.70 Hz | 6.49 Hz | 18 | 23 | *** | <0.001 |
|  |  |  | W2-W3 | W6-W7 | 15.70 Hz | 8.38 Hz | 18 | 19 | ** | 0.002 |
|  |  |  | W2-W3 | W8-W9 | 15.70 Hz | 10.02 Hz | 18 | 16 | * | 0.036 |
|  |  |  | W4-W5 | W6-W7 | 6.49 Hz | 8.38 Hz | 23 | 19 | ns | 0.307 |
|  |  |  | W4-W5 | W8-W9 | 6.49 Hz | 10.02 Hz | 23 | 16 | ns | 0.074 |
|  |  |  | W6-W7 | W8-W9 | 8.38 Hz | 10.02 Hz | 19 | 16 | ns | 0.703 |
| Fig.<br>Supp7<br>(lower) | DN | SHAM | W2-W3 | W4-W5 | 15.57 Hz | 15.49 Hz | 28 | 30 | ns | 1 |
|  |  |  | W2-W3 | W6-W7 | 15.57 Hz | 14.47 Hz | 28 | 19 | ns | 0.980 |
|  |  |  | W2-W3 | W8-W9 | 15.57 Hz | 15.62 Hz | 28 | 28 | ns | 1 |
|  |  |  | W4-W5 | W6-W7 | 15.49 Hz | 14.47 Hz | 30 | 19 | ns | 0.984 |
|  |  |  | W4-W5 | W8-W9 | 15.49 Hz | 15.62 Hz | 30 | 28 | ns | 1 |
|  |  |  | W6-W7 | W8-W9 | 14.47 Hz | 15.62 Hz | 19 | 28 | ns | 0.977 |
| Fig.<br>Supp7<br>(lower) | DN | SHAM<br>PREV | W2-W3 | W4-W5 | 25.75 Hz | 8.78 Hz | 10 | 12 | *** | <0.001 |
|  |  |  | W2-W3 | W6-W7 | 25.75 Hz | 8.23 Hz | 10 | 11 | *** | <0.001 |
|  |  |  | W2-W3 | W8-W9 | 25.75 Hz | 12.22 Hz | 10 | 13 | *** | <0.001 |
|  |  |  | W4-W5 | W6-W7 | 8.78 Hz | 8.23 Hz | 12 | 11 | ns | 0.997 |
|  |  |  | W4-W5 | W8-W9 | 8.78 Hz | 12.22 Hz | 12 | 13 | ns | 0.570 |
|  |  |  | W6-W7 | W8-W9 | 8.23 Hz | 12.22 Hz | 11 | 13 | ns | 0.464 |
| Fig.<br>Supp7<br>(lower) | FN | SHAM | W2-W3 | W4-W5 | 15.59 Hz | 15.16 Hz | 7 | 8 | ns | 0.994 |
|  |  |  | W2-W3 | W8-W9 | 15.59 Hz | 10.37 Hz | 7 | 3 | ns | 0.623 |
|  |  |  | W4-W5 | W8-W9 | 15.16 Hz | 10.37 Hz | 8 | 3 | ns | 0.660 |
| Fig.<br>Supp7<br>(lower) | FN | SHAM<br>PREV | W2-W3 | W4-W5 | 26.22 Hz | 11.01 Hz | 18 | 14 | *** | <0.001 |
|  |  |  | W2-W3 | W6-W7 | 26.22 Hz | 7.32 Hz | 18 | 15 | *** | <0.001 |
|  |  |  | W2-W3 | W8-W9 | 26.22 Hz | 11.81 Hz | 18 | 16 | *** | <0.001 |
|  |  |  | W4-W5 | W6-W7 | 11.01 Hz | 7.32 Hz | 14 | 15 | ns | 0.714 |
|  |  |  | W4-W5 | W8-W9 | 11.01 Hz | 11.81 Hz | 14 | 16 | ns | 0.995 |
|  |  |  | W6-W7 | W8-W9 | 7.32 Hz | 11.81 Hz | 15 | 16 | ns | 0.545 |

**Table S10:**
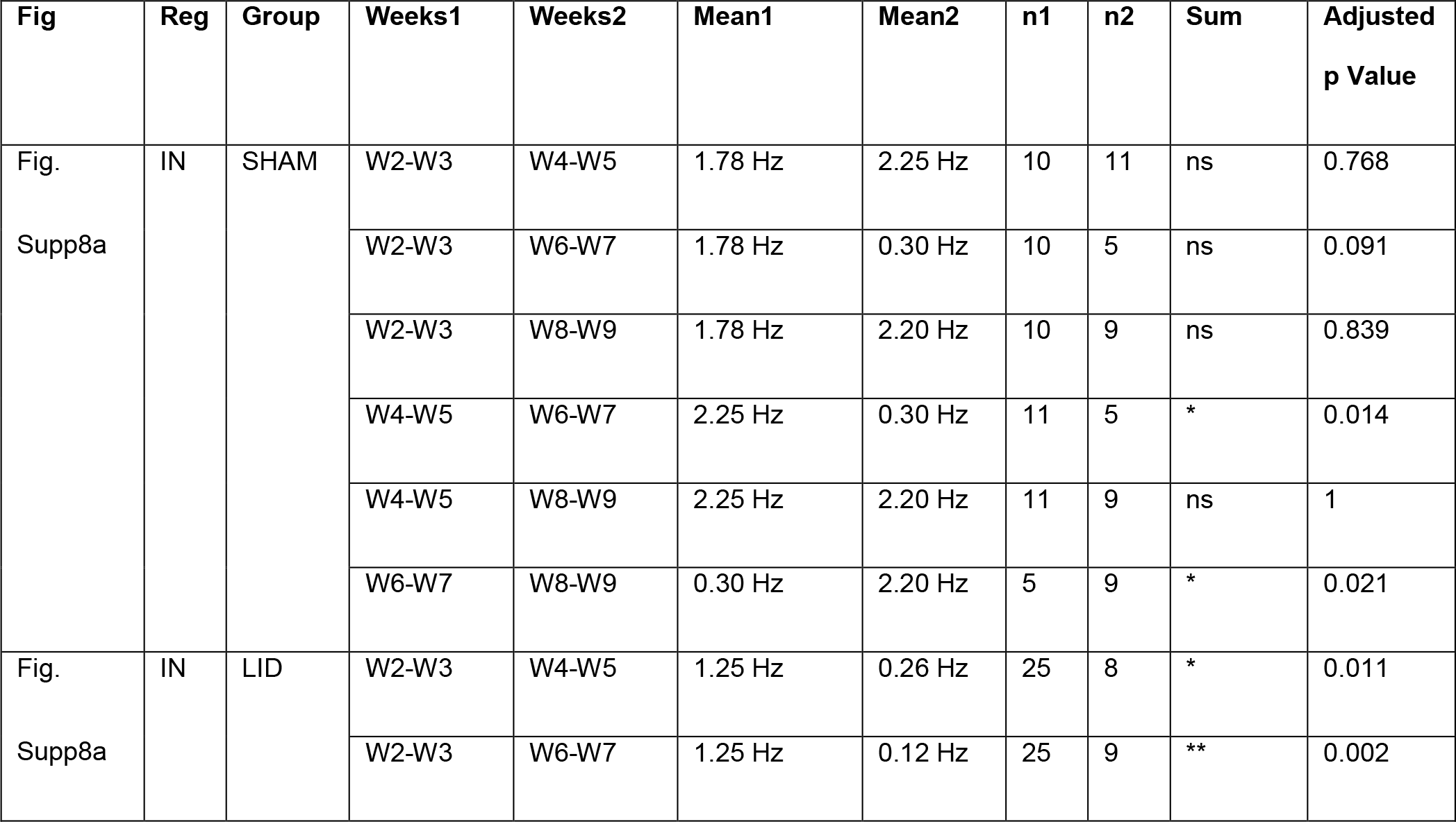
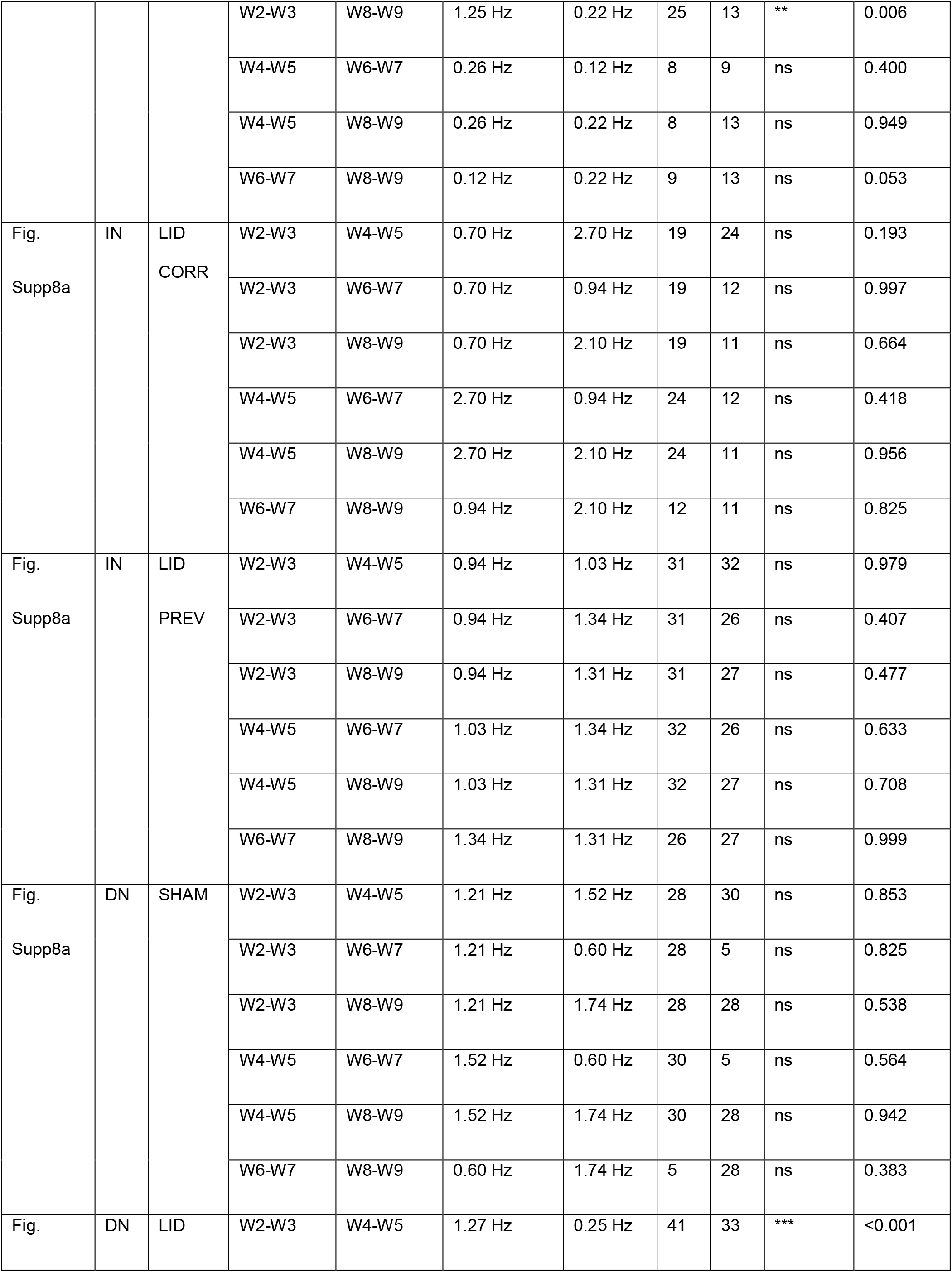
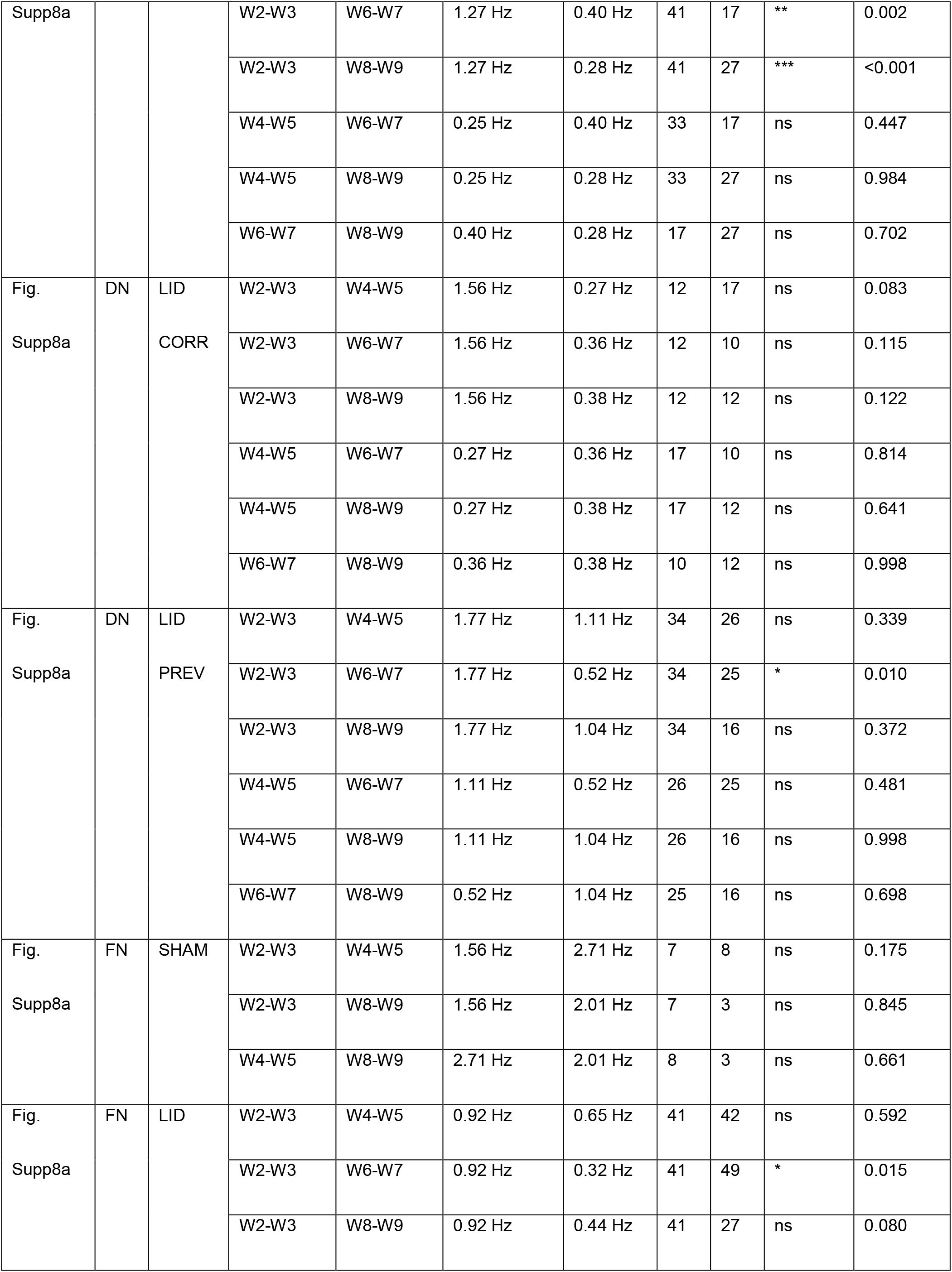
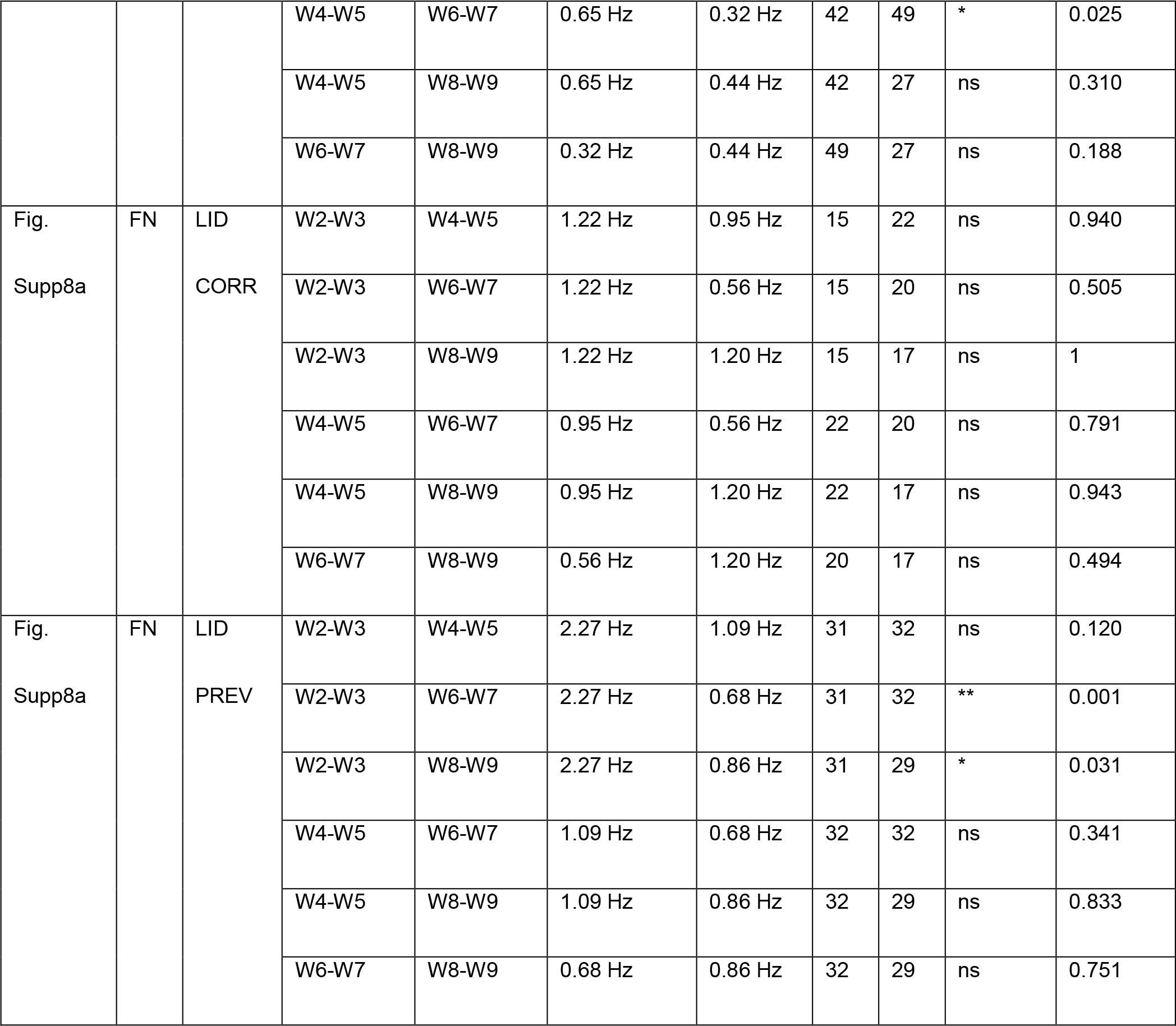
Precision measures, exact p-values, and replicate date relevant to Supplementary Figure8a.

**Table S11:**
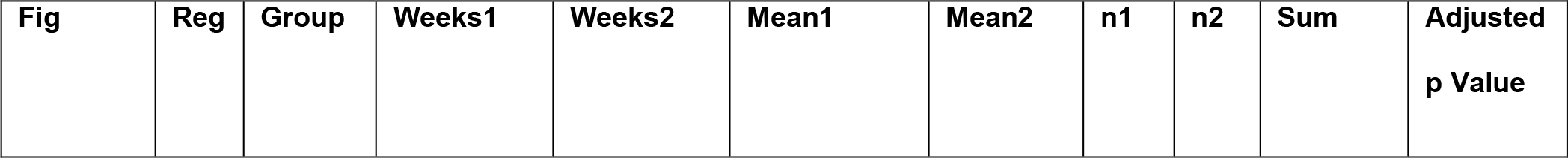
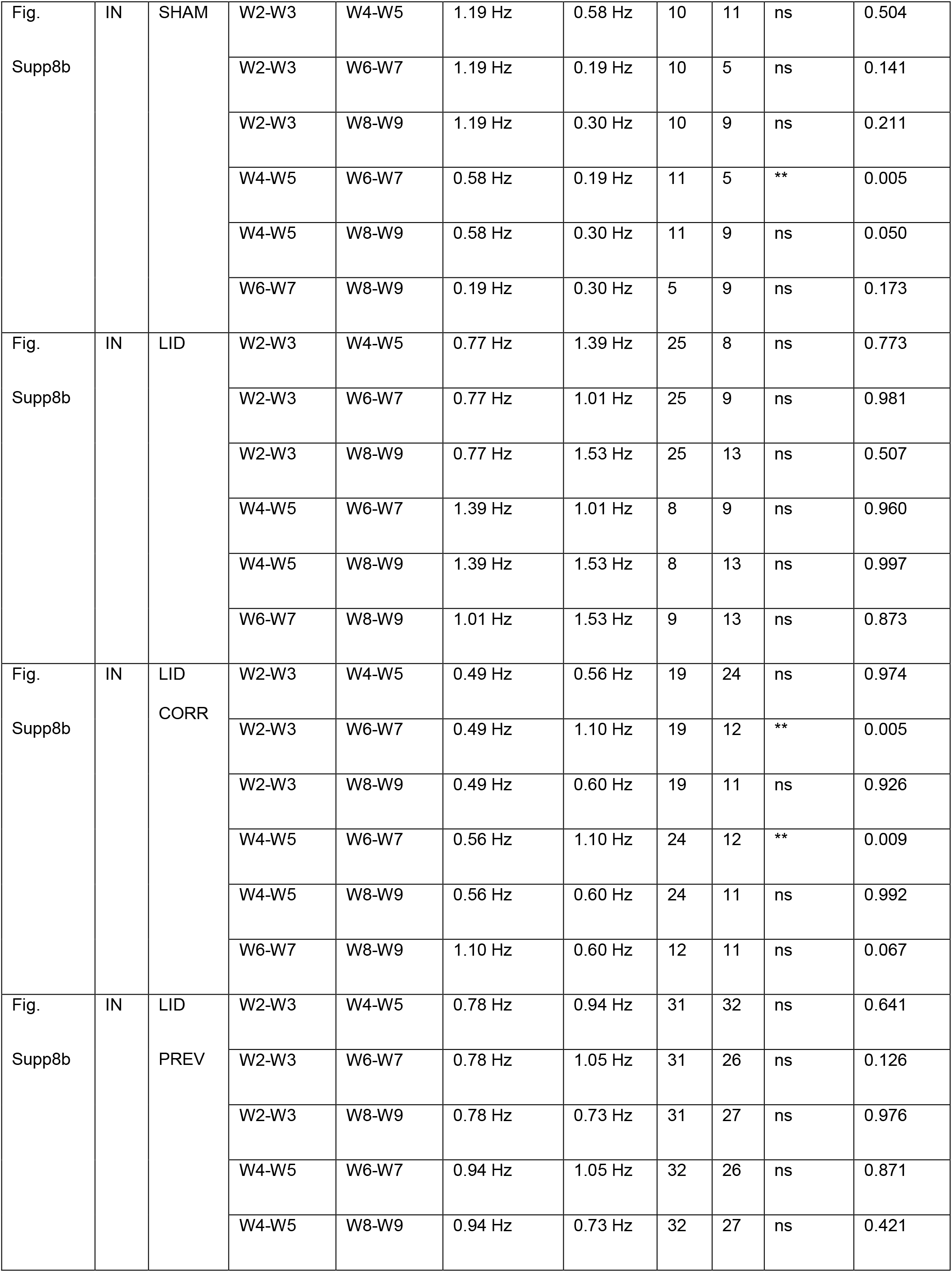
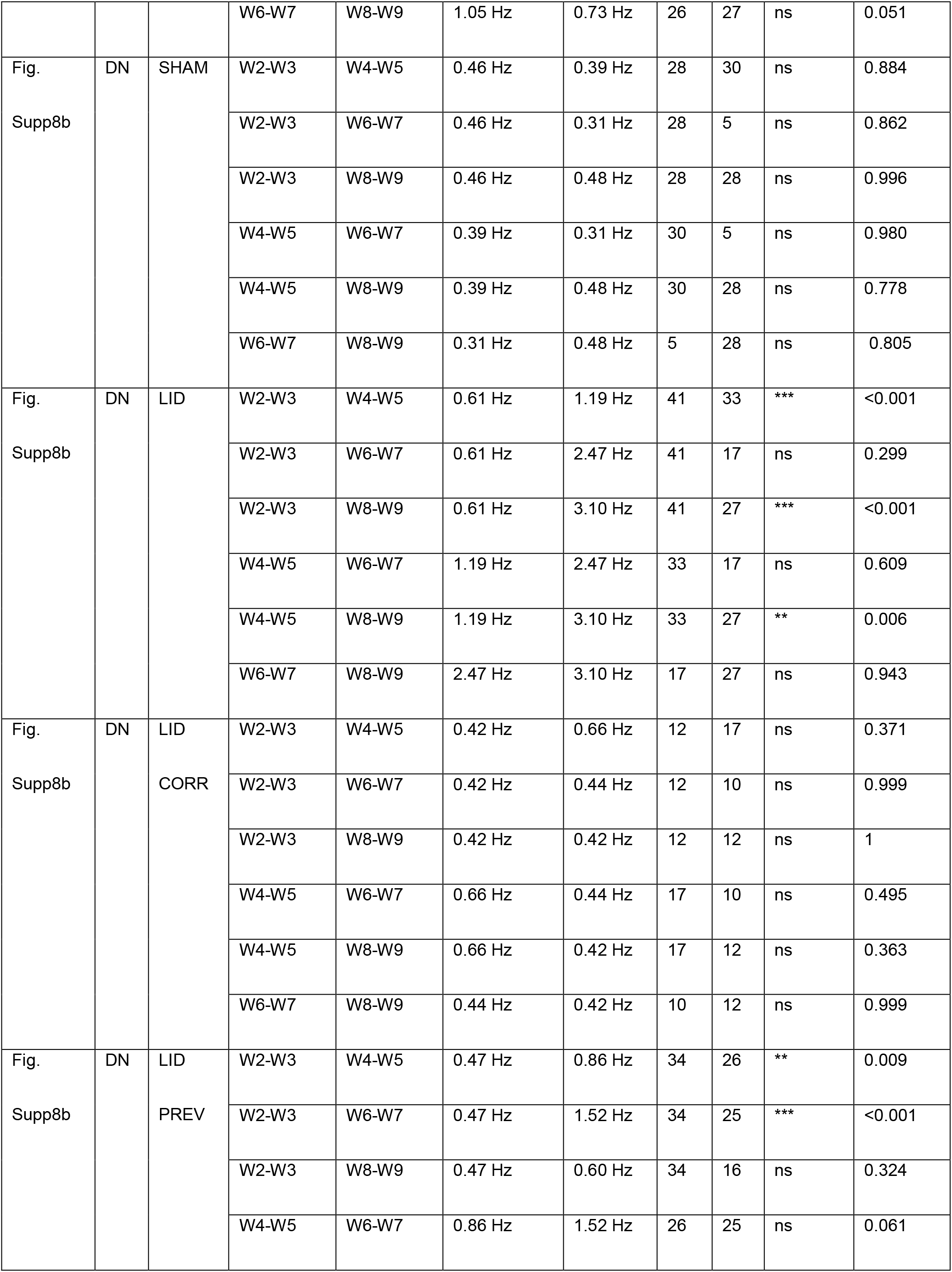
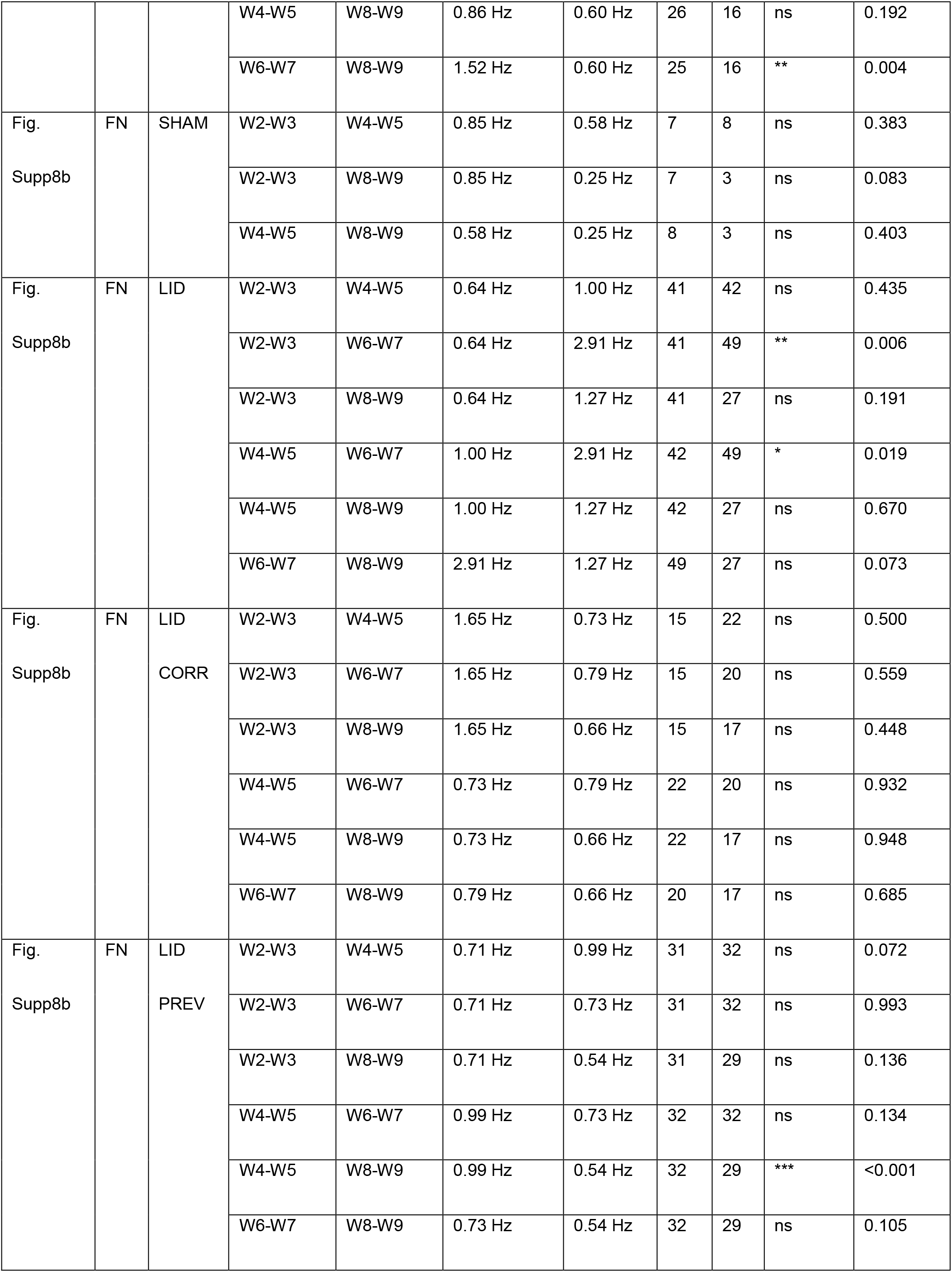
Precision measures, exact p-values, and replicate date relevant to Supplementary Figure8b.

**Table S12:** STDP in D2^-^ and D2^+^ MSN from SHAM, SHAM_PREV, LID and LID_PREV mice. Related to Figure 6.

| Group | D2<sup>-</sup> MSN | D2<sup>+</sup> MSN |
| --- | --- | --- |
| SHAM (N=5) | 149 ± 2, p<0.0001 (n=11) | 142 ± 4, p<0.0001 (n=10) |
| SHAM_PREV (N=5) | 123 ± 1, p<0.0001 (n=10) | 127 ± 1, p<0.0001 (n=10) |
| LID (N=7) | 172 ± 1, p<0.0001 (n=12) | 132 ± 3, p<0.0001 (n=9) |
| LID_PREV (N=5) | 65 ± 1, p<0.0001 (n=9) | 142 ± 6, p=0.0029 (n=10) |

